# A Retinoic Acid Autoregulatory Loop Governing Prefrontal-Motor Arealization

**DOI:** 10.64898/2025.12.23.696038

**Authors:** Lin Yang, Mikihito Shibata, Saejeong Park, Yuting Liu, Iva Salamon, Jia Liu, Suel-Kee Kim, Akemi Shibata, Ashley Deveau-French, Xoel Mato Blanco, Suxia Bai, Timothy Nottoli, Xiaojun Xing, Narjes Rohani, Stephan J. Sanders, Rothem Kovner, Kartik Pattabiraman, Nenad Sestan

## Abstract

The frontal lobe comprises the prefrontal association cortex (PFC), which supports complex cognition and goal-directed behavior, and the motor cortex (MC), which executes movement. A hallmark of primate brain evolution is PFC expansion accompanied by a posterior displacement of the MC. Retinoic acid (RA) signaling has emerged as a key regulator of PFC specification and expansion. However, the mechanisms that spatially confine RA signaling within the developing PFC, and the downstream RA-responsive gene networks, remain poorly understood. Here we defined an RA-associated gene regulatory network (RA-GRN) in the developing human PFC and identified *MEIS2*, which encodes a transcription factor linked to intellectual disability and autism spectrum disorder (ASD), as its key hub of this network. Conditional deletion of *Meis2* in postmitotic cortical excitatory neurons in mice results in a partial respecification of prospective prefrontal association territories toward motor-like molecular and connectional features, highlighting a critical role of postmitotic neurons in establishing and maintaining cortical areal identities. Concomitant with *Meis2* loss, the population of excitatory neurons expressing the RA-synthesizing enzyme ALDH1A3, and consequently RA signaling itself, is markedly reduced in the developing medial prefrontal cortex (mPFC). These findings revealed a conserved autoregulatory loop: RA → MEIS2 → ALDH1A3 → RA that reinforces a PFC-enriched RA gradient and organizes the MC–PFC axis. Together, our findings reveal a postmitotic mechanism by which specific features of neuronal identity reinforce RA signaling to define key features of prefrontal and motor cortical territories, linking a classic morphogen to transcriptional identity, neural circuit formation and function, and potentially to psychiatric disorders.

## Introduction

The frontal lobe is critically involved in executive processes, including decision-making, abstract cognition, planning, impulse regulation, motor control and personality^1–7^. In primates, the PFC’s disproportionate expansion relative to motor and sensory cortices, together with its protracted development, correlates with greater behavioral complexity and the emergence of higher-order cognitive processes^7–14^.

We previously found that broad gene expression gradients emanating from the fetal frontal and temporal lobes delineate prospective transmodal association cortices, including the PFC, whereas prospective primary and secondary sensorimotor areas exhibit more spatially restricted expression patterns^15–19^. Moreover, genes associated with RA signaling are embedded within the (pre)frontal gradient and are enriched in the developing medial frontal limbic cortex and PFC relative to the MC^15–18,20–23^, implicating RA in PFC development. RA, a metabolite of vitamin A, plays diverse and essential roles in neural development, including regional patterning, neuronal differentiation, and circuit formation^24–27^. In mice, alterations in RA signaling disrupt multiple aspects of medial frontal limbic and prefrontal gene expression and development^18,20–23^. In humans, genetic variants affecting components of the RA pathway have been linked to neurodevelopmental disorders and schizophrenia^24,28–30^. These observations led us to hypothesize that the mechanisms restricting and mediating RA signaling in the developing prefrontal cortex, and distinguishing it from the prospective motor cortex, could be elucidated by constructing an RA-GRN and functionally interrogating its key hub genes.

## Results

### Human mid-fetal prefrontal RA-GRN

We constructed a RA-GRN within the granular dorsolateral PFC (dlPFC), an anthropoid primate specialization implicated in multiple neurodevelopmental and psychiatric disorders^2–7,31–34^, during human mid-fetal development, a critical period for neuronal specification and circuit formation in the PFC^35^, using an integrative multi-omic approach. First, to map the global landscape of gene regulatory networks, we performed single-nucleus multiome (sn-multiome) profiling, generating paired snRNA-seq and snATAC-seq data from individual nuclei in four de-identified postmortem specimens at postconceptional weeks (PCW) 18–22 (**Methods; Supplementary Table 1**). We applied SCENIC+ to infer the dlPFC GRN based on core transcriptional regulators identified within each neuronal subtype (**Extended Data Fig. 1a-e; Supplementary Table 2)** and their top target genes (**Extended Data Fig. 1f**). Second, to identify a RA-regulated subnetwork, we generated genome-wide maps of binding sites for the RA receptors RARA, RARB, and RXRG, as well as active regulatory and transcriptional epigenetic markers (H3K4me3 and H3K27ac), in the human mid-fetal dlPFC (PCW 19-23; **Supplementary Table 1; Extended Data Fig. 2a-c**) using Cleavage under Target and Tagmentation (CUT&Tag)^36^. RAR/RXR binding sites were identified in proximal and distal regulatory regions (**Supplementary Table 3**) and were enriched for genes involved in axonogenesis, synaptogenesis, and forebrain development (**Extended Data Fig. 2d–g**). Finally, we integrated these two human datasets with human PFC-enriched genes identified using BrainSpan^37^ (**Extended Data Fig. 3; Supplementary Table 4**) to construct a putative RA-GRN in the human mid-fetal dlPFC (**Fig. 1a; Supplementary Table 5**).

**Figure. 1.**
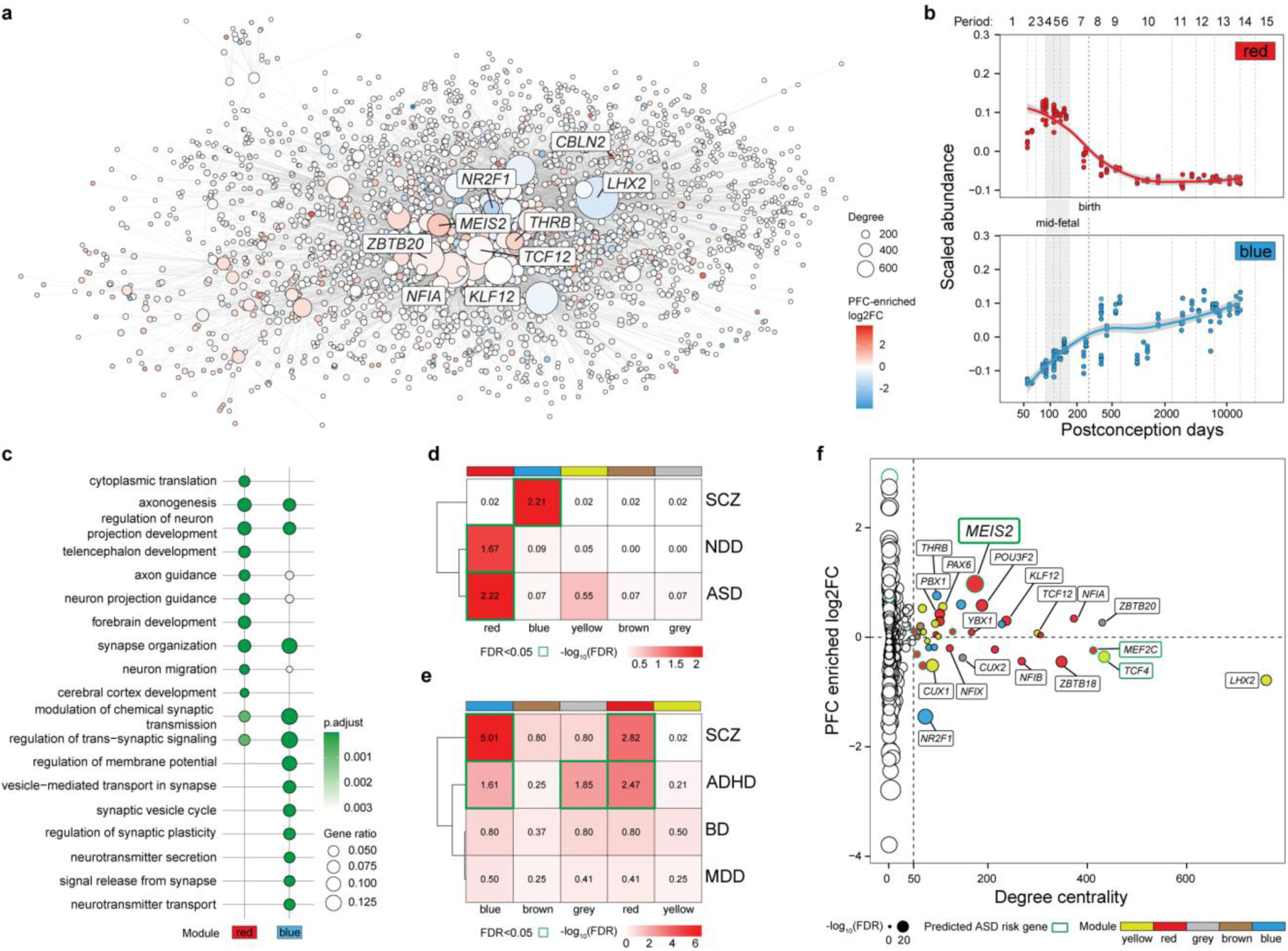
A retinoic acid–associated gene regulatory network in the human mid-fetal prefrontal cortex. **a,** Integrative strategy for constructing the RA-GRN in the human mid-fetal dlPFC, combining three datasets: (1) sn-multiome to infer GRN; (2) RAR/RXR CUT&Tag profiles to define RA-responsive target genes; and (3) BrainSpan to identify PFC-enriched genes. Visualization of constructed RA-GRN, with node size indicating degree centrality and node color reflecting PFC enrichment log₂ fold-change. **b,** Temporal expression modules of RA-GRN across prenatal and postnatal periods (using BrainSpan). Five modules were identified; two are shown here and three are shown in Extended Data Fig. 4a. Dashed lines indicate human development and adulthood periods. The red module peaks during the mid-fetal period, and the blue module peaks in the postnatal period. **c,** GO biological process enrichment of modules shown in b. The red module is associated with forebrain development and axon-related processes, and the blue module is associated with synapse-related processes. **d,** Association of RA-GRN modules with psychiatric disorders gene sets. The ASD gene set is from SFARI Gene (gene.sfari.org/); The NDD gene set is from the NIMH; The SCZ gene set is from a previous study^59^. FDR values are from gene set enrichment analysis; green borders indicate FDR < 0.05. **e,** Enrichment of common variants associated with other psychiatric disorders in RA-GRN modules. Numbers denote MAGMA linear-regression coefficients (β). FDR values are from MAGMA analyses of GWAS summary statistics; green borders indicate FDR < 0.05. **f,** RA-GRN analysis. Scatter plot of degree centrality (x-axis) versus PFC enrichment (log₂ fold-change; y-axis). White nodes indicate degree ≤ 50, whereas colored nodes (degree > 50) represent hub genes and correspond to module colors. Node size scales with −log₁₀(FDR) for PFC enrichment. Green node and label stroke marks the predicted ASD risk genes from the SFARI database. ASD, autism spectrum disorder; NDD, neurodevelopmental disorders; SCZ, schizophrenia; BD, bipolar disorder; ADHD, attention-deficit/hyperactivity disorder; MDD, major depressive disorder.

Weighted Gene Co-expression Network Analysis (WGCNA) of RA-GRN revealed five major RA-associated gene modules, underscoring the pleiotropic role of RA signaling in PFC development **(Fig. 1b,c; Extended Data Fig. 4a-c; Supplementary Table 6)**. Of these, the red module was highly expressed during mid-fetal development and enriched for genes involved in forebrain development and axon-related processes, whereas the blue module showed a progressive developmental increase and enrichment for synapse-related processes **(Fig. 1b,c)**. Integration with ASD, schizophrenia (SCZ), and neurodevelopmental disorder (NDD) risk gene databases revealed significant associations for genes within red and blue modules (**Fig. 1d; Extended Data Fig. 4d; Supplementary Table 7**). Both red and blue modules also showed significant enrichment for common variants associated with SCZ and attention-deficit/hyperactivity disorder (ADHD) in genome-wide association studies (GWAS) **(Fig. 1e; Extended Data Fig. 4e; Supplementary Table 8)**.

Network connectivity analysis of RA-GRN revealed multiple highly connected hubs with high degree centrality, each targeting more than 50 genes (**Fig. 1f**). Further characterization showed that these hub genes are direct RAR/RXR targets (**Extended Data Fig. 5a,b**), are enriched in specific cell subtypes (**Extended Data Fig. 5c,d**), and exhibit distinct spatiotemporal expression trajectories (**Extended Data Fig. 5e,f**). Among these hub genes, we identified *MEIS2* as a key hub within the red module, showing the strongest PFC enrichment and an association with ASD **(Fig. 1f)**, making it an especially compelling RA-regulated target. *MEIS2* encodes a highly conserved TALE-class homeobox transcription factor^38,39^ and was originally identified as an RA-regulated gene in the hindbrain and limbs^40,41^ (**Extended Data Fig. 6**). Heterozygous disruption of *MEIS2* is associated with both ASD and neurodevelopmental delay at genome-wide significance and is often accompanied by microcephaly. In addition, heterozygous loss of *MEIS2* is likely to underlie intellectual and motor disabilities observed with 15q14 deletions^42–47^.

### RA regulates *MEIS2* along the MC–PFC axis

Characterization of *MEIS2* spatiotemporal expression using human BrainSpan^37^ and spatial transcriptomic data^48^ showed its enrichment in multiple prospective areas of the human PFC relative to the MC during the mid-fetal period (**Extended Data Fig. 5e; Extended Data Fig. 7a-c**). Rostral enrichment of *Meis2* was also observed from postconceptional day (PCD) 13 to postnatal day (PD) 3 in mouse brain by whole-mount in situ hybridization (WISH) (**Extended Data Fig. 7d-f).** Complementary characterization of MEIS2 protein expression revealed a conserved MC–PFC gradient across species, including human PCW 18, macaque PCD 149, and mouse PD 7 (**Fig. 2a; Extended Data Fig. 7g-i; Extended Data Fig. 8j**). Longitudinal characterization of cell-type and laminar expression in mouse and human using sn-multiome and IHC showed *MEIS2* expression across neuronal lineages, with significant enrichment in SATB2^+^ excitatory upper-layer neurons^49^ in the mPFC (**Extended Data Fig. 8**). Given the similar expression profiles of *MEIS2* and RAR/RXR, we confirmed their colocalization in human and macaque cortical layers (**Extended Data Fig. 9**), as well as colocalization of *Meis2* and RA signaling in *RARE–lacZ* reporter mouse brains (**Extended Data Fig. 7; Extended Data Fig. 10a)**.

**Figure. 2.**
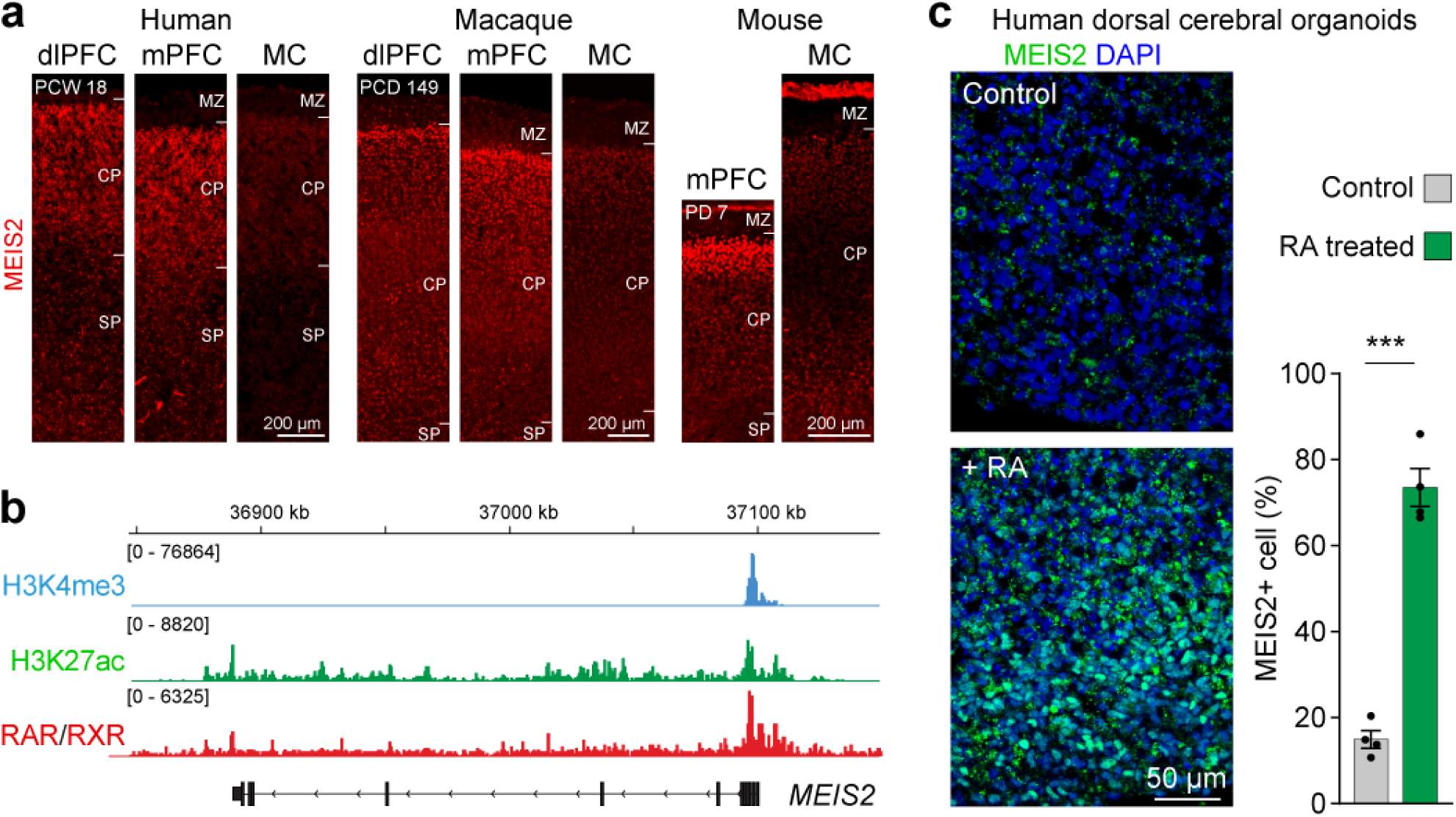
Retinoic acid regulates *MEIS2* expression along the developing motor–prefrontal cortical axis. **a,** IHC of MEIS2 in human PCW 18, macaque PCD 149 and mouse PD 7 cortex (dlPFC, mPFC and MC), highlighting a MC–PFC gradient with PFC enrichment. Quantitative analyses confirming this gradient are shown in Extended Data Fig. 7i and Extended Data Fig. 8j. MZ, Marginal zone; CP, cortical plate; SP, subplate. **b,** CUT&Tag profiles showing H3K4me3, H3K27ac and RAR/RXR binding at the *MEIS2* promoter in the human mid-fetal dlPFC. **c,** IHC of MEIS2 (green) and DAPI (blue) in 90-day human dorsal cerebral organoids. Compared with controls (top), 30-day RA treatment markedly increases the proportion of MEIS2⁺ cells (bottom). Scale bars, see images; statistical details, see Methods.

Supporting potential direct regulation of *MEIS2* by RA receptors, RAR/RXR CUT&Tag data showed direct binding to the human *MEIS2* promoter, and luciferase assays demonstrated synergistic transcriptional activation by the *Rarb*/*Rxrg* heterodimer via mouse *Meis2* promoter (**Fig. 2b; Extended Data Fig. 5a; Extended Data Fig. 10b**). Consistently, *Meis2* expression was markedly reduced in *Rarb/Rxrg* double-knockout brains (**Extended Data Fig. 10c**). Moreover, 30-day RA treatment of human cortical organoids robustly induced MEIS2 upregulation (**Fig. 2c).** Together, these findings demonstrate that RA signaling positively regulates *MEIS2* along the MC–PFC axis across species.

### Loss of *Meis2* disrupts arealization along the MC–PFC axis

To investigate the role of *Meis2* in PFC development, we generated a conditional *Meis2* mutant mouse line and deleted *Meis2* in postmitotic excitatory cortical neurons using *Nex1-Cre* (referred to as *Meis2* cKO) (**Extended Data Fig. 11).** Morphometric analysis revealed a significant reduction of frontal association areas in *Meis2* cKO mice **(Extended Data Fig. 12a,b)**. To identify potential patterning deficits in *Meis2* cKO mice, we examined the expression of developing PFC markers (*Cbln2 and Plxnc1*)^22^, MC markers (*Cyp26b1 and Etv5*)^18^, and primary somatosensory cortex (SSp) marker (*Bhlhe22*) (**Fig. 3a, b; Extended Data Fig. 12c-e**). *Cbln2 and Plxnc1* were nearly absent in *Meis2* cKO cortex (**Fig. 3a; Extended Data Fig. 12c**). Conversely, *Cyp26b1* and *Etv5* expanded medially from anterolateral motor cortex (ALM) and M2 region into the areas normally occupied by the mPFC (**Fig. 3b,c; Extended Data Fig. 12d,f**). We also observed anterior expansion of SSp markers RORB, BHLHE22, VGLUT2, and SEMA7A^19^ into the primary and secondary motor cortex (M1/MOp and M2/MOs; MOs/p) region (**Extended Data Fig. 12e; Extended Data Fig. 13; Extended Data Fig. 14**). Together, these findings support a model (**Fig. 3d**) in which *Meis2* deletion causes a partial loss of mPFC molecular identity, accompanied by an anterior shift of ALM and M2 region into the mPFC and a rostral expansion of M1 and somatosensory cortex (SS; SSs/p).

**Figure. 3.**
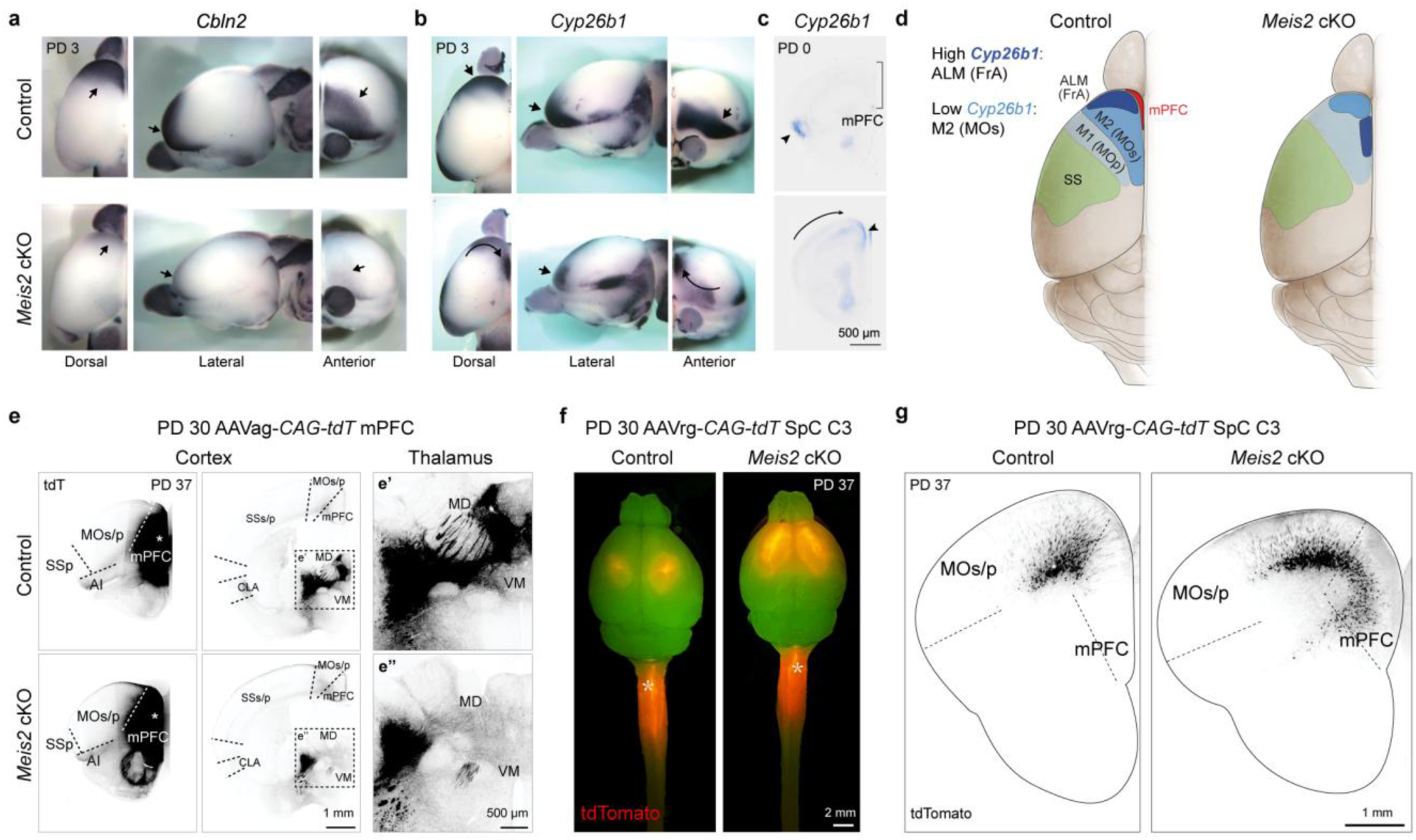
Loss of *Meis2* induces a shift in areal identity and connectivity in the frontal cortex. **a,b,** Expression of regional markers *Cbln2* (PFC and MC) and *Cyp26b1* (MC) in control and *Meis2* cKO brains. Arrows indicate *Cbln2* expression in the PFC and MC, and *Cyp26b1* expression in the MC. Curved arrows indicate ectopic medial expression of *Cyp26b1*. **c,** *Cyp26b1* expression in the rostral sections of control and *Meis2* cKO brain at PD 0. Arrows indicate *Cyp26b1* expression in the MC, and curved arrows indicate *Cyp26b1* ectopic medial expression. **d,** Schematic model of areal changes along the MC-PFC axis in *Meis2* cKO mice, illustrating dramatic shrinking or loss of the mPFC, a medial shift of the ALM (FrA) and M2, and rostral expansion of the M1 and SSp. **e,** Anterograde tracing of mPFC projection neurons using AAVag-*CAG–tdTomato* injected at PD 30. Enlarged views (e′–e′′) highlight tdTomato signal in the thalamic nuclei (MD, VM). Quantitative analyses confirming this observation are shown in Extended Data Fig. 15b. **f,g,** Retrograde tracing from the SpC C3. In *Meis2* cKO mice, tdTomato-labeled corticospinal-projecting neurons expand into the mPFC region. Quantitative analyses confirming this observation are shown in Extended Data Fig. 17e. Scale bars, see images; statistical details, see Methods.

We hypothesized that the altered areal molecular identities in *Meis2* cKO mice would also manifest as a reorganization of long-range connectivity. Reciprocal connectivity with medial thalamic nuclei, including the mediodorsal (MD) and ventromedial (VM) nuclei, and basolateral amygdala (BLA) is a defining connectivity signature of the mPFC^50^. In contrast, the defining efferent projection of the MC is the corticospinal tract (CST), and it receives afferent inputs from multiple sensorimotor areas^12^.

Characterization of PFC efferent projections using anterograde viral tracers at PD 30 revealed a significant reduction in projections to MD, VM, and BLA, whereas projections to the spinal cord (SpC) were increased in *Meis2* cKO mice compared to controls (**Fig. 3e; Extended Data Fig. 15a,b**). Similar decrease of the mPFC–MD connectivity and ectopic mPFC–SpC connectivity were observed in retrograde viral tracing from the MD and lipophilic DiI tracing (**Extended Data Fig. 15c; Extended Data Fig. 16**). Retrograde tracer injections into the C3 segment of the SpC (SpC C3) at PD 30 and PD120 confirmed a longitudinal expansion of CST-projecting L5 extratelencephalic (ET) neurons into the rostral and medial frontal cortex (**Fig. 3f,g; Extended Data Fig. 17**). These findings indicate a partial retention or acquisition of a key long-range CST connectivity feature that is characteristic of the mature (MC but not the PFC). During late fetal development in monkeys and the early postnatal period in rats and mice, CST projections are normally progressively refined and restricted to the motor and somatosensory cortices, and therefore do not originate from the mature PFC^51–53^.

To assess functional consequences of altered connectivity, we conducted a battery of behavioral assays. *Meis2* cKO mice showed impaired Y-maze performance with reduced spontaneous alternation **(Extended Data Fig. 18a)**, indicating working or spatial memory deficits. They also exhibited increased locomotor activity in the open-field test **(Extended Data Fig. 18b,c)**, as reflected by increased distance traveled, velocity, and movement time **(Extended Data Fig. 18d)**, while rotarod performance remained unchanged **(Extended Data Fig. 18e)**. In the nest-building assay, which reflects goal-directed spontaneous behavior, mutants showed impaired performance, consistently leaving nestlets largely intact **(Extended Data Fig. 18f,g)**. Together, these molecular and behavioral results demonstrate that conditional *Meis2* deletion in cortical excitatory neurons disrupts frontal lobe regional identities and impairs behaviors associated with prefrontal function.

### *Meis2* shapes the RA gradient along the MC–PFC axis across species

In addition to alterations in subcortical projections, rostral interhemispheric callosal projections were nearly absent in *Meis2* cKO mice, with labeling redirected to the ipsilateral cortex (**Fig. 4a; Extended Data Fig. 15b; Extended Data Fig. 19a,b**). Of note, among the 17 reported cases with heterozygous disruptive *MEIS2* variants and available clinical details, two describe a thin corpus callosum^42–45,47^. We identified a specific reduction in the rostral corpus callosum (CC) and anterior commissure (COA) in *Meis2* cKO mice, defined by L1CAM expression, whereas labeling in CST was increased (**Extended Data Fig. 19c,d**).

**Figure. 4.**
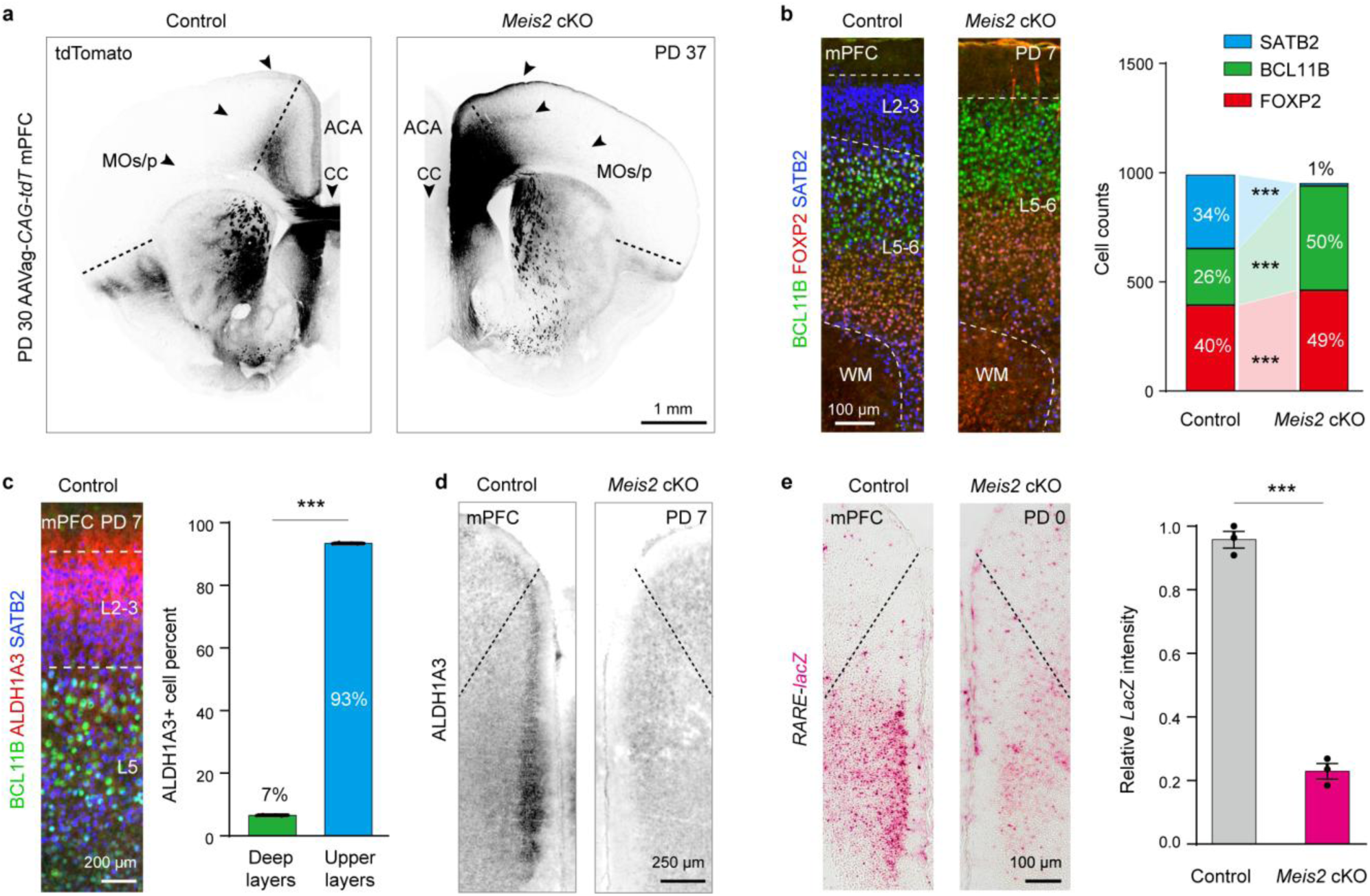
*Meis2* restricts retinoic acid signaling to the developing medial prefrontal cortex via ALDH1A3. **a,** Anterograde tracing of mPFC projection neurons using AAVag-*CAG–tdTomato* injected at PD 30 shows increased tdTomato-labeled projections from the mPFC to adjacent cortical areas (MOs/p) and reduced interhemispheric connectivity via the corpus callosum in *Meis2* cKO mice. Arrowheads indicate the axons projecting to the ipsilateral cortex. Quantitative analyses confirming these observations are shown in Extended Data Fig. 15b and Extended Data Fig. 19a,b. **b,** Laminar profiling of the mPFC in *Meis2* cKO mice using SATB2, BCL11B, and FOXP2. Loss of *Meis2* results in a near-complete depletion of SATB2⁺ upper-layer neurons and an increase in BCL11B⁺ layer 5 neurons. **c,** IHC of ALDH1A3 with SATB2 and BCL11B in the mPFC, showing that ALDH1A3⁺ cells are enriched in SATB2⁺ upper-layer projection neurons. **d,** IHC of ALDH1A3 in control and *Meis2* cKO mice. In *Meis2* cKO mice, ALDH1A3 expression is markedly reduced in the mPFC. Quantitative analyses confirming this reduction are shown in Extended Data Fig. 24f. **e,** Loss of RA signaling in the upper layers of the mPFC in *Meis2* cKO mice, as indicated by reduced β-galactosidase activity. Scale bars, see images; statistical details, see Methods.

As the CC and COA originate from upper-layer neurons and the CST from layer 5 neurons, we next assessed laminar organization in *Meis2* cKO mice and identified a pronounced reduction of SATB2⁺ upper-layer neurons^54^ accompanied by an expansion of BCL11B⁺ (also known as CTIP2, layer 5)^55^, and FOXP2⁺ (layer 6)^56^ neurons in the mPFC at PCD 16, PD 0, and PD 7 (**Fig. 4b; Extended Data Fig. 20**). EdU labeling of cortical neurons generated at PCD 16 revealed a marked reduction in the total number of EdU⁺ cells in *Meis2* cKO mPFC (**Extended Data Fig. 21a**). Further analysis showed a selective increase in EdU⁺BCL11B⁺ cells and a decrease in EdU⁺SATB2⁺ cells in the *Meis2* cKO mPFC (**Extended Data Fig. 21b**), suggesting a fate shift from upper- to deep-layer neurons.

To gain further insights into *Meis2* downstream genes relevant for PFC specification, we performed RNA-seq on the mPFC of control and *Meis2* cKO mice at PD 0 (**Extended Data Fig. 22a,b; Supplementary Table 9**). Among the differentially expressed genes (DEGs), those upregulated in the mutant were enriched for immune and apoptotic processes (**Extended Data Fig. 22c**). Consistent with this, increased numbers of cleaved CASPASE3^+^ cells and IBA1⁺ microglia were observed in the rostral cortex, along with modest cortical thinning along the rostral–caudal axis (**Extended Data Fig. 23**). In contrast, downregulated genes, including *Cbln2, Cdh8,* and *Syt4*, are involved in synapse-related functions (**Extended Data Fig. 22a,b,d**).

*Aldh1a3*, which encodes a key RA-synthesizing enzyme^57^, was significantly downregulated (**Extended Data Fig. 22a**). At PD 7, ALDH1A3 expression was specifically enriched in the SATB2⁺, or MEIS2⁺ upper-layer neurons of the mPFC (**Fig. 4c; Extended Data Fig. 24a-c**). Moreover, co-expression of ALDH1A3 and MEIS2 in the mPFC was well conserved in human, macaque, and mouse (**Extended Data Fig. 24d,e,g**). In *Meis2* cKO mouse brains, ALDH1A3 expression was nearly abolished at PD 7 (**Fig. 4d; Extended Data Fig. 24d,f**) with a concomitant reduction of RA signaling in the upper layers of the mPFC (**Fig. 4e)**. In human cortical organoids, 30-day RA treatment significantly increased both ALDH1A3^+^ cells and MEIS2^+^ALDH1A3^+^ cells (**Extended Data Fig. 24h,i**).

To better understand *MEIS2* gene regulatory networks, we examined MEIS2 chromatin occupancy in human PCW 21 dlPFC using CUT&Tag (**Supplementary Table 1; Extended Data Fig. 25a-c; Supplementary Table 10**). Functional enrichment analysis of MEIS2 target genes highlighted axonogenesis, synapse organization, and forebrain development (**Extended Data Fig. 25d,e; Extended Data Fig. 26**), closely paralleling biological processes associated with RAR/RXR and suggesting coordinated roles in PFC network regulation (**Extended Data Fig. 2e,g**; **Extended Data Fig. 25f,g**). Strong MEIS2 binding was observed at the *SATB2 and BCL11B* loci, but not at *ALDH1A3* (**Extended Data Fig. 27a; Extended Data Fig. 26a**), suggesting that ALDH1A3 downregulation likely results from indirect regulation through upper-layer specialization in the mPFC (**Fig. 4b-d)**. Moreover, RA-treated human cortical organoids exhibited an increase in ALDH1A3⁺ neurons (**Extended Data Fig. 24j,k**). Together, these findings demonstrate that *MEIS2* reinforces RA signaling within PFC species by promoting the production of *ALDH1A3*⁺ neurons in the mPFC.

## Discussion

In this study, the human mid-fetal PFC RA-GRN we reconstructed reveals mechanisms underlying morphogen self-stabilization and the establishment of the MC–PFC axis. The RA-GRN comprises genes strongly implicated in fundamental neurodevelopmental processes and are enriched for genes associated with ASD, SCZ, and ADHD, underscoring its value in identifying additional risk genes and the underlying mechanisms^24,28–30^.

Within this network, we identify the high-confidence ASD risk gene *MEIS2* as a central hub. Multi-scale analyses uncover an intrinsic RA autoregulatory loop: RA → MEIS2 → ALDH1A3 → RA that stabilizes the PFC-high to MC-low RA gradient (**Extended Data Fig. 28**). RA activates RAR/RXR signaling to sustain *MEIS2* expression, and MEIS2 promotes production of *ALDH1A3*⁺ neurons by regulating the *SATB2*–*BCL11B* balance in the mPFC. Disruption of this loop in postmitotic neurons of *Meis2* cKO mice leads to a collapse of the RA gradient within the cortical plate and destabilization of the MC–PFC axis, highlighting the importance of postmitotic neurons in establishing and maintaining key molecular and connectional features of cortical areal identity. *MEIS2* also governs neuronal fate and survival in the striatum^39,58^, another RA-enriched region, suggesting that similar mechanisms may operate in the ventral forebrain.

We further propose that *MEIS2* acts as a key hub within the developmental program of (para)limbic and transmodal association cortices, consistent with the MIND model^19^, which posits competing transcriptomic identity programs governing primary sensorimotor versus association cortex formation. *Meis2* and *Plxnc1* are broadly expressed in these association territories, with peak expression in the PFC (**Extended Data Fig. 7d**; **Extended Data Fig. 12c**). In line with this framework, *Meis2* deletion in our study contracts transmodal association cortices, including the PFC, temporal association cortex, and retrosplenial cortex marked by *Plxnc1*, and leads to a relative expansion of motor and sensory identities (**Extended Data Fig. 12c–e; Extended Data Fig. 14**). Together with prior work, our findings support a unified model explaining how molecular mechanisms balance the patterning of association versus motor-sensory cortices.

## Acknowledgements

The authors thank Alvaro Duque for providing macaque tissues and NeuroInfo software, and Shaojie Ma, Anadita Nadkarni, and Sirisha Pochareddy for help with data generation or analyses. This work was also supported by NIH grants NS095654, MH106934, MH116488, MH110926, MH129981, MH129751, and MH122681 as well as by the Simons Foundation (SFARI #736613).

## Contributions

L.Y., M.S., and N.S. conceived and designed the research. M.S., S.B., and T.N. and X.X. designed and generated the *Meis2 fl*/*fl* mouse. I.S., J.L., A.D., and R.K. processed human tissue samples and prepared the sn-multiome libraries. M.S. and K.P. processed mouse tissue and prepared the sn-multiome libraries. S.P. and Y.L. analyzed the mouse and human sn-multiome data, respectively. S.-K.K. carried out organoid experiments, including microscopy. A.S. carried out ISH staining and 3D reconstructions. M.S. bred mice, and performed ISH, WISH, and DiI tracing experiments. L.Y. bred mice, performed IHC staining, viral axonal tracing, and CUT&Tag experiments, analyzed the data, generated RNA-seq libraries, conducted behavioral tests, performed bioinformatics analyses (including RA-network, RNA-seq, sn-multiome, GWAS and MAGMA analyses), prepared all figures, and wrote the first draft of the manuscript. S.J.S., K.P. and N.S. provided funding for the study. N.S. supervised data quality control, data analysis, and manuscript preparation, and edited the final version of the manuscript. All authors read and approved the final version of the manuscript.

---

**Extended Data Figure. 1.**
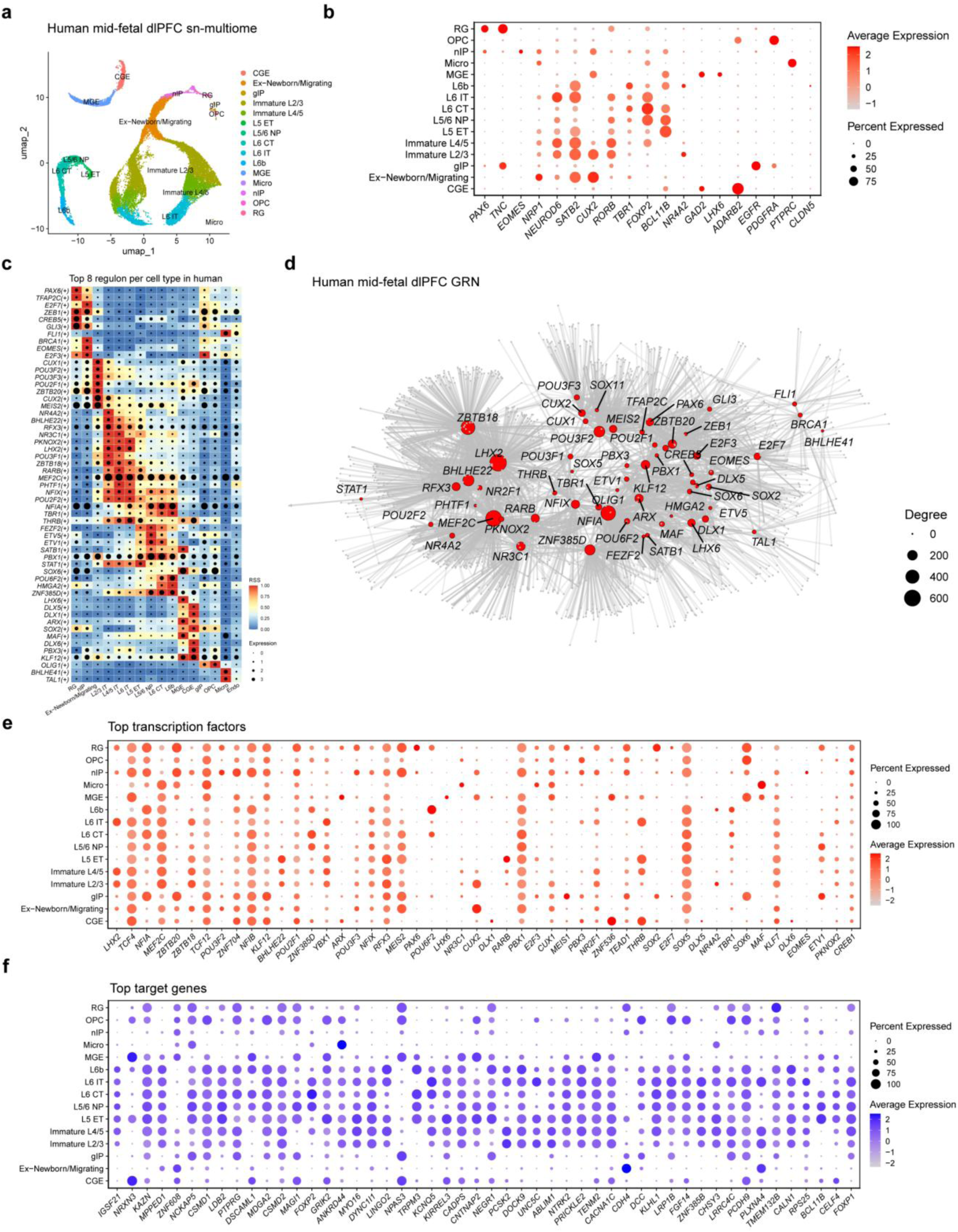
GRN of the human mid-fetal dlPFC. **a,** UMAP visualization of sn-multiome data, annotated by major cortical cell types. **b,** Cell-type classification based on canonical marker-gene expression. **c,** Heatmaps showing transcription-factor regulon activity (regulon specificity scores, RSS) and corresponding gene expression across cortical cell types; each row represents a regulon and each column a cell type. **d,** Graph representation of the dlPFC GRN. Nodes represent transcription factors (TFs) and target genes, and edges denote predicted regulatory interactions inferred by SCENIC+. Node size reflects degree centrality. **e,** Cell-type-resolved expression patterns of top TFs within the dlPFC GRN. **f,** Cell-type-resolved expression patterns of top target genes within the dlPFC GRN.

**Extended Data Figure. 2.**
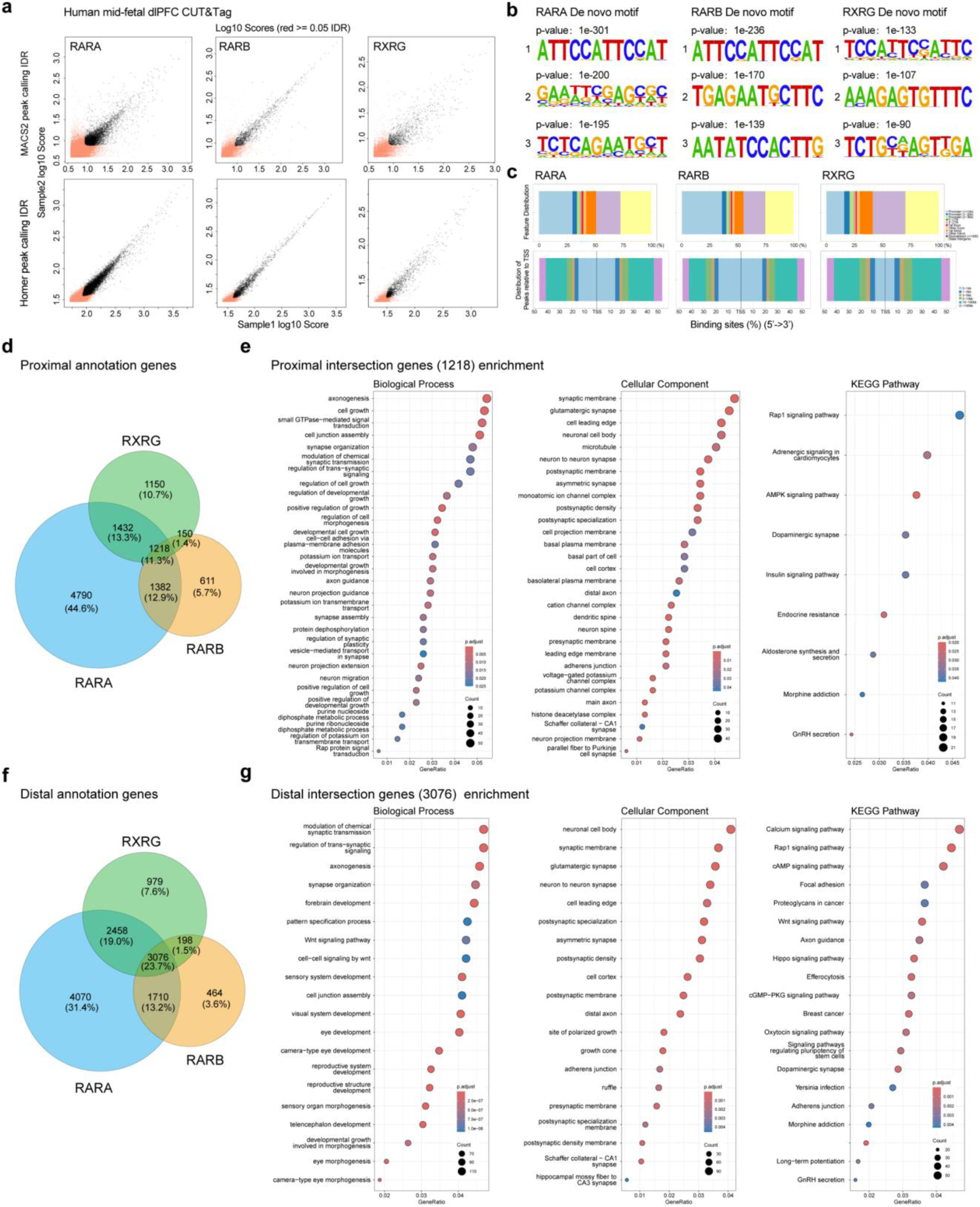
Genome-wide profiling of RAR/RXR binding in the human mid-fetal dlPFC. **a,** Reproducibility of CUT&Tag replicates for RAR/RXR, with peaks called by MACS2 (top) and HOMER (bottom) and evaluated by IDR (IDR < 0.05, black). **b,** Top three de novo motifs enriched in RAR/RXR peaks. **c,** Genomic annotation of RAR/RXR peaks across promoters, UTRs, exons, introns and distal intergenic regions, and distribution relative to transcription start sites. **d,** Overlap of proximal peak–gene annotations for RAR/RXR using ChIPseeker. **e,** GO biological process, cellular-component and KEGG pathway enrichment of 1,217 shared proximal targets. Dot size indicates gene count; color denotes adjusted *P* value. **f,** Overlap of distal peak–gene annotations for RAR/RXR identified by rGREAT. **g,** GO biological process, cellular-component and KEGG pathway enrichment of 3,076 shared distal targets.

**Extended Data Figure. 3.**
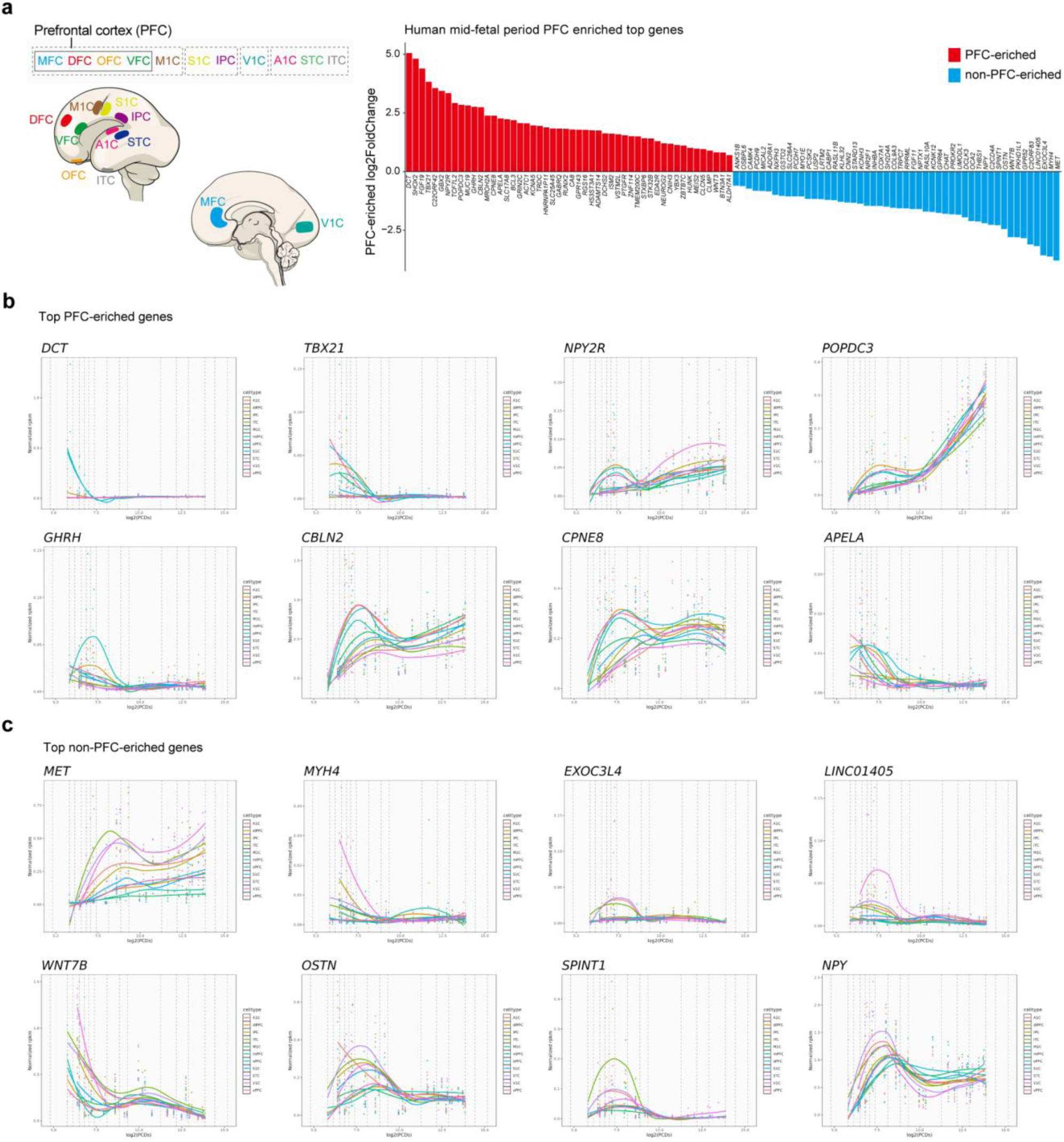
Identification of human mid-fetal PFC–enriched genes. **a,** Cross-regional transcriptomic comparison using BrainSpan dataset. Bar plot showing genes enriched in the PFC during the mid-fetal period, ranked by log₂ fold-change (PFC versus other cortical areas; red, enriched; blue, non-enriched). **b,** Spatiotemporal expression profiles of representative PFC-enriched genes. **c,** Spatiotemporal expression profiles of representative non–PFC-enriched genes. The dashed lines indicate the periods of human development and adulthood.

**Extended Data Figure. 4.**
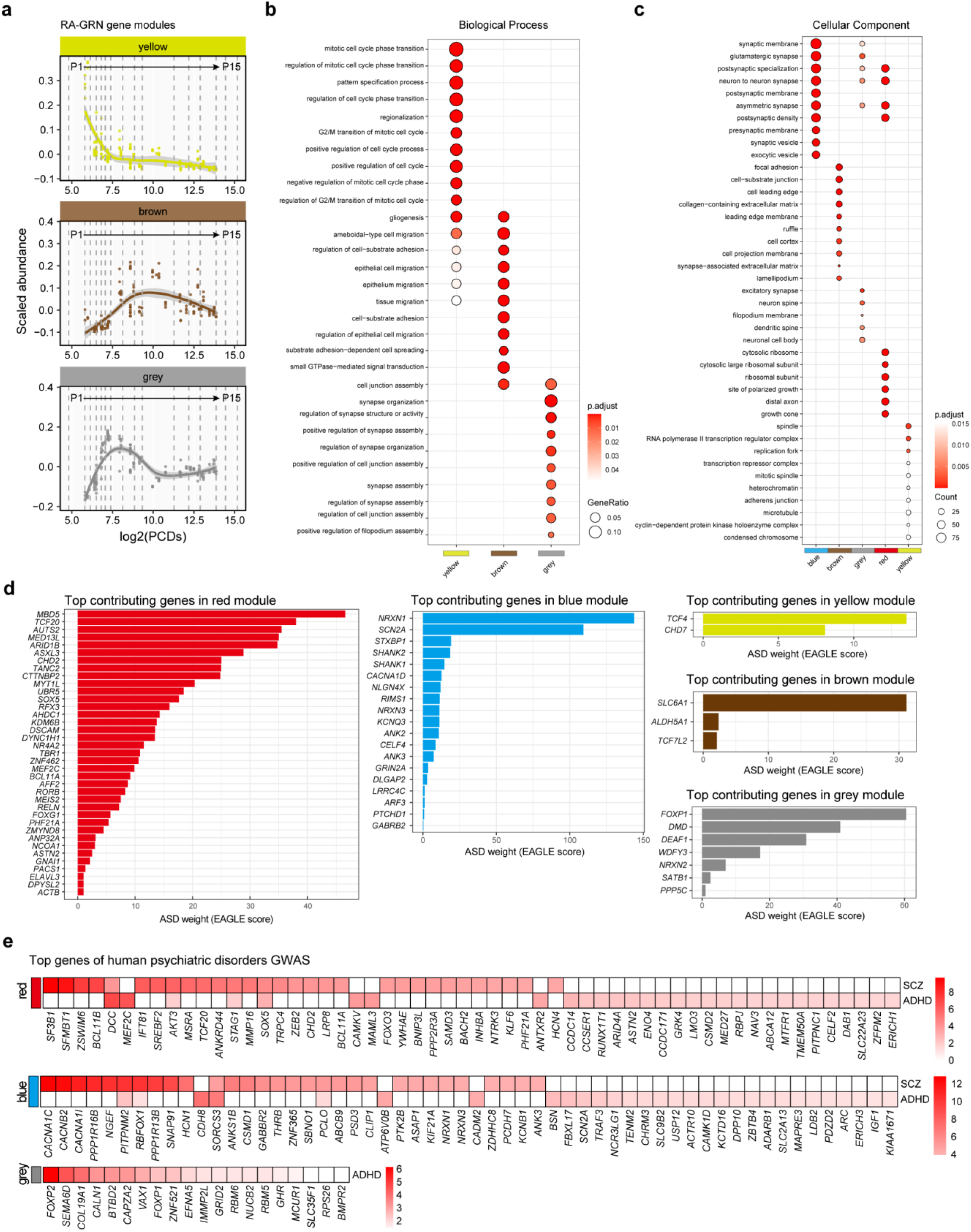
RA-GRN temporal gene modules associations with function and human psychiatric disorder. **a,** RA-GRN temporal gene modules across prenatal development (using BrainSpan dataset). Five modules were identified and three of them are shown here. Another two modules are shown in Fig. 1b. The dashed lines indicate the periods of human development and adulthood. **b,** GO biological process enrichment of the modules in a, as shown in Fig. 1c. The yellow module (rapidly declining) is enriched for replication-fork activity; the brown module (transiently increasing) for neuronal migration; and the grey module (increasing then plateauing) for synapse assembly. **c,** GO cellular component enrichment of the modules in a and Fig.1b. The yellow module (rapidly declining) is enriched for cell-division processes; the red module (slowly declining) for distal axons; the brown module (transiently increasing) for cell junctions; the grey module (increasing then stabilizing) for dendritic spines; and the blue module (continuously increasing) for synaptic structures. **d,** Top contributing genes (EAGLE score, from SFARI database) within RA-GRN temporal modules associated with ASD. Each bar plot shows genes contributing to ASD enrichment in the indicated module. **e,** Top genes enriched in modules showing significant associations across psychiatric disorders GWAS. Heatmaps indicate gene-level significance across disorders. ASD, autism spectrum disorder; SCZ, schizophrenia; ADHD, attention-deficit/hyperactivity disorder.

**Extended Data Figure. 5.**
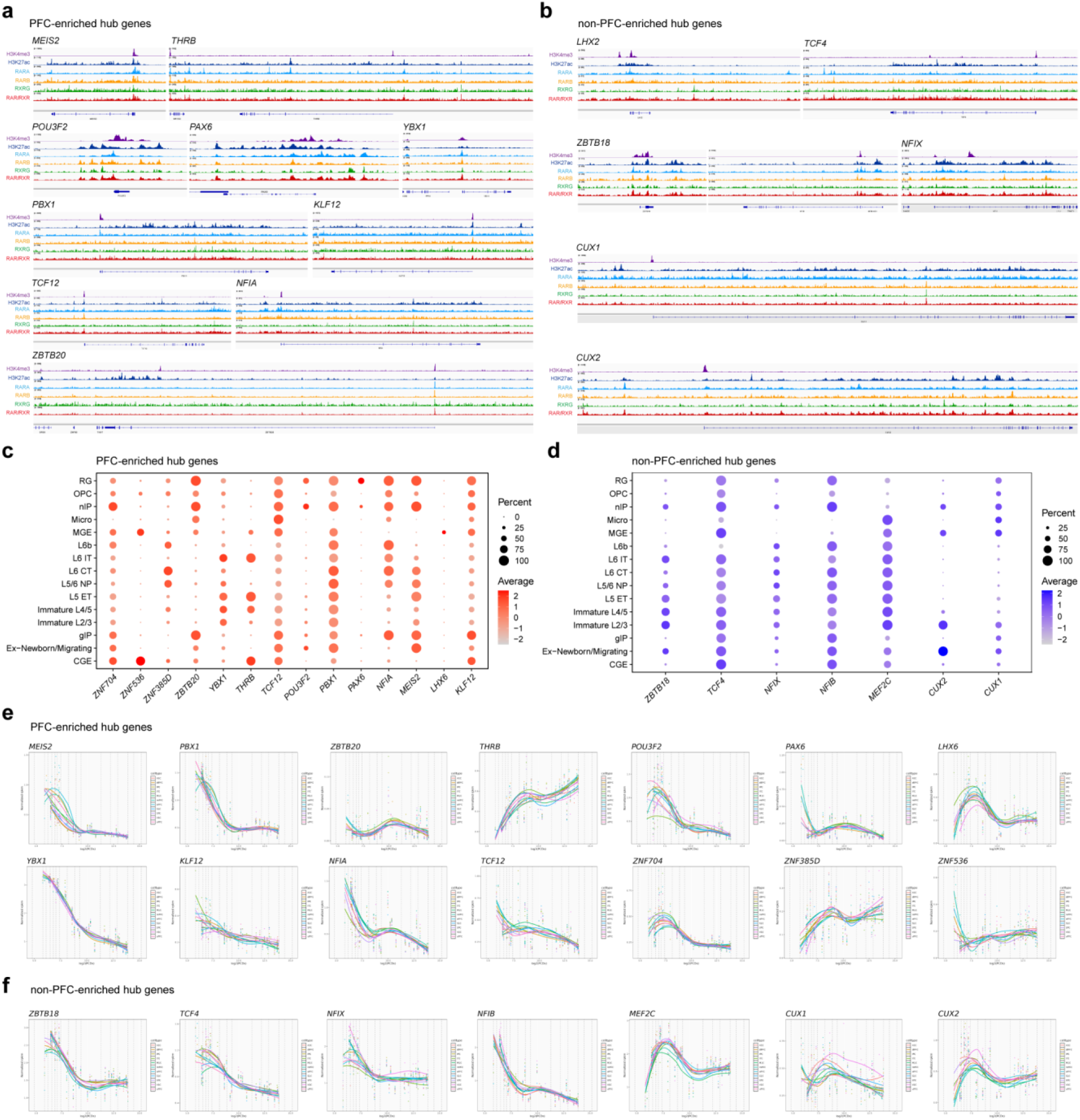
Characterization of RA–GRN hub genes. **a, b,** CUT&Tag tracks for H3K4me3, H3K27ac, and RAR/RXR binding sites in the human mid-fetal dlPFC. Genome-browser views illustrate representative RA-GRN hub genes that are PFC-enriched or non–PFC-enriched. **c,** Dot plot of RA-GRN hub genes which are PFC-enriched across major cortical cell types in the human mid-fetal dlPFC. Dot size indicates the proportion of expressing cells, and the color scale denotes average expression level. **d,** Dot plot of RA-GRN hub genes which are non–PFC-enriched across major cortical cell types in the human mid-fetal dlPFC. **e,** Spatiotemporal expression profiles of RA-GRN hub genes which are PFC-enriched across human brain regions and developmental time points. Normalized expression values are plotted against post-conceptional days, with fitted curves indicating developmental trajectories. **f,** Spatiotemporal expression profiles of RA-GRN hub genes which are non–PFC-enriched across human brain regions and developmental time points.

**Extended Data Figure. 6.**
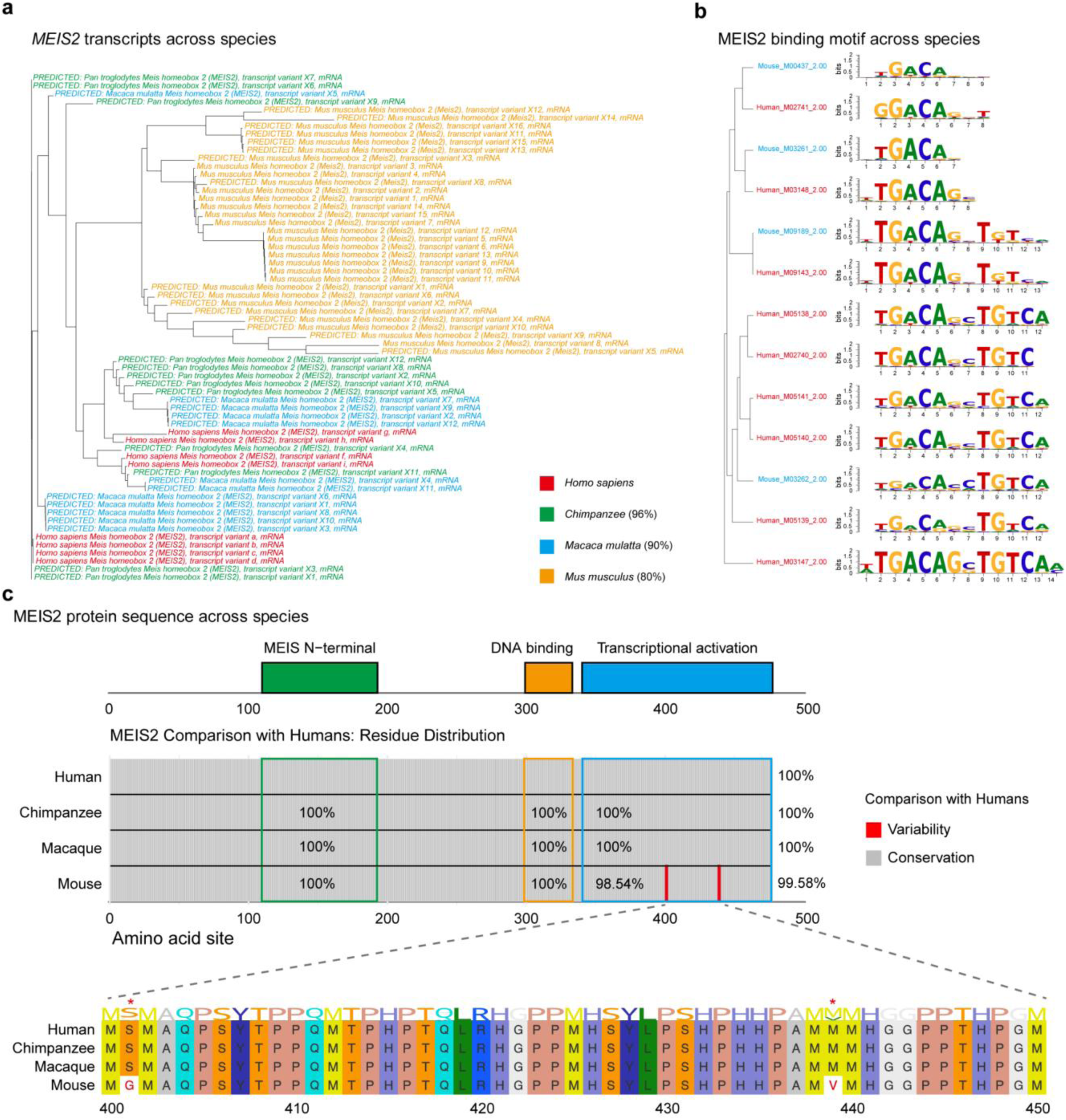
Evolutionary conservation of *MEIS2* transcript isoforms, DNA-binding motifs and protein domains across species. **a,** Phylogenetic tree of *MEIS2* transcript isoforms in human, chimpanzee, macaque and mouse based on nucleotide sequence alignment. Transcript diversity and species-specific clustering are indicated by color. **b,** Sequence logos of MEIS2 DNA-binding motifs across species derived from position weight matrix alignment, showing conserved core motif structure with subtle divergence in flanking bases in mouse and macaque relative to human. **c,** Protein-level conservation of MEIS2 across species. Top, domain architecture showing N-terminal, DNA-binding and transcriptional-activation domains. Middle, amino-acid conservation relative to human across domains, with near-complete identity in primates and slightly reduced conservation in mouse, particularly in the C-terminal region. Bottom, alignment of amino acid residues 400–450 highlighting species-specific substitutions within the transcriptional activation domain.

**Extended Data Figure. 7.**
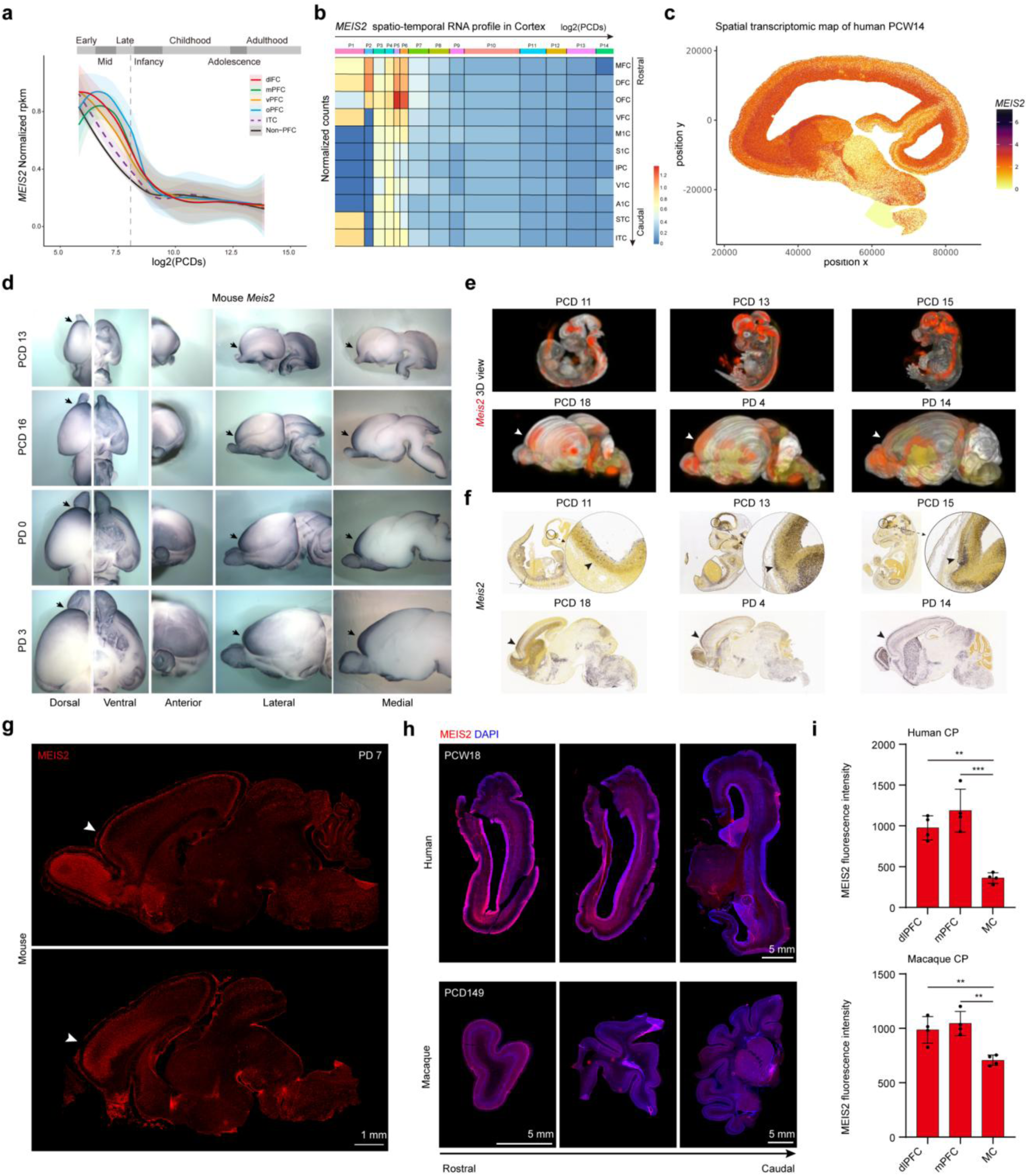
Spatiotemporal dynamics of *MEIS2* expression across species. **a,** Spatiotemporal expression of *MEIS2* in the human cortex from BrainSpan dataset. Expression peaks during the mid-fetal period across prefrontal subregions (dlPFC, mPFC, vPFC, oPFC), inferotemporal cortex (ITC) and non-prefrontal areas, with consistently higher levels in the PFC. The dashed lines indicate the periods of human development and adulthood. **b,** Expression trajectories of *MEIS2* across cortical regions from early post-conception stages to infancy, showing progressive enrichment in the PFC. **c,** Spatial transcriptomic map of *MEIS2* expression in a mid-sagittal section of the human brain at PCW 14. Each dot represents an individual spot, colored by normalized expression (yellow, low; purple, high). **d,** Mouse *Meis2* expression detected by WISH in the developing brain at PCD 13, PCD 16, PD 0, and PD 3. Rostral expression is indicated by arrows. **e,** 3D images of whole-mount mouse embryos and brains from PCD 11 to PD 14 stained for *Meis2* RNA (from Allen Brain Atlas), showing dynamic and regionally enriched forebrain expression (white arrowheads). **f,** ISH of *Meis2* at matched developmental stages from PCD 11 to PD 14 (from Allen Brain Atlas), confirming progressive enrichment in the dorsal telencephalon. Insets highlight rostral forebrain localization (black arrowheads). **g,** IHC of MEIS2 protein in sagittal mouse brain sections at PD 7, showing rostral enrichment in the cortex (white arrowheads). **h,** IHC of MEIS2 protein in the human brain at PCW 18 and in the macaque brain at PCD 149, revealing a rostro–caudal gradient with higher expression in rostral (PFC) than caudal areas (MC). **i,** Quantification of MEIS2 fluorescence intensity in cortical plate (CP) across three cortical regions (dlPFC, mPFC and MC) in the human and macaque, corresponding to Fig. 2a. Scale bars, see images; statistical details, see Methods.

**Extended Data Figure. 8.**
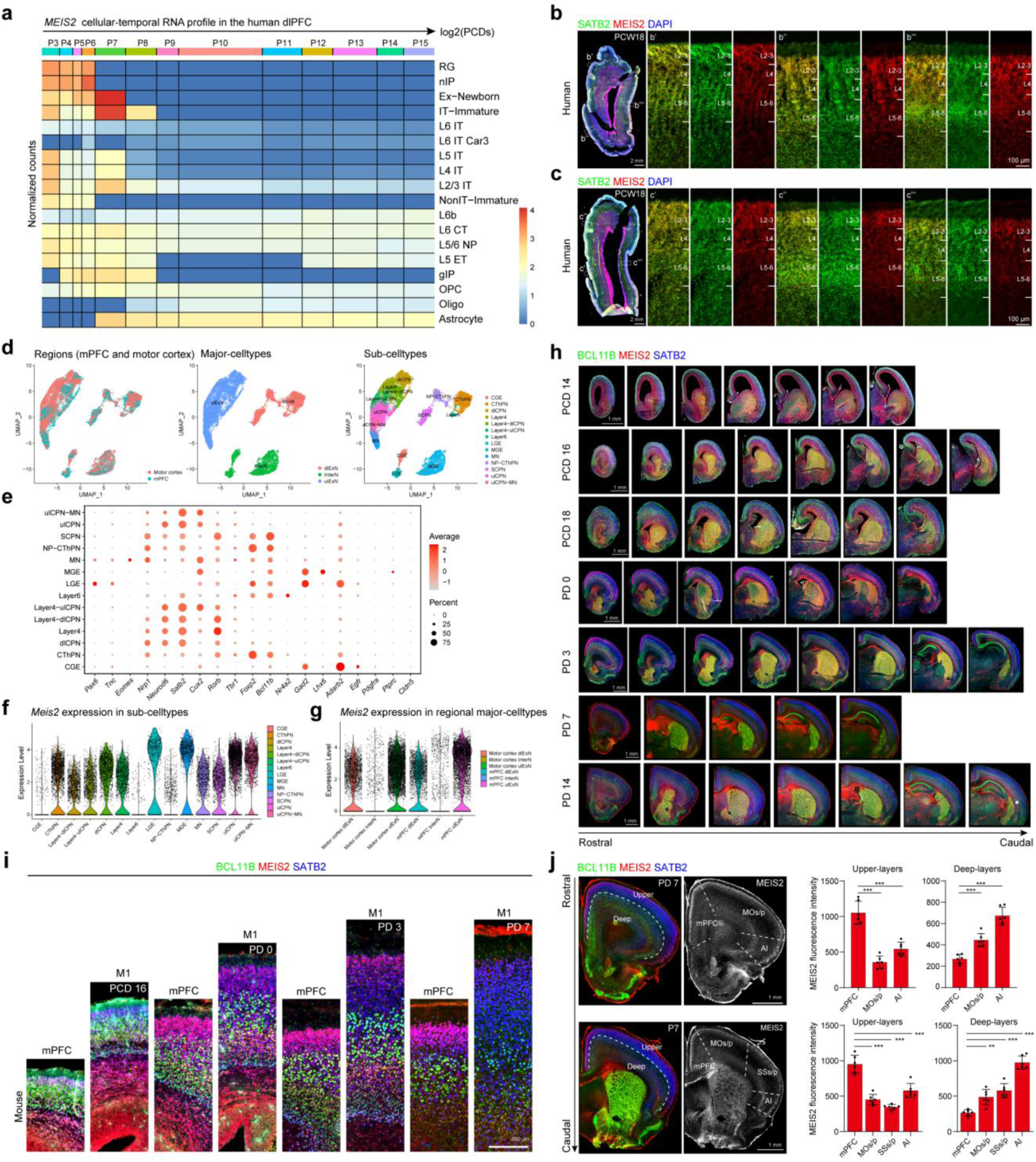
Cellular dynamics of *MEIS2* expression across species. **a,** Cell-type–resolved expression of *MEIS2* in the developing dlPFC (unpublished data), showing stage-specific enrichment in excitatory neuronal lineages. **b, c,** IHC of human brain sections at PCW 18 stained for SATB2 (green), MEIS2 (red) and DAPI (blue). Boxed regions are enlarged in a’-a’’’, and b’-b’’’. **d,** UMAP visualization of sn-multiome data from the mouse P0 mPFC and motor cortex, colored by region (left), major cell types (middle) and sub-cell types (right). **e,** Dot plot of representative marker gene expression across identified sub-cell types. Dot size indicates the proportion of expressing cells, and color reflects average expression levels. **f,** Violin plots of *Meis2* expression across sub-cell types. **g,** Violin plots showing regional *Meis2* expression in major cell types of the mPFC and motor cortex. **h,** IHC of coronal mouse brain sections from PCD 14 to PD 14 stained for MEIS2 (red), BCL11B (green) and SATB2 (blue), showing spatiotemporal dynamics of MEIS2 protein expression along the caudal–rostral axis. Strong colocalization with SATB2⁺ upper-layer neurons is observed from PCD 16 onwards, particularly in rostral cortical regions of the mPFC. **i,** IHC of coronal brain sections from PCD 16 to PD 7 stained for MEIS2 (red), BCL11B (green) and SATB2 (blue), showing spatiotemporal dynamics of MEIS2 protein expression along the MC–PFC axis. Strong colocalization with SATB2⁺ upper-layer neuron is observed from PCD 16 onwards, particularly in the mPFC. **j,** Coronal section of the PD 7 mouse cortex, highlighting selective MEIS2 protein expression in upper layers of the mPFC. Quantification of MEIS2 fluorescence intensity across four cortical regions (mPFC, MOs/p, SSs/p and agranular insular (AI)) in upper (left) and deep (right) layers shows MEIS2 significant enrichment in upper-layer neurons of the mPFC. Scale bars, see images; statistical details, see Methods.

**Extended Data Figure. 9.**
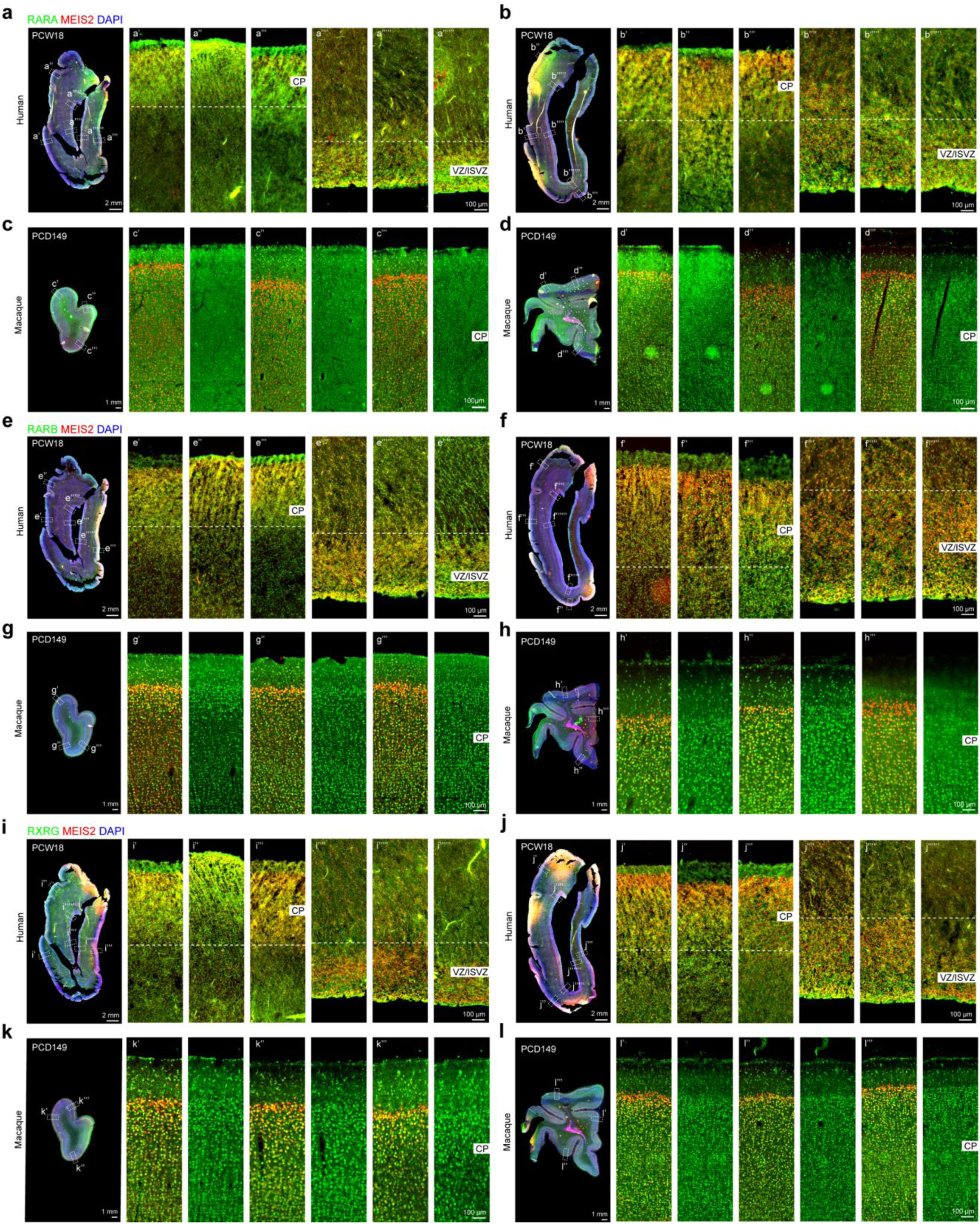
Colocalization of RAR/RXR and MEIS2 across species. **a, b,** IHC of RARA (green), MEIS2 (red) and DAPI (blue) in coronal sections of human brain at PCW 18, showing co-expression in both the cortical plate (CP) and ventricular/inner subventricular zones (VZ/ISVZ). Enlarged views (a′–a′′′′′′, b′–b′′′′′′) highlight RARA–MEIS2 colocalization across CP–VZ/ISVZ. **c, d,** IHC of RARA (green), MEIS2 (red) and DAPI (blue) in coronal sections of macaque brain at PCD 149, showing similar co-expression patterns in the CP. Enlarged views (c′–c′′′′′′, d′–d′′′′′′) highlight RARA–MEIS2 colocalization within the CP. **e, f,** IHC of RARB (green), MEIS2 (red) and DAPI (blue) in coronal sections of human brain at PCW 18, showing co-expression in both the CP and VZ/ISVZ. Enlarged views (e′–e′′′′′′, f′–f′′′′′′) highlight RARB–MEIS2 colocalization across CP–VZ/ISVZ. **g, h,** IHC of RARB (green), MEIS2 (red) and DAPI (blue) in coronal sections of macaque brain at PCD 149, showing similar co-expression patterns in the CP. Enlarged views (g′–g′′′′′′, h′–h′′′′′′) highlight RARB–MEIS2 colocalization within the CP. **i, j,** IHC of RXRG (green), MEIS2 (red) and DAPI (blue) in coronal sections of human brain at PCW 18, showing co-expression in both the CP and VZ/ISVZ. Enlarged views (i′–i′′′′′′, j′–j′′′′′′) highlight RXRG–MEIS2 colocalization across CP–VZ/ISVZ. **k, l,** IHC of RXRG (green), MEIS2 (red) and DAPI (blue) in coronal sections of macaque brain at PCD 149, showing similar co-expression patterns in the CP. Enlarged views (k′–k′′′′′′, l′–l′′′′′′) highlight RXRG–MEIS2 colocalization within the CP. Scale bars, see images.

**Extended Data Figure. 10.**
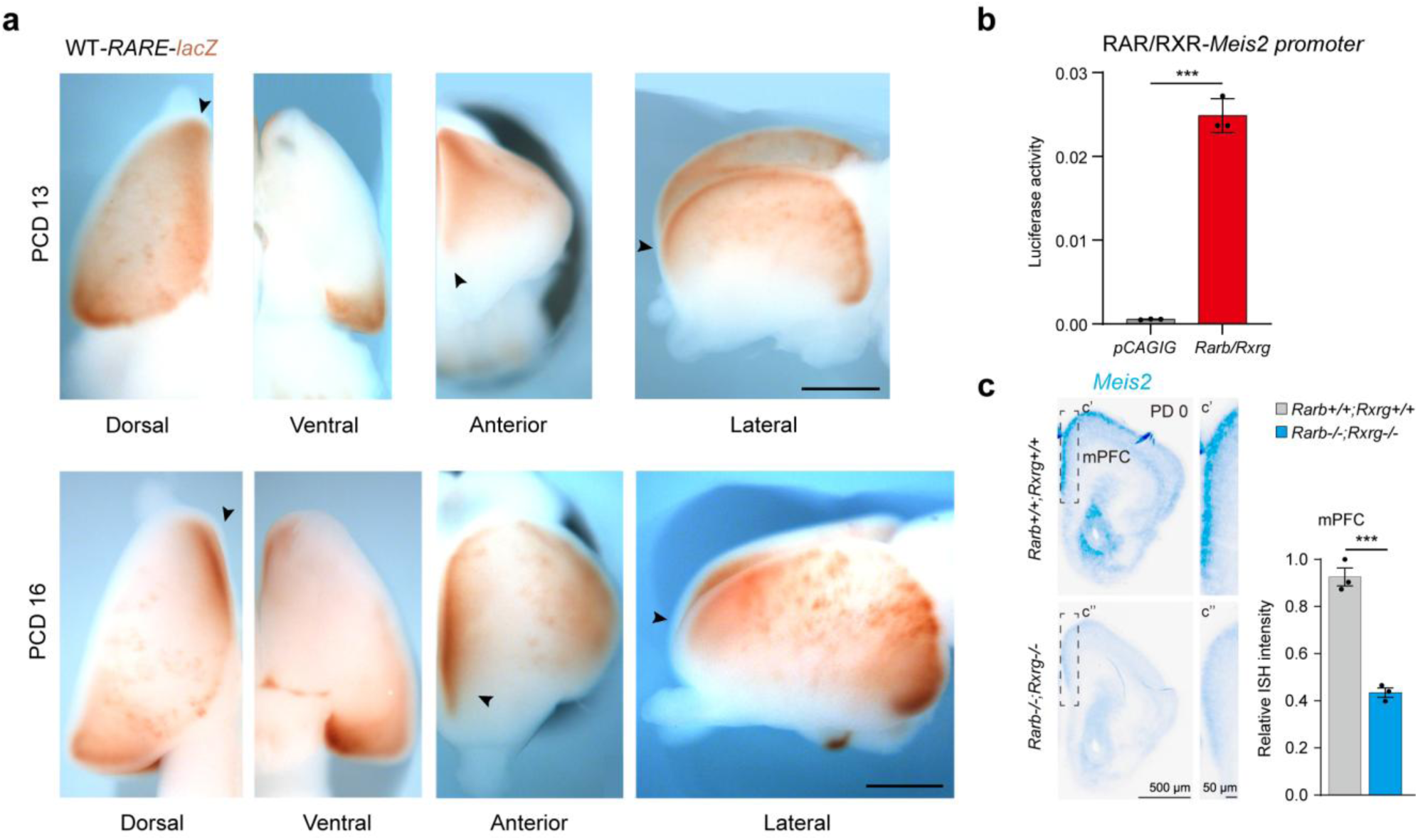
RA signaling regulates *Meis2* expression in the mouse mPFC. **a,** Whole-mount b-Galactosidase staining of the *RARE*–*lacZ* mouse brains at PCD 13 and 16. **b,** *Meis2* promoter luciferase assay in Neuro2a cells with mouse *Rxrg* and *Rarb*. **c,** *Meis2* expression in PD 0 *Rarb/Rxrg* double-knockout (dKO) and control mice, showing reduced *Meis2* expression in the mPFC. Enlarged views (c′–c′′) highlight *Meis2* ISH signal in the mPFC. Scale bars, see images; statistical details, see Methods.

**Extended Data Figure. 11.**
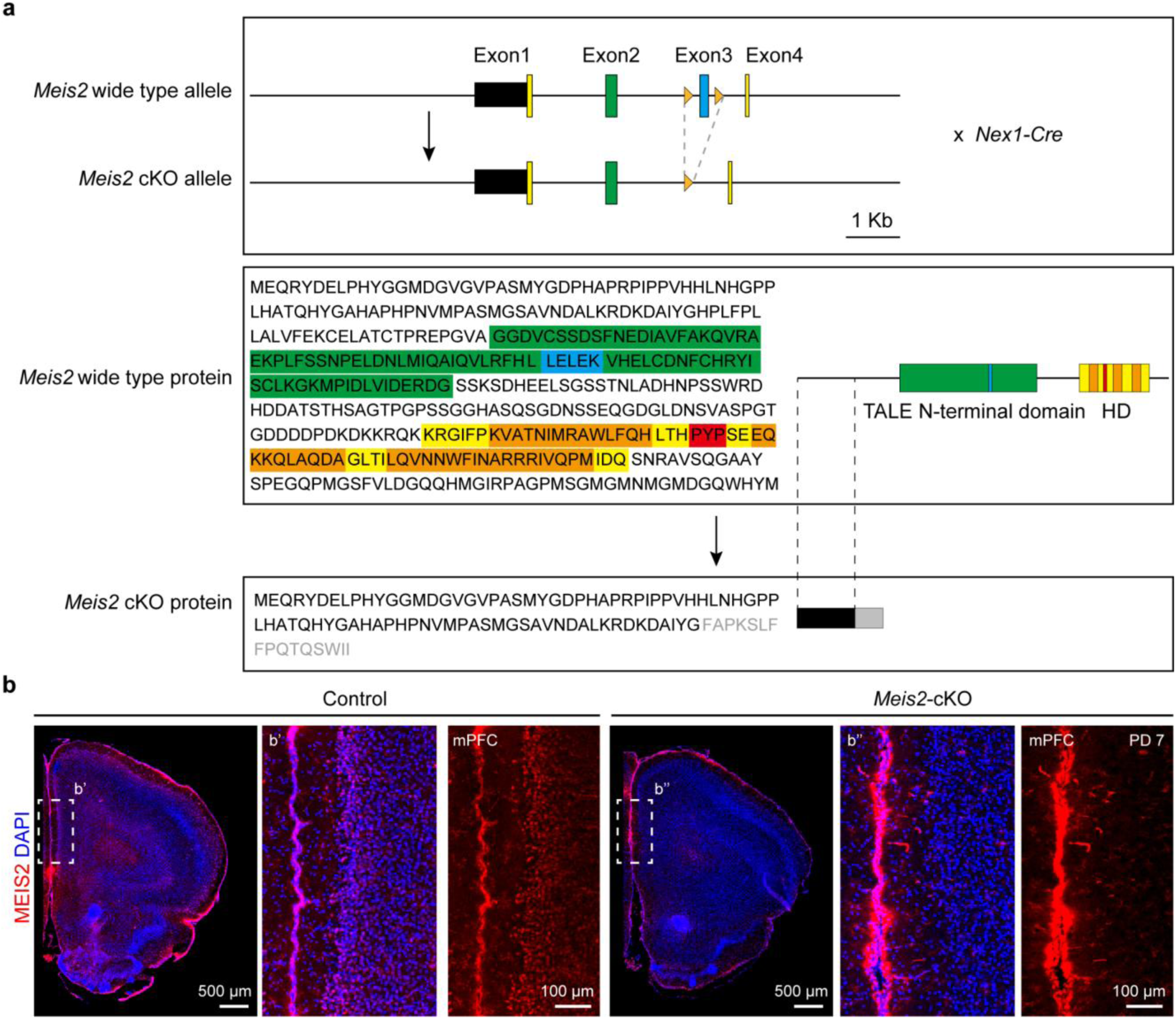
Generation and validation of *Meis2* conditional knockout mice. **a,** Schematic of the *Meis2* wild-type and cKO alleles. LoxP sites (orange triangles) were inserted flanking exon 3 (blue), resulting in exon excision upon Cre recombination. Protein sequences of *Meis2* wild-type and *Meis2* cKO alleles are shown. In the WT protein, conserved motifs within the TALE N-terminal domain (green) and homeodomain (HD; yellow) are preserved. Key residues, including the nuclear export signal (blue), nuclear localization signal (red), and helix loops (orange), are highlighted. The truncated cKO protein lacks these functional domains, with premature termination indicated in grey. Scale bar, 1 kb. **b,** IHC of MEIS2 (red) and DAPI (blue) in PD 7 brain sections from control and *Meis2* cKO mice. Enlarged views (**b′–b′′**) highlight the mPFC region, where MEIS2 signal is markedly reduced in *Meis2* cKO mice. Scale bars, see images.

**Extended Data Figure. 12.**
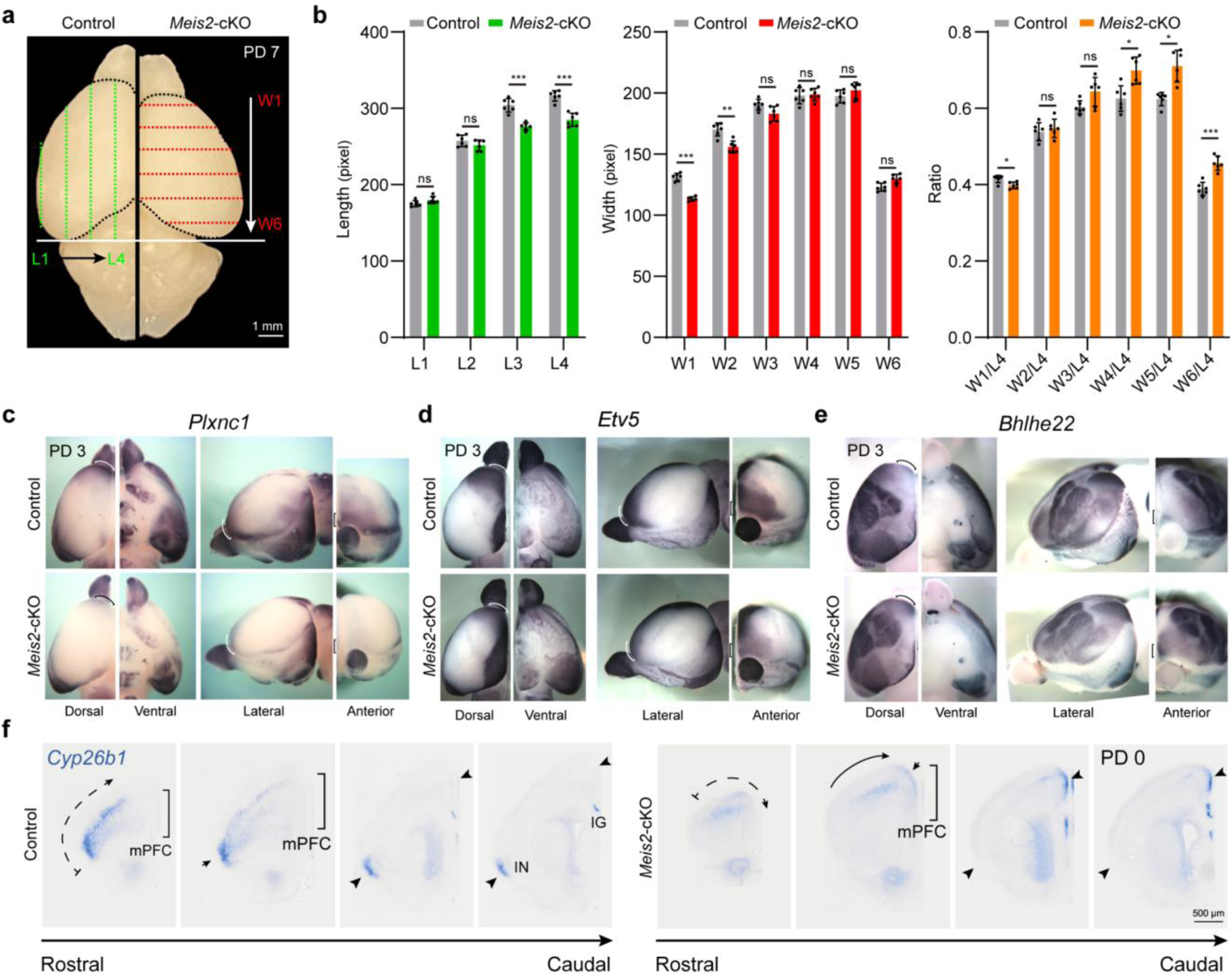
Altered rostral cortical regions in *Meis2* cKO mice. **a,** Dorsal view of PD 7 mouse brains showing the cortical grid used to measure rostro–caudal length (L1–L4) and medio–lateral width (W1–W6) in control and *Meis2* cKO mice. **b,** Quantification of cortical length, width and W/L ratios across defined grid positions. *Meis2* cKO mice exhibit reduced rostral cortical length (L3, L4), decreased medial width (W1, W2) and lower W/L ratios (W1/L4). **c–e,** Expression of regional markers *Plxnc1* (PFC), *Etv5* (MC), and *Bhlhe22* (sensory cortex) in control and *Meis2* cKO brains. Markers show changes in the rostral regions (Curves and straight lines). **f,** *Cyp26b1* expression along the rostro–caudal axis at PD 0 in control and *Meis2* cKO coronal brain sections. Arrows indicate *Cbln2* expression in the PFC and MC, and *Cyp26b1* expression in the MC. Curved arrows indicate *Cyp26b1* ectopic medial expression. Scale bars, see images; statistical details, see Methods.

**Extended Data Figure. 13.**
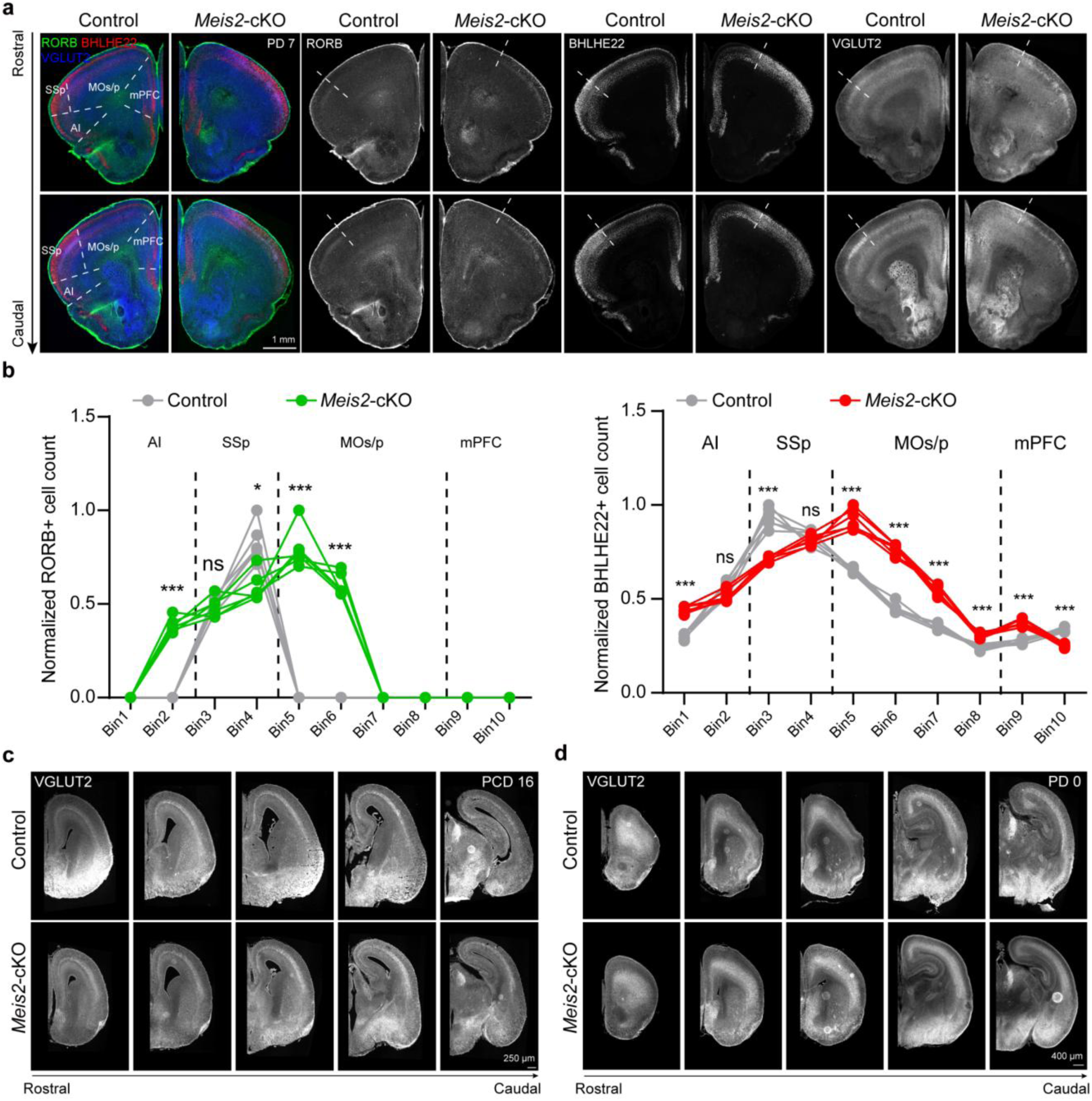
Expression of RORB, BHLHE22 and VGLUT2 in *Meis2* cKO mice. **a,** IHC of RORB (green), BHLHE22 (red) and VGLUT2 (blue) in coronal sections of PD 7 mouse brains from control and *Meis2* cKO mice. **b,** Quantification of fluorescence intensity across cortical bins (lateral Bin1 to medial Bin10; Bin1–2, AI; Bin3–4, SSp; Bin5–8, MOs/p; Bin9–10, mPFC). In *Meis2* cKO mice, RORB-, BHLHE22- and VGLUT2-positive signals expand into MC areas. **c,d,** IHC of VGLUT2 in coronal sections of control and *Meis2* cKO mice at PCD 16 and PD 0, spanning the rostro–caudal cortical axis. Scale bars, see images; statistical details, see Methods.

**Extended Data Figure. 14.**
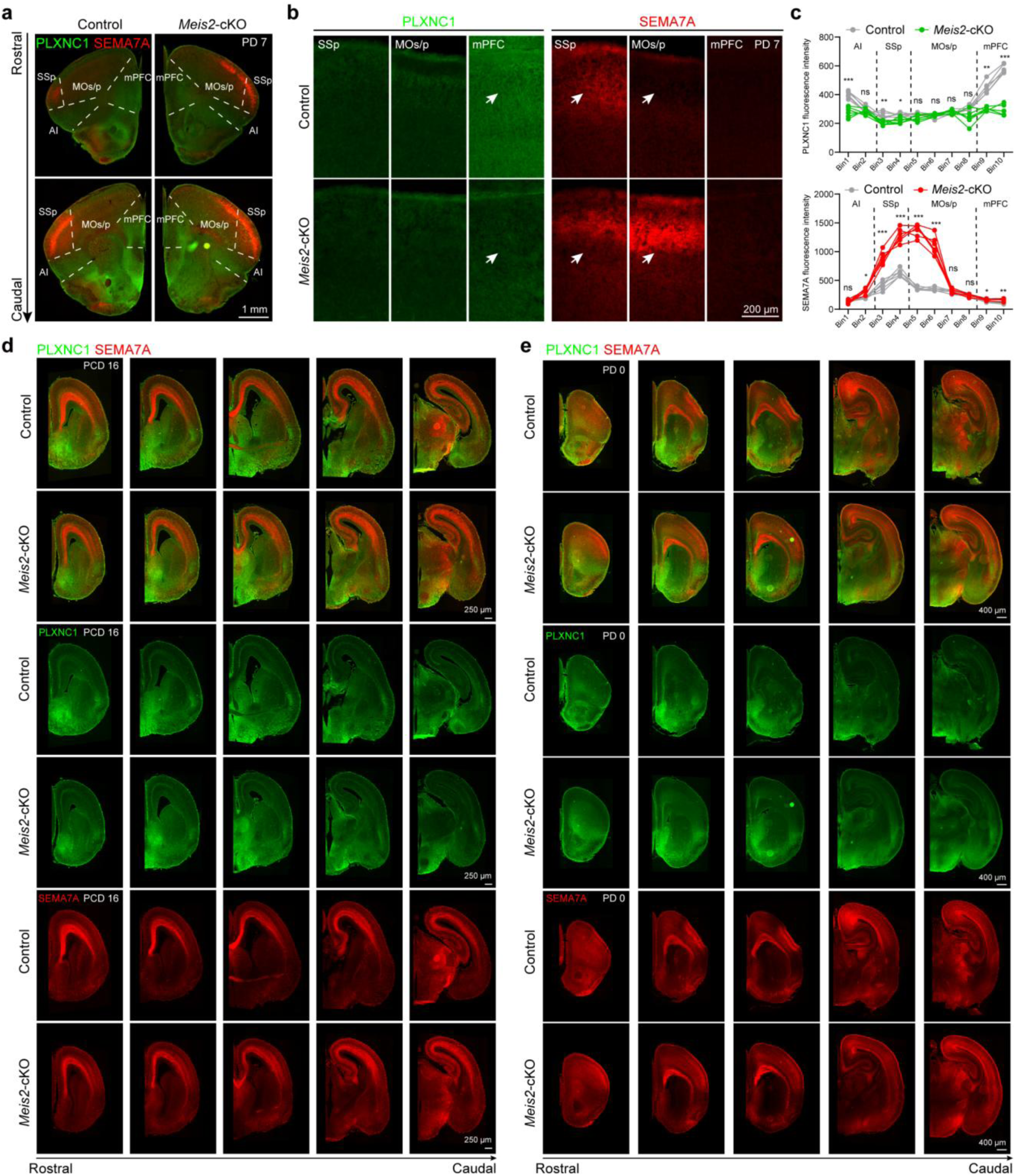
Expression of PLXNC1 and SEMA7A in *Meis2* cKO mice. **a, b,** IHC of PLXNC1 (green) and SEMA7A (red) in coronal sections of PD 7 mouse brains. In control mice, PLXNC1 is enriched in the mPFC and SEMA7A in the SSp; these spatial patterns are reduced or altered in *Meis2* cKO mice. Markers show changes in the SSp, MOs/p and mPFC (white arrows). **c,** Quantification of fluorescence intensity across cortical bins (lateral Bin1 to medial Bin10; Bin1–2, AI; Bin3–4, SSp; Bin5–8, MOs/p; Bin9–10, mPFC). In *Meis2* cKO mice, PLXNC1 expression is reduced in the mPFC, whereas SEMA7A expression expands into MC regions. **d,e,** IHC of PLXNC1 (green) and SEMA7A (red) in coronal sections of control and *Meis2* cKO mice at PCD 16 and PD 0, spanning the rostro–caudal axis. Single-channel images for PLXNC1 and SEMA7A show corresponding expression differences in the cortex of *Meis2* cKO mice compared to controls. Scale bars, see images; statistical details, see Methods.

**Extended Data Figure. 15.**
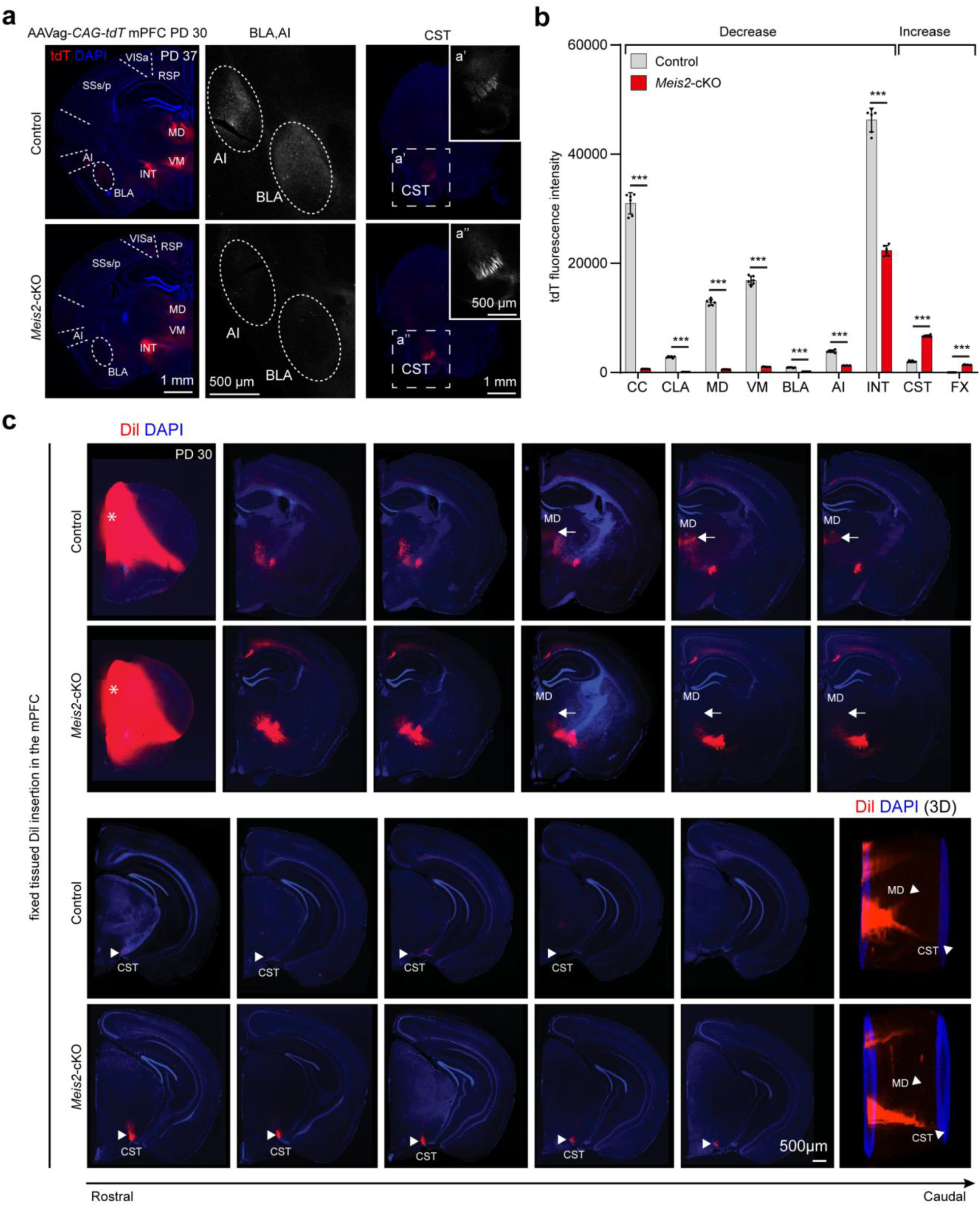
Axonal projection defects from the mPFC in *Meis2* cKO mice. **a,** Anterograde tracing of mPFC projection neurons using AAVag-*CAG–tdTomato* injected at PD 30. After 7 days, control projections target prefrontal connectivity regions (MD, VM, AI, BLA), whereas in *Meis2* cKO mice, labeling is more obvious in CST, indicating that the former mPFC region now exhibits MC-like projections. Enlarged views (a′–a′′) highlight tdTomato signals in the BLA, AI and CST. **b,** Quantification of projection density (tdTomato fluorescence intensity) across brain regions. Inputs to prefrontal-associated targets (CC, MD, VM, BLA, CLA, AI, INT) are reduced, whereas those to motor-associated regions (CST, FX) are increased in *Meis2* cKO mice. **c,** DiI placement in the mPFC (asterisks) in fixed control and *Meis2* cKO brains at PD 30. Rostral-to-caudal series of coronal sections and 3D reconstructions show the Dil-labeled projection fibers (red) and nuclei (DAPI, blue). Compared with controls, *Meis2* cKO mice exhibit altered axonal trajectories, characterized by reduced long-range connectivity between the mPFC and MD, and increased CST. Arrows and arrowheads indicate MD and CST, respectively. Scale bars, see images; statistical details, see Methods. CC, corpus callosum; CLA, claustrum; MD, mediodorsal thalamic nucleus; VM, ventral medial thalamic nucleus; BLA, basolateral amygdala; AI, agranular insular cortex; INT, internal capsule; CST, corticospinal tract; FX, the columns of the fornix.

**Extended Data Figure. 16.**
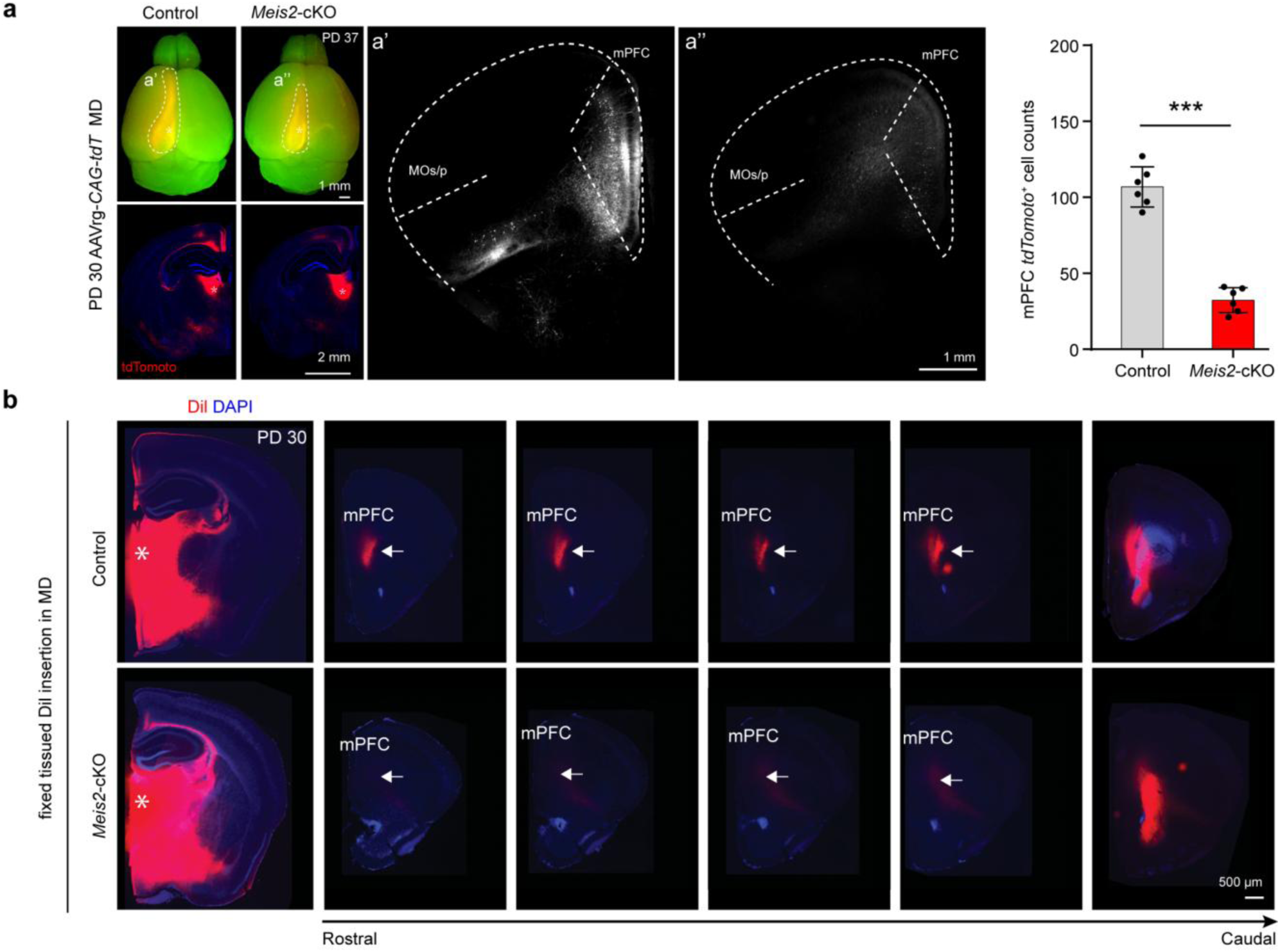
Axonal projection defects from the MD in *Meis2* cKO mice. **a,** Retrograde tracing from the MD showing reduced retrogradely labeled neurons in the mPFC of *Meis2* cKO mice, confirming loss of mPFC–thalamic projections. Quantification demonstrates a significant reduction in labeled neurons. **b,** DiI placement in the MD (asterisks) of fixed control and *Meis2* cKO brains at PD 30. Rostral-to-caudal series of coronal sections show Dil-labeled projection fibers (red) and nuclei (DAPI, blue). Compared with controls, *Meis2* cKO mice display altered axonal trajectories and reduced long-range projections between the MD and mPFC. Arrows indicate mPFC. Scale bars, see images; statistical details, see Methods.

**Extended Data Figure. 17.**
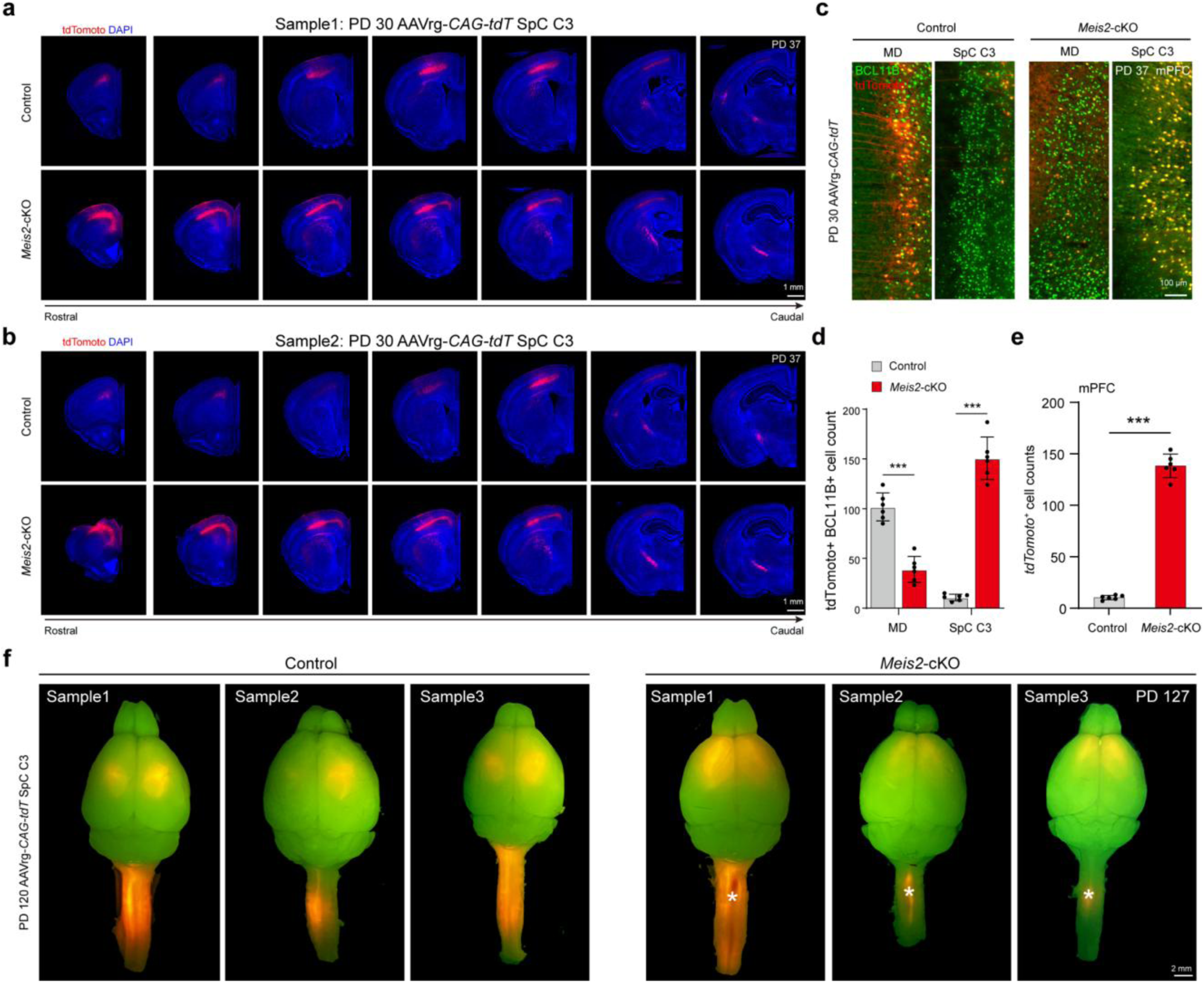
Additional replicates of retrograde tracing from SpC C3. **a, b,** AAVrg–*CAG–tdTomato* was injected into the spinal cord C3 at PD 30 to label retrogradely projecting cortical neurons. **c, d,** Retrograde tracing of corticothalamic and corticospinal projections in *Meis2* cKO mice. Representative coronal sections of the mPFC at PD 37 following retrograde labeling of L5ET neurons after PD 30 AAVrg-*CAG-tdTomato* injection into the MD or SpC C3. Sections were immunostained for BCL11B (green) and retrogradely labeled neurons (tdTomato, red). In controls, thalamic injections predominantly label corticothalamic neurons, whereas SpC injections label corticospinal neurons. In *Meis2* cKO mice, corticothalamic projections are markedly reduced, whereas corticospinal projections are increased. **e,** Quantification of Fig. 3g showing results from retrograde tracing of SpC C3 injections. In *Meis2* cKO mice, tdTomato-labeled corticospinal-projecting neurons are ectopically detected in the mPFC region, reflecting a shift of projection identity toward motor cortex projection neurons. **f,** AAVrg–*CAG–tdTomato* was injected into the SpC C3 at PD 120 to label retrogradely projecting cortical neurons. Scale bars, see images; statistical details, see Methods.

**Extended Data Figure. 18.**
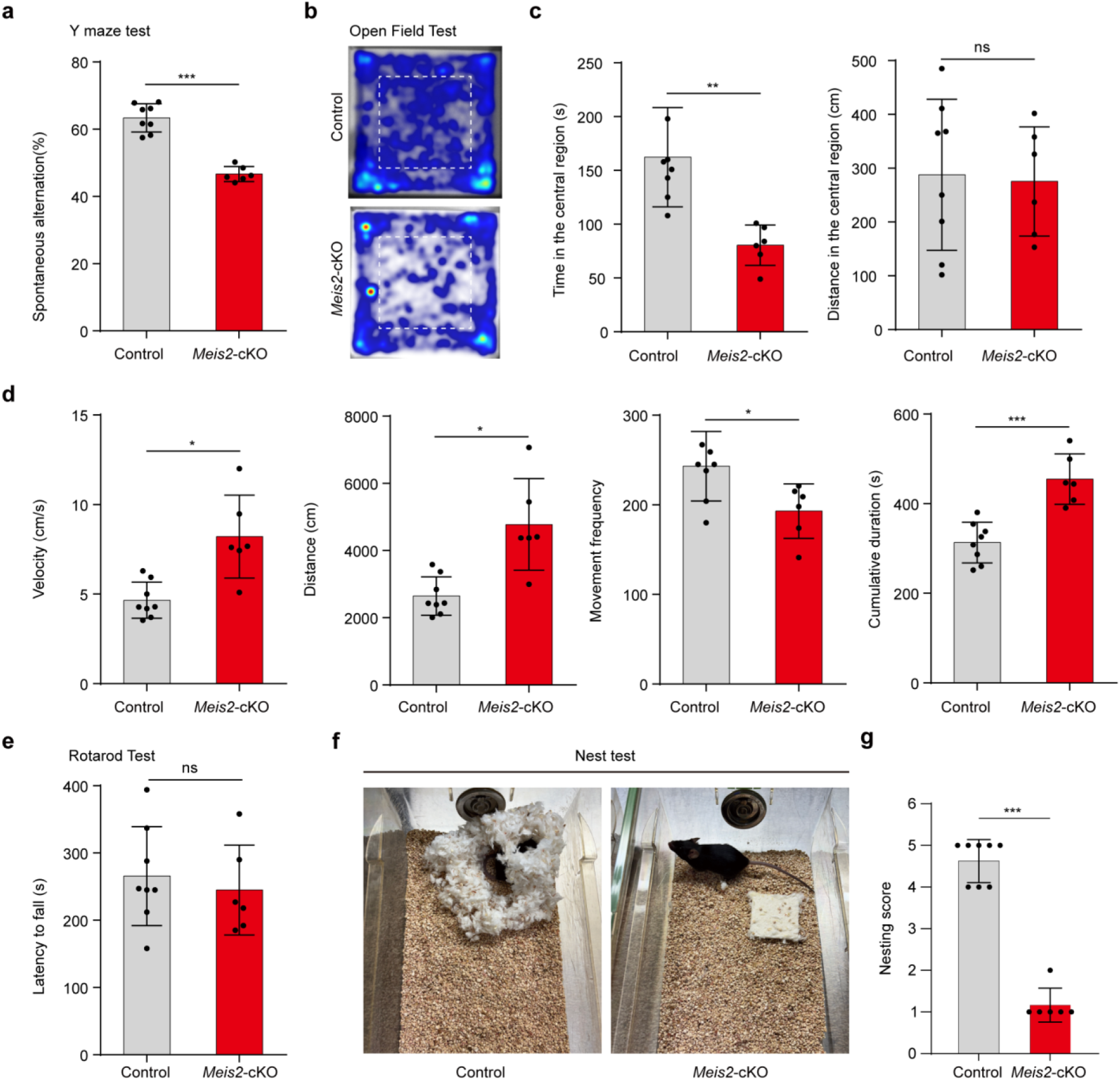
Behavioral characterization of *Meis2* cKO mice. **a,** Y-maze test of spontaneous alternation showing reduced alternation rates in *Meis2* cKO mice, consistent with impaired spatial working memory. **b,** Heat maps from the open-field test showing locomotor trajectories. **c,** *Meis2* cKO mice spend less time and travel shorter distances in the center of the open field, reflecting increased anxiety-like behavior. **d,** Locomotor activity metrics in the open-field test—including velocity, total distance, movement frequency and cumulative movement duration—show elevated activity levels in *Meis2* cKO mice. **e,** Rotarod test showing no significant difference in latency to fall between *Meis2* cKO and control mice, indicating preserved motor coordination and balance. **f, g,** Nest-building assay showing impaired nest-construction behavior in *Meis2* cKO mice. Representative images (f) and quantification (g) indicate markedly reduced nesting performance compared with control littermates. Statistical details, see Methods.

**Extended Data Figure. 19.**
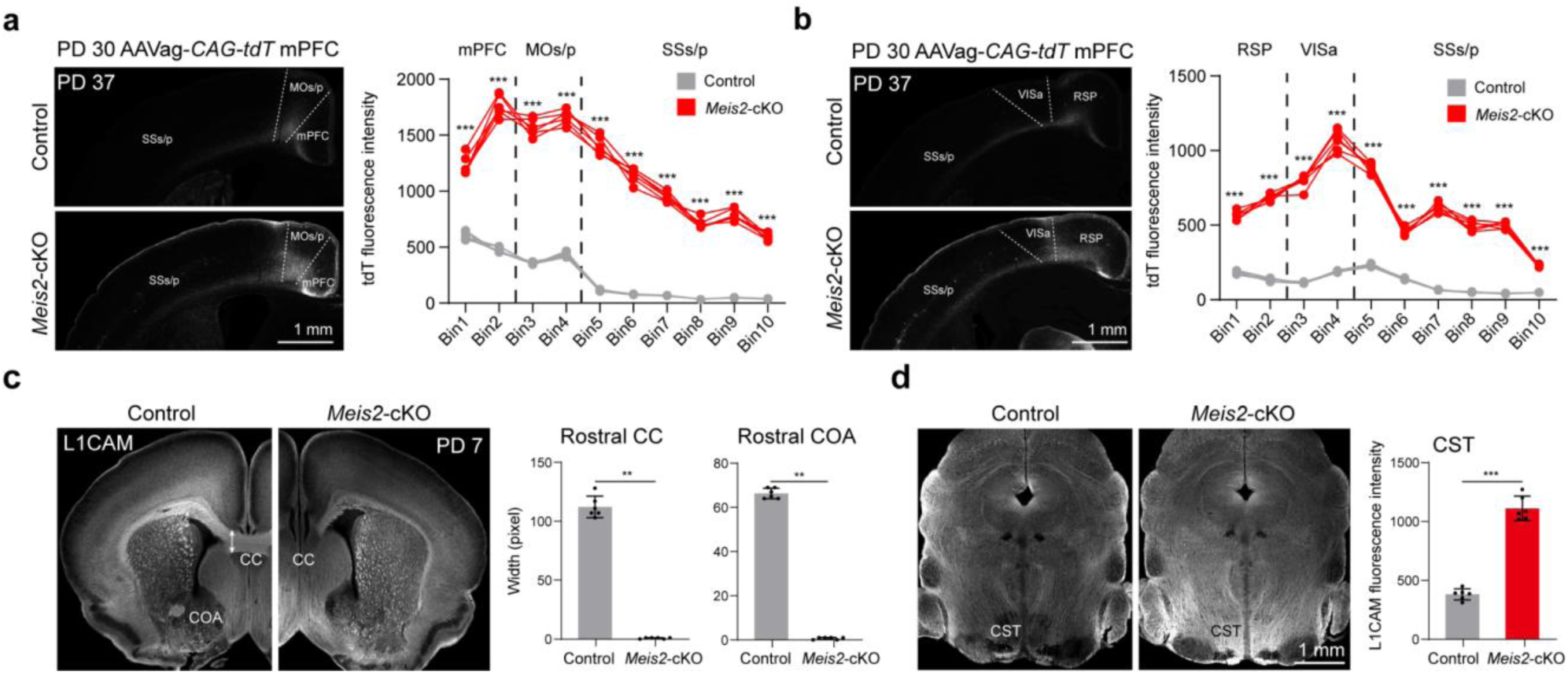
Altered axonal connectivity in *Meis2* cKO mice. **a, b,** Quantification of tdTomato signal across ten cortical bins showing altered axonal projection patterns from contralateral to ipsilateral hemispheres in *Meis2* cKO mice. **c,** Coronal brain sections at PD 7 IHC for L1CAM, labeling major axon tracts including the CC, COA and CST. Control mice show robust midline crossing of callosal axons and well-defined COA structures, both of which are markedly reduced in the rostral cortex of *Meis2* cKO mice. **d,** Quantification of rostral CC and COA widths and L1CAM fluorescence intensity in the CST. *Meis2* cKO mice exhibit severe reductions in interhemispheric tracts (CC and COA), accompanied by increased CST signal. Scale bars, see images; statistical details, see Methods.

**Extended Data Figure. 20.**
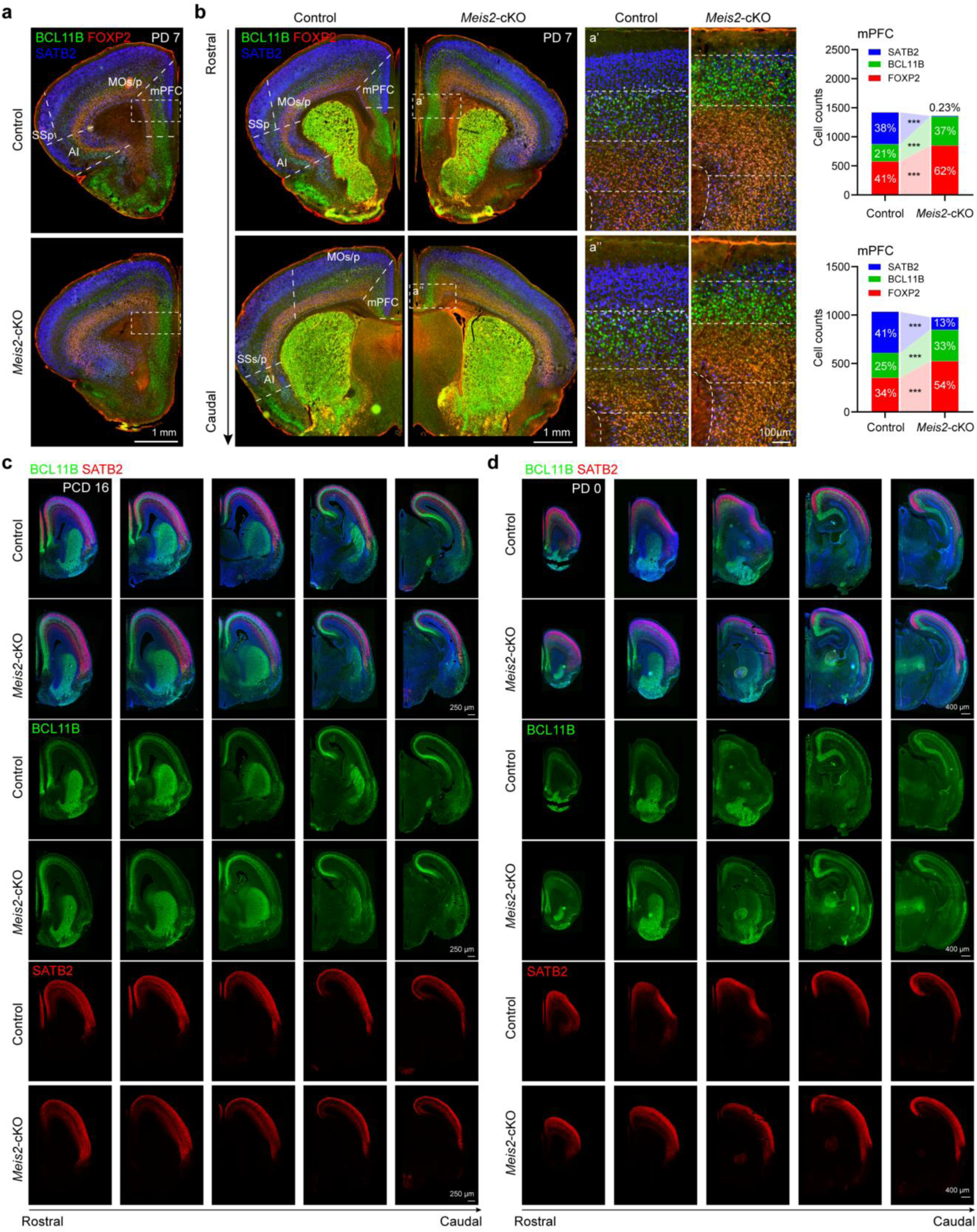
Altered cortical layer organization in the mPFC of *Meis2* cKO mice. **a,b,** IHC of PD 7 coronal brain sections stained for layer-specific markers: SATB2 (upper layers, blue), BCL11B (layer V, green) and FOXP2 (layer VI, red) in control and *Meis2* cKO mice. Enlarged views (a′–a′′) of the mPFC show altered distributions of layer markers. Quantification of SATB2⁺, BCL11B⁺ and FOXP2⁺ neurons in the mPFC demonstrates a significant reduction of SATB2⁺ upper-layer neurons and an increase in BCL11B⁺ and FOXP2⁺ deep-layer neurons in *Meis2* cKO mice. **c, d,** IHC of BCL11B (green) and SATB2 (red) in coronal sections of control and *Meis2* cKO mice at PCD 16 (c) and PD 0 (d), spanning the rostro–caudal cortical axis. Single-channel images show expression differences between *Meis2* cKO and control cortices. Scale bars, see images; statistical details, see Methods.

**Extended Data Figure. 21.**
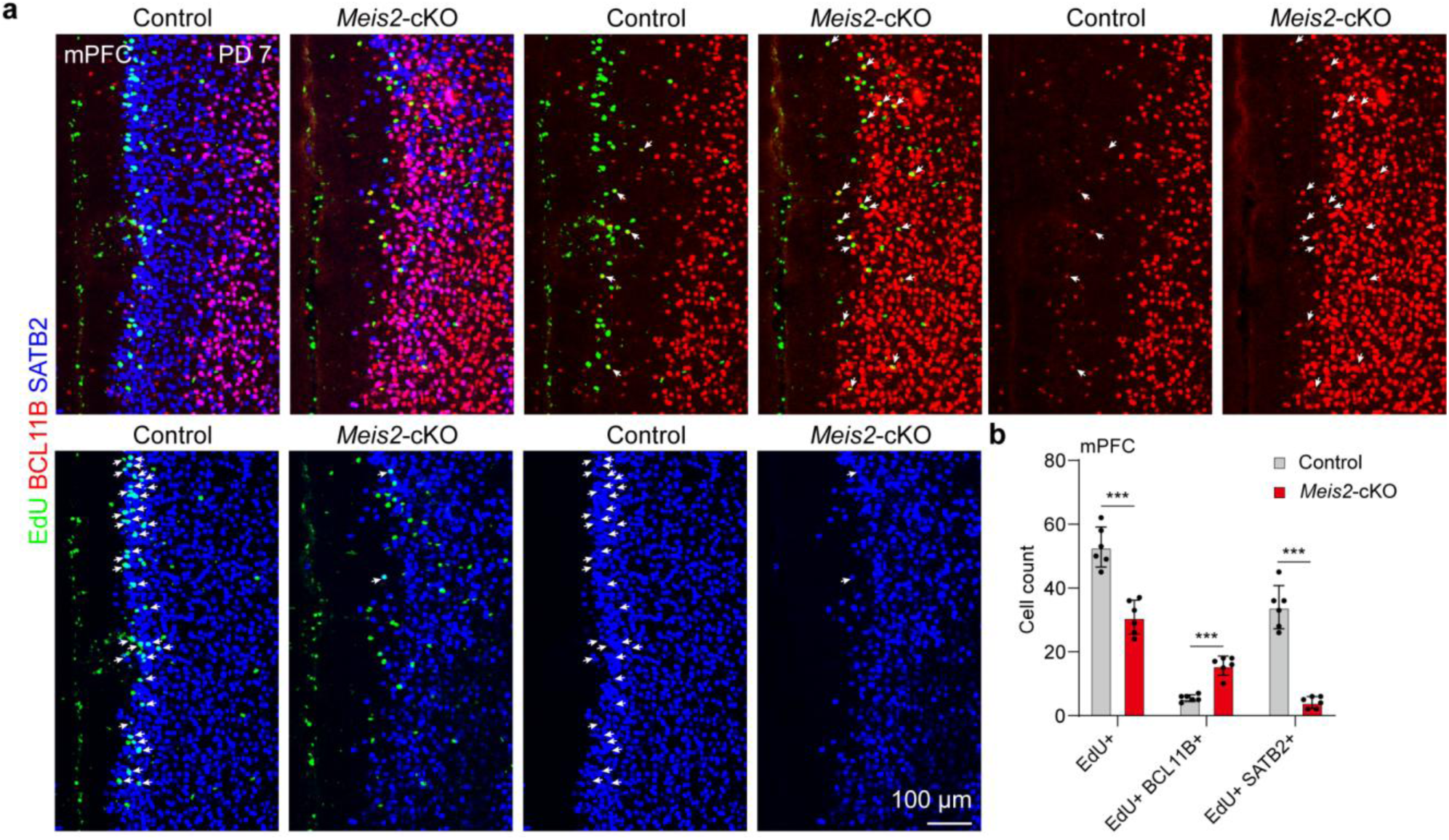
*Meis2* regulates cortical layers organization in the mPFC. **a,** IHC of PD 7 mPFC sections from control and *Meis2* cKO mice injected with EdU at PCD 16. Sections were co-stained for EdU (green), BCL11B (red) and SATB2 (blue). White arrows indicate EdU⁺ cells co-expressing layer-specific markers. **b,** Quantification of EdU⁺ cells, EdU⁺BCL11B⁺ cells and EdU⁺SATB2⁺ cells in the mPFC. *Meis2* cKO mice show a reduction in EdU⁺ and EdU⁺SATB2⁺ cells, accompanied by an increase in EdU⁺BCL11B⁺ cells. Scale bars, see images; statistical details, see Methods.

**Extended Data Figure. 22.**
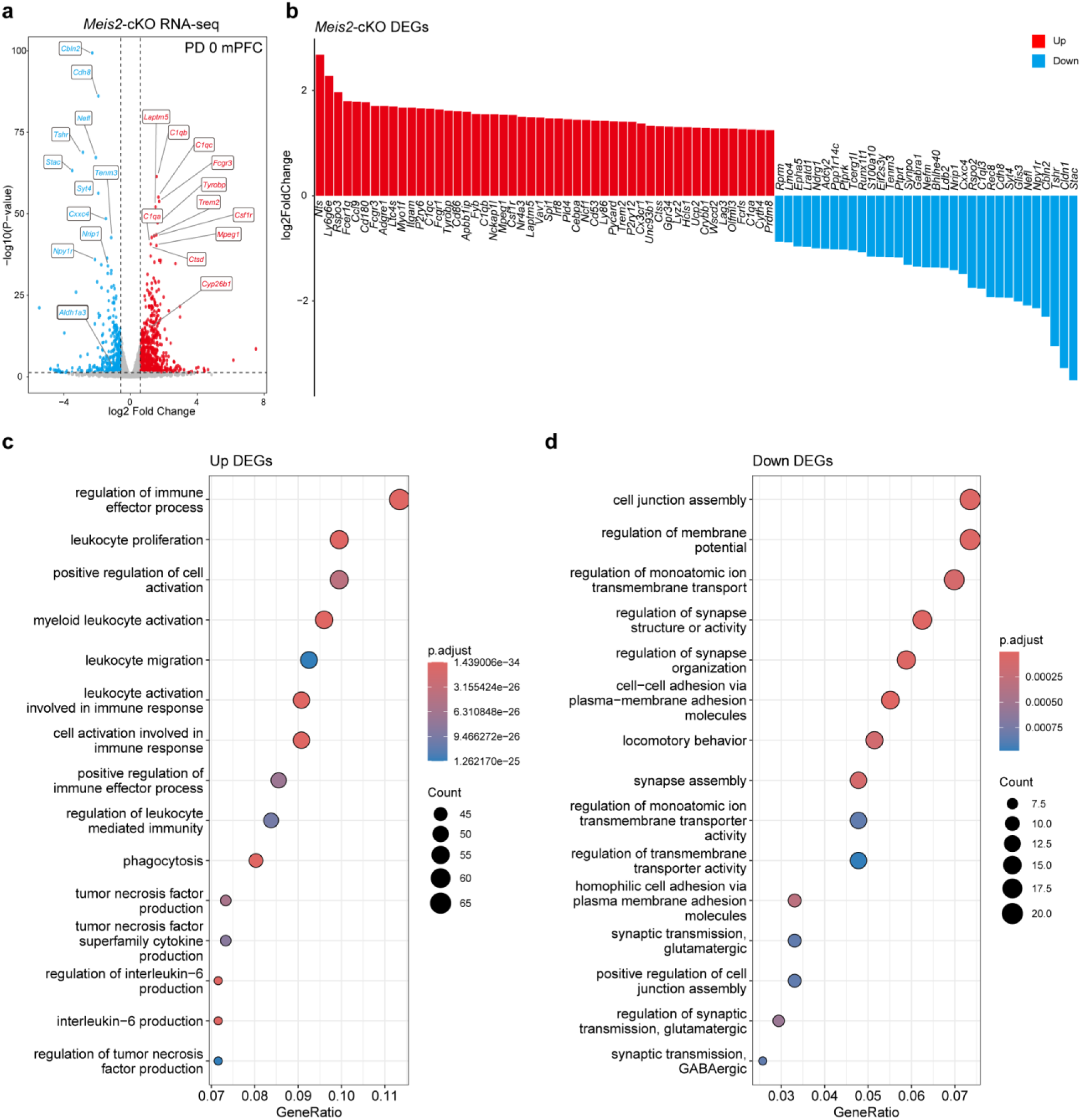
Transcriptomic alterations in the mPFC of *Meis2* cKO mice at PD 0. **a,** RNA-seq analysis of the mPFC from *Meis2* cKO and control mice at PD 0. Several mPFC-enriched synaptic genes (*Cbln2*, *Cdh8*, *Syt4*) and the RA synthase *Aldh1a3* are significantly downregulated. **b,** Bar graph of DEGs in the *Meis2* cKO mPFC at PD 0. Upregulated genes are shown in red, and downregulated genes in blue (log₂ fold-change). **c,** GO biological process enrichment of upregulated DEGs, highlighting immune-related processes such as leukocyte activation, cytokine signaling and inflammatory responses. **d,** GO biological process enrichment of downregulated DEGs, highlighting synaptic and neuronal communication pathways including glutamatergic transmission, synapse assembly and cell junction organization.

**Extended Data Figure. 23.**
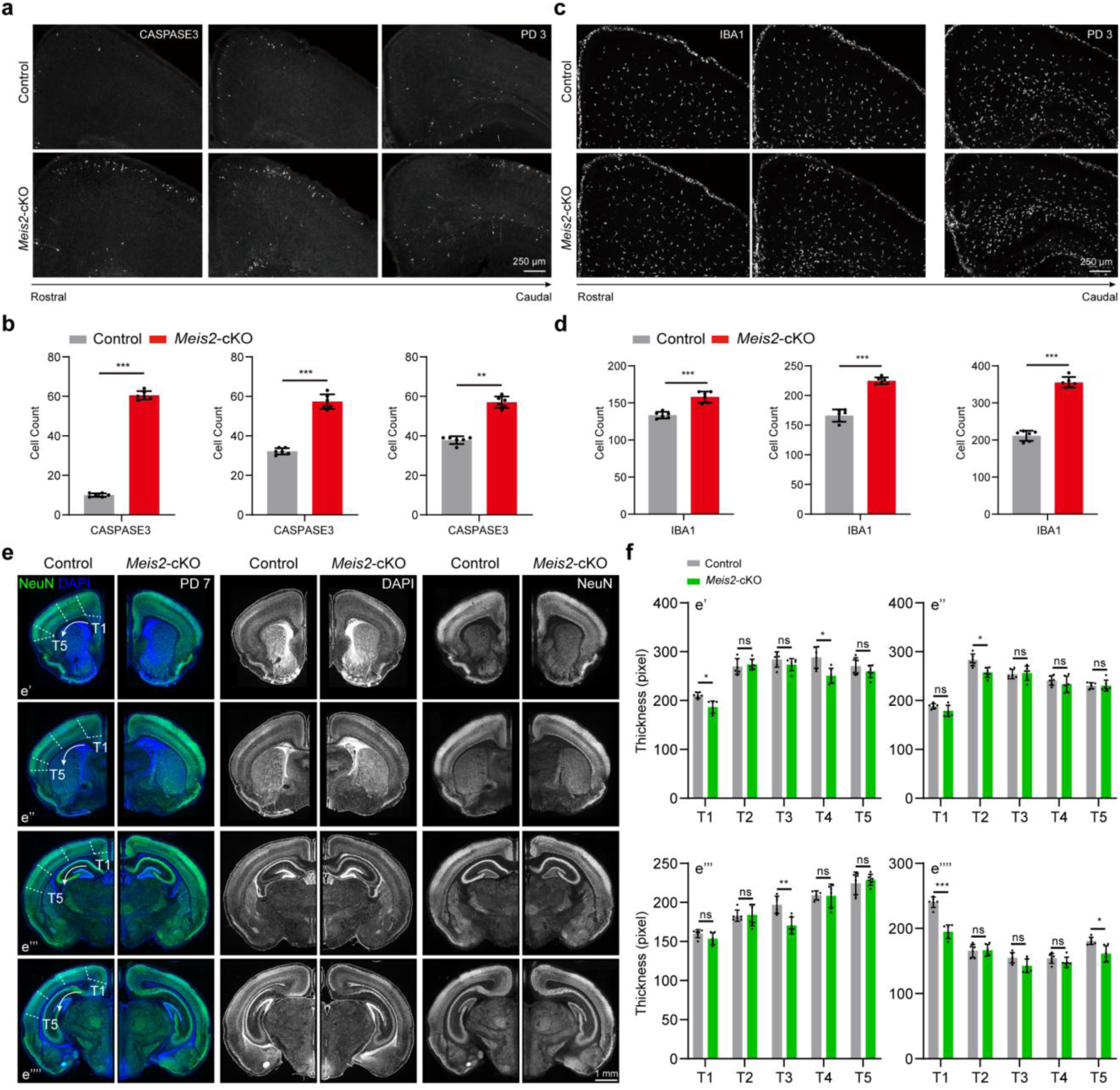
Increased neuronal apoptosis in *Meis2* cKO mice. **a,b,** IHC and quantification of CASPASE3⁺ cells in rostral-to-caudal cortical sections at PD 3 in control and *Meis2* cKO mice. *Meis2* cKO mice show increased CASPASE3 immunoreactivity, particularly at rostral regions. **c,d,** IHC and quantification of IBA1⁺ cells in rostral-to-caudal cortical sections at PD 3 in control and *Meis2* cKO mice, showing elevated microglial abundance and activation. **e,** Coronal sections from rostral-to-caudal cortex at PD 7 stained for NeuN (green) and DAPI (blue), comparing neuronal thickness (T1–T5) between control and *Meis2* cKO mice. **f,** Quantification of neuronal thickness across T1–T5, revealing a slight reduction in *Meis2* cKO mice. Scale bars, see images; statistical details, see Methods.

**Extended Data Figure. 24.**
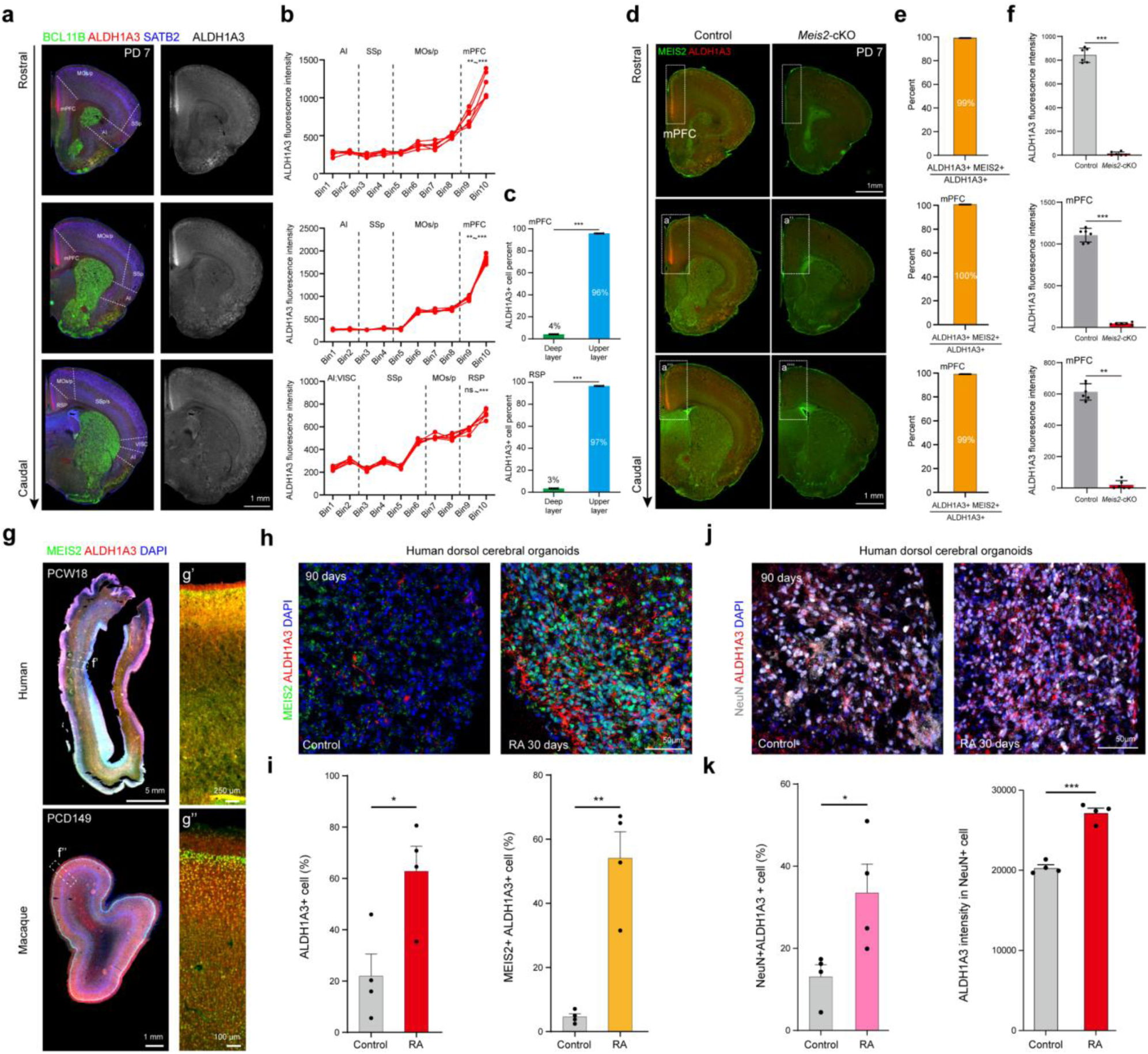
*MEIS2* promotes the generation of *ALDH1A3*⁺ neurons across species. **a,** IHC of ALDH1A3 (red), SATB2 (blue) and BCL11B (green) in PD 7 mouse brain, demonstrating strong ALDH1A3 signal in the mPFC upper layers and lower expression in adjacent cortical regions (SSp, MOs/p, VISC and AI). **b,** Quantification of ALDH1A3 fluorescence intensity across ten rostro–caudal cortical bins, revealing significant mPFC enrichment. **c,** ALDH1A3⁺ cells co-expressing SATB2 or BCL11B in the mPFC. Nearly all ALDH1A3⁺ cells co-express SATB2, whereas only a small fraction co-express BCL11B, indicating strong upper-layer enrichment in the mPFC. **d,** IHC of PD 7 brain sections from control and *Meis2* cKO mice showing ALDH1A3 (red) and MEIS2 (green) expression along the rostro–caudal axis of the mPFC. Enlarged views (a′–a′′′′) highlight the mPFC region, where ALDH1A3 signal is markedly reduced in *Meis2* cKO mice. **e,** Quantification of co-expression showing that more than 99% of ALDH1A3⁺ cells in the mPFC also express MEIS2, indicating near-complete overlap of expression. **f,** Quantification of ALDH1A3 fluorescence intensity across rostro–caudal sections of the mPFC showing a significant reduction of ALDH1A3 signal in *Meis2* cKO mice. **g,** IHC of MEIS2 and ALDH1A3 in human brain at PCW 18 and macaque brain at PCD 149. Enlarged views (d′–d′′) highlight ALDH1A3 and MEIS2 colocalization in the mPFC. **h, i,** IHC of MEIS2 and ALDH1A3 in 90-day human dorsal cerebral organoids. Compared with controls, RA treatment for 30 days markedly increases the proportion of ALDH1A3⁺ and MEIS2⁺ALDH1A3⁺ cells. **j,** IHC of 90-day human dorsal cerebral organoids treated with RA for 30 days, stained for NeuN (white), ALDH1A3 (red) and DAPI (blue). **k,** Quantification of NeuN⁺ALDH1A3⁺ cells and ALDH1A3 intensity in NeuN^+^ cells. RA treatment significantly increases the proportion of ALDH1A3⁺ neurons and ALDH1A3 intensity. Scale bars, see images; statistical details, see Methods.

**Extended Data Figure. 25.**
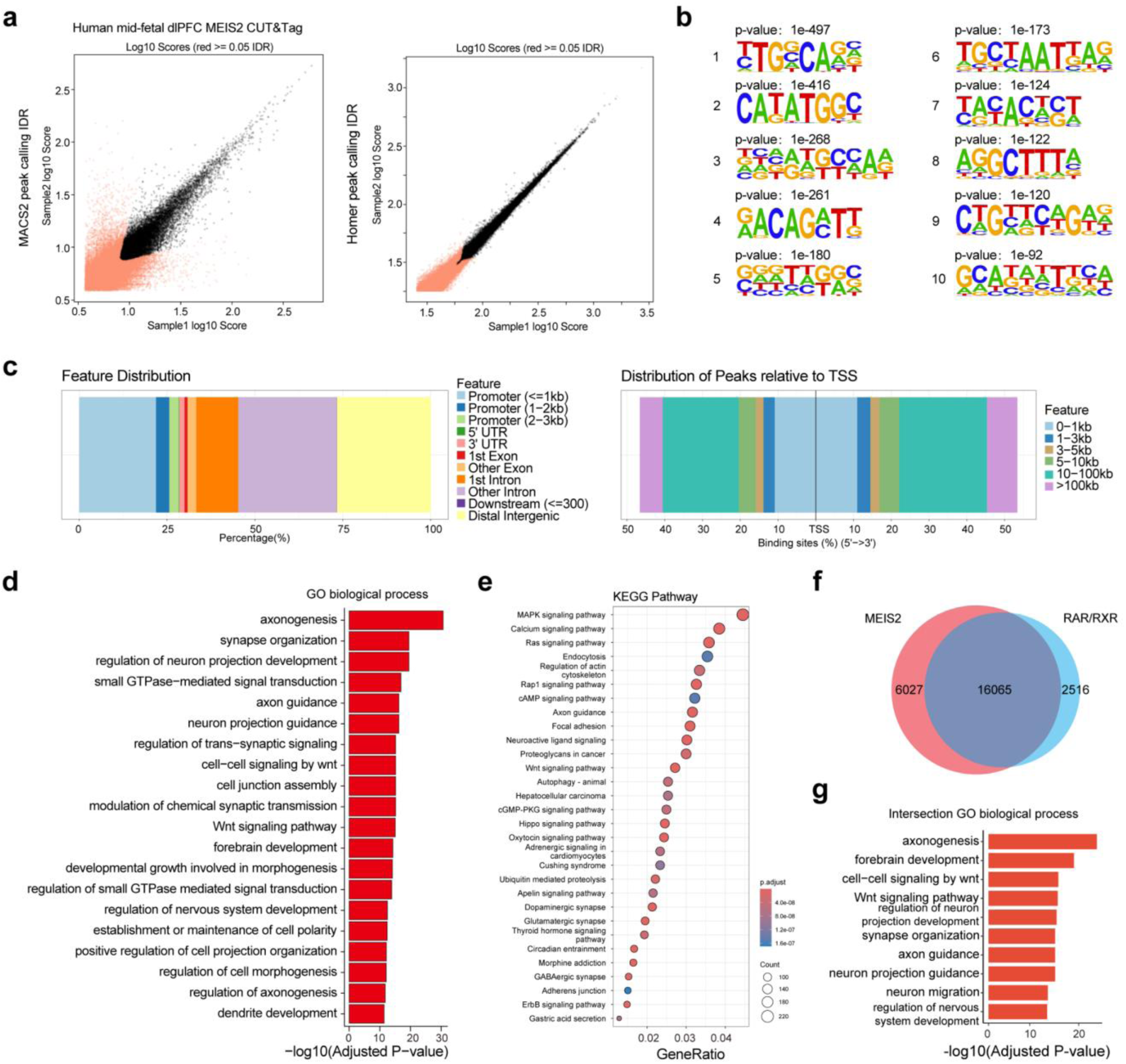
CUT&Tag profiling of MEIS2 in the human mid-fetal dlPFC. **a,** Scatter plots showing pairwise replicate reproducibility for CUT&Tag experiments targeting MEIS2. Peak calling was performed using MACS2 (top) and HOMER (bottom), followed by IDR analysis. Each point represents a genomic locus with log-transformed peak scores from replicate 1 (x-axis) and replicate 2 (y-axis). Black points denote reproducible peaks (IDR < 0.05), and red points indicate non-reproducible loci. Both algorithms show high concordance between biological replicates, supporting the robustness of MEIS2 CUT&Tag profiling. **b,** Top five de novo motifs enriched in MEIS2 binding peaks in the human mid-fetal dlPFC. **c,** Genomic annotation of MEIS2 binding peaks, showing distribution across promoters, untranslated regions, exons, introns, and distal intergenic regions. The majority of peaks are located within 3 kb upstream of transcription start sites. **d, e,** GO biological process and KEGG pathway enrichment of MEIS2-bound genes annotated by ChIPseeker as proximal targets. Dot size represents the number of genes per category, and color indicates adjusted significance. **f,** Venn diagram showing the overlap between MEIS2- and RAR/RXR-binding genes identified by CUT&Tag in the human mid-fetal dlPFC. The intersection (16,065 genes) indicates co-binding by MEIS2 and RARs. **g,** GO biological process enrichment of genes co-bound by MEIS2 and RAR/RXR.

**Extended Data Figure. 26.**
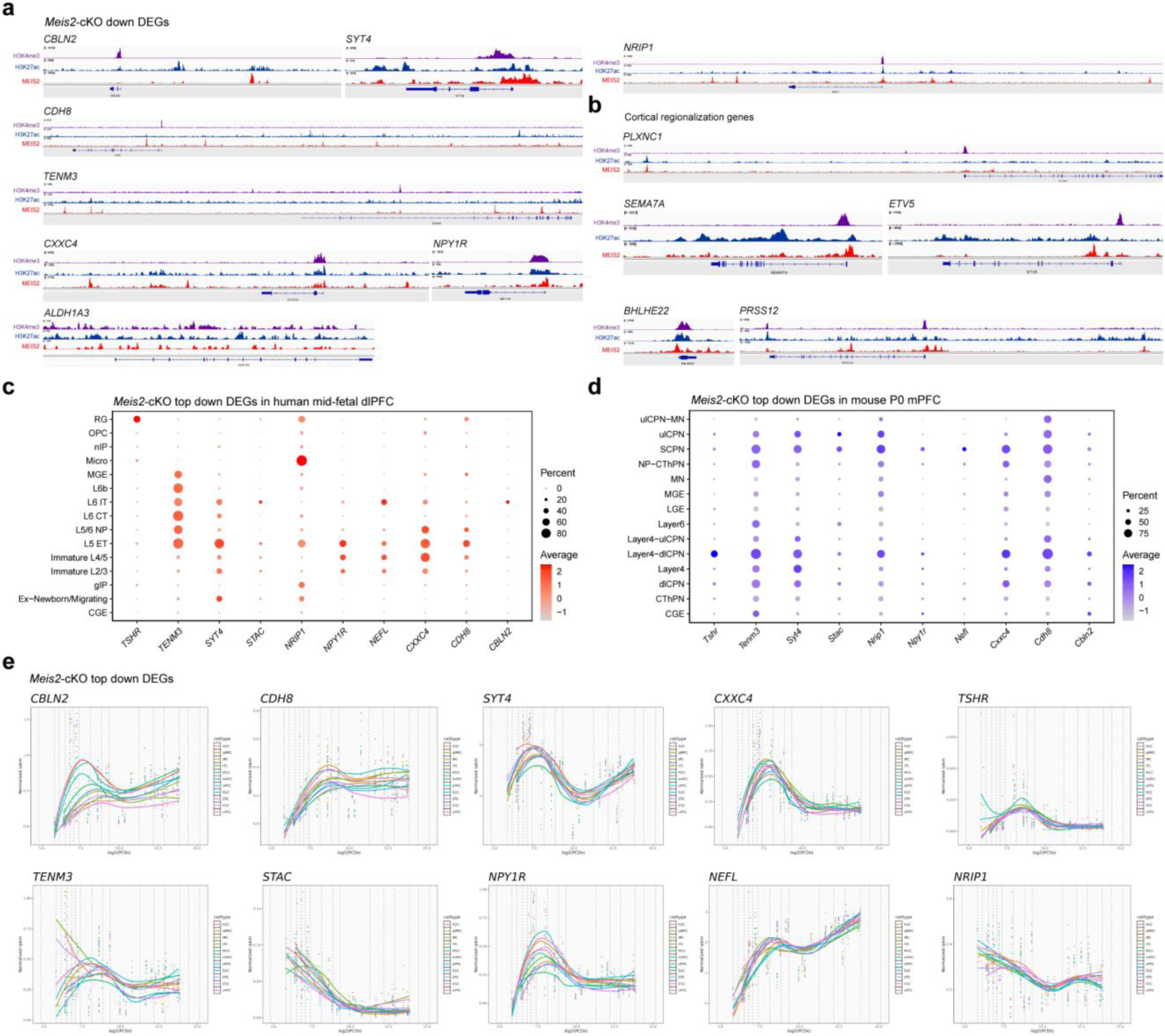
Characterization of *Meis2* cKO down DEGs. **a,** CUT&Tag profiles of H3K4me3, H3K27ac and MEIS2 binding in the human mid-fetal dlPFC. Genome browser views show *Meis2* cKO downregulated DEGs. **b,** CUT&Tag profiles of H3K4me3, H3K27ac and MEIS2 binding in the human mid-fetal dlPFC. Genome browser views show representative cortical regionalization genes. **c,** Dot plot showing cell-type expression of top *Meis2* cKO downregulated DEGs in the human mid-fetal dlPFC. **d,** Dot plot showing cell-type expression of top *Meis2* cKO downregulated DEGs in the mouse mPFC at PD 0. Dot size represents the proportion of expressing cells, and color scale indicates mean expression level. **e,** Spatiotemporal expression profiles of the top downregulated DEGs in *Meis2* cKO across human brain regions and developmental stages. Each panel shows normalized expression levels plotted against post-conceptional weeks, with fitted curves representing developmental trajectories. The dashed lines indicate the periods of human development and adulthood.

**Extended Data Figure. 27.**
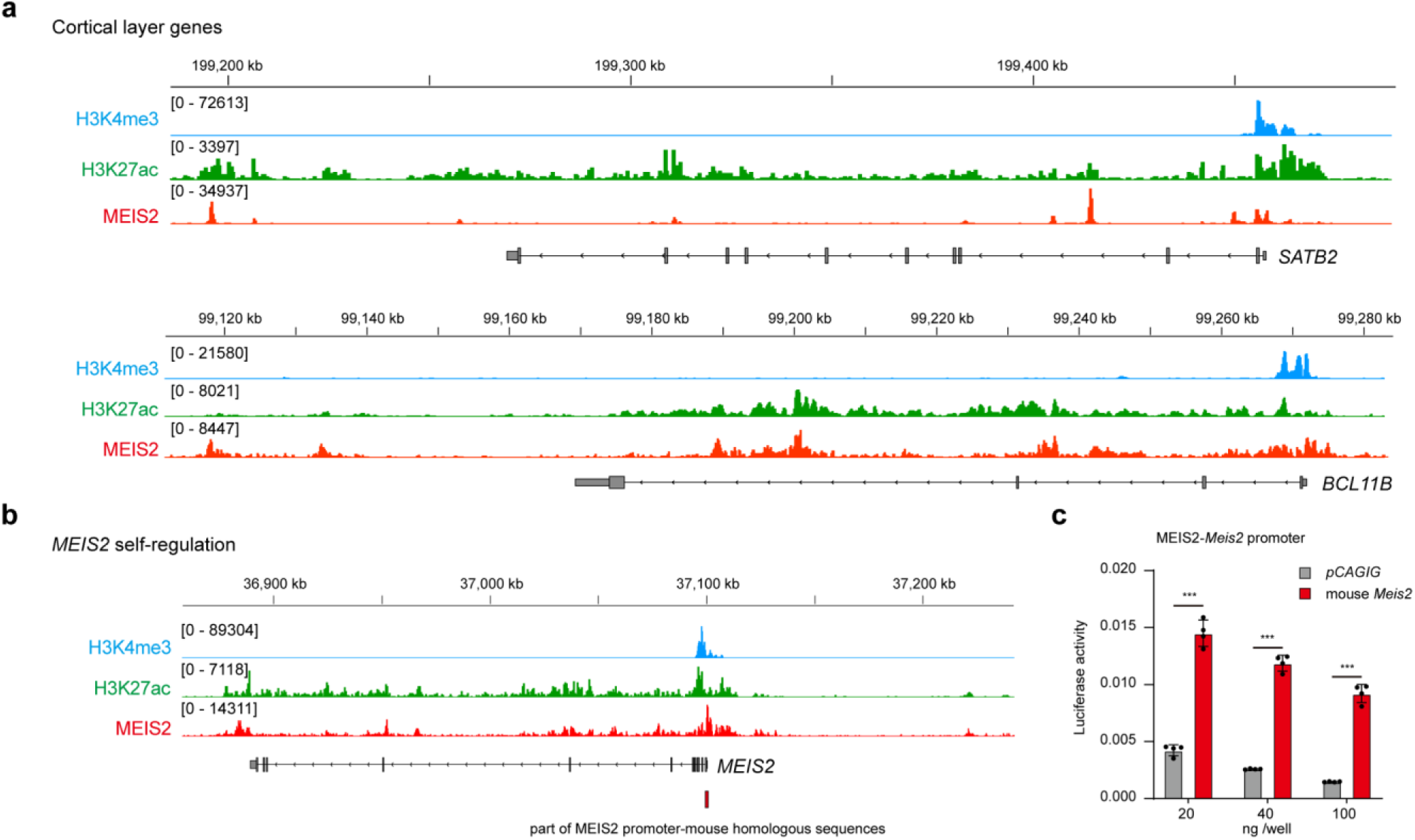
MEIS2 CUT&Tag profiles of *SATB2*, *BCL11B* and *MEIS2* in the human mid-fetal dlPFC. **a,** CUT&Tag profiles of H3K4me3, H3K27ac and MEIS2 binding in the human mid-fetal dlPFC. Genome-browser views show representative cortical layer genes (*SATB2*, *BCL11B*). **b,** CUT&Tag profiles of H3K4me3, H3K27ac and MEIS2 binding in the human mid-fetal dlPFC. Genome-browser views show MEIS2 binding peaks at the *MEIS2* locus. **c,** Mouse *Meis2* promoter luciferase assay in Neuro2a cells with increasing concentrations of mouse MEIS2. Statistical details, see Methods.

**Extended Data Figure. 28.**
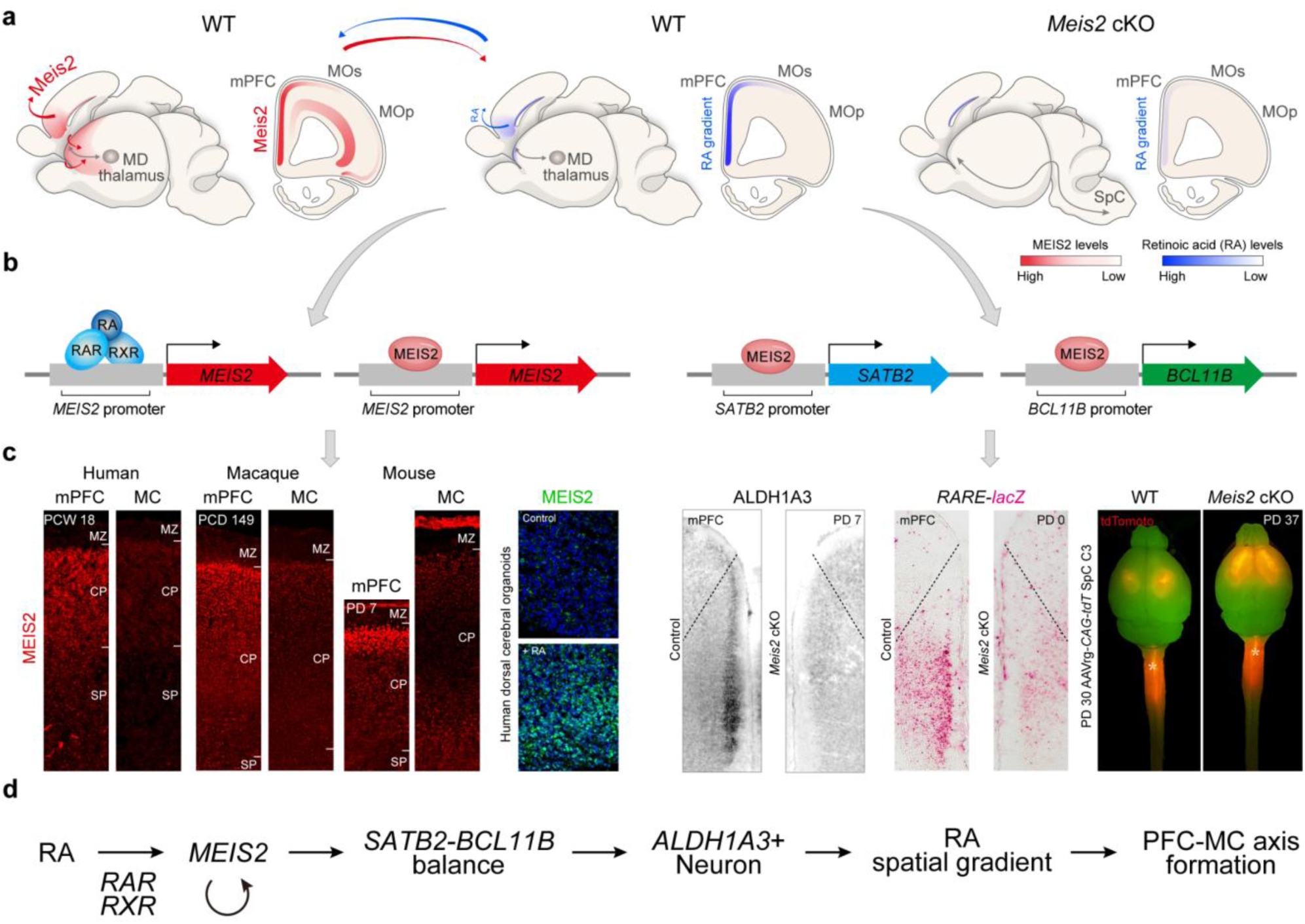
*MEIS2* maintains prefrontal RA gradients and regional identity across species. **a,** Schematic model illustrating *MEIS2*-dependent maintenance of RA gradients and prefrontal connectivity. In wild-type mice, *Meis2* is enriched in the mPFC and promotes RA synthesis, sustaining high anterior and low caudal RA levels. In *Meis2* cKO mice, RA gradients collapse, accompanied by loss of mPFC–MD thalamic projections and increased CST. **b,** *MEIS2* and RA form a transcriptional feedback circuit. RA–RAR/RXR signaling induces *MEIS2* transcription, while MEIS2 regulates *SATB2* and *BCL11B* promoters to specify upper- and deep-layer neuronal identities, respectively. **c,** (1) MEIS2 expression in MC–PFC axis of human (PCW 18), macaque (PCD 149) and mouse (PD 7). MEIS2 is enriched in the PFC and reduced in MC. (2) Human cortical organoids treated with RA show significantly increased *MEIS2* expression. (3) In *Meis2* cKO mice, ALDH1A3 expression and RARE–LacZ reporter activity in the mPFC are significantly reduced, accompanied by (4) PFC-motor regional reprogramming. **d,** Working model. RA activates RAR/RXR signaling, inducing *MEIS2* expression which can also self-regulate (Fig.2; Extended Data Fig. 10; Extended Data Fig. 27b,c). *MEIS2* regulates *SATB2*–*BCL11B* balance (Fig.4b; Extended Data Fig.20; Extended Data Fig. 27a), establishing an RA-rich mPFC and RA-poor MC (Extended Data Fig. 10a). This *MEIS2*-dependent autoregulatory loop underlies cortical RA gradient formation and MC–PFC arealization (Fig.4d,e; Fig.3f,g; Extended Data Fig.17).

## Methods

### Human tissue and ethical approval

Postmortem human brain specimens were obtained from the Department of Neuroscience at Yale University School of Medicine, the Birth Defects Research Laboratory at the University of Washington, Advanced Bioscience Resources Inc. (ABR), the Human Brain Collection Core (HBCC), the Brain and Tissue Bank at the University of Maryland, the MRC–Wellcome Trust Human Developmental Biology Resource at the Institute of Human Genetics, University of Newcastle (UK), and the Human Fetal Tissue Repository at the Albert Einstein College of Medicine (AECOM). All tissue was collected with informed consent from parents or next of kin and under protocols approved by the institutional review boards of Yale University School of Medicine, the National Institutes of Health, and the corresponding institutions from which specimens were obtained. Tissue processing complied with NIH ethical guidelines and the principles of the WMA Declaration of Helsinki (https://www.wma.net/policies-post/wma-declaration-of-helsinki/). De-identified information for each specimen is provided in **Supplementary Table 1**.

### Macaque tissue and ethical approval

Rhesus macaque brain samples were collected postmortem at PCD 149 from Yale MacBrain Resource Center (MBRC). Whole slabs or whole hemispheres were post-fixed in 4% paraformaldehyde (PFA) for 48 hours and then cryoprotected in an ascending sucrose gradient (10%, 20%, 30%), with tissue held for one week at each step. Tissue was handled in accordance with ethical guidelines and regulations for the research use of human brain tissue set forth by the NIH (http://bioethics.od.nih.gov/humantissue.html) and the principles of the World Medical Association Declaration of Helsinki (https://www.wma.net/policies-post/wma-declaration-of-helsinki/). All experiments using macaque were carried out in accordance with protocols approved by Yale University’s Committee on Animal Research and NIH guidelines.

### Mice used in this study

All experiments involving mice (*Mus musculus*) were conducted under protocols approved by Yale University’s Institutional Animal Care and Use Committee and in accordance with National Institutes of Health (NIH) guidelines. Mice were housed under controlled environmental conditions (25 °C, 56% relative humidity, 12-h light–dark cycle) with ad libitum access to food and water. Experimental cohorts included both sexes. The day of vaginal plug detection was designated as embryonic day 0.5 (E0.5), and the day of birth as PD 0. The following mouse lines were used: CD-1, C57BL/6J, *Rarb*-null^18^, *Rxrg*-null^18^, RARE-lacZ (Tg(RARE-Hspa1b/lacZ)12Jrt; Jackson Laboratory), and Neurod6-Cre (Nex1-Cre)^60^.

### Generation of *Meis2*-flox line

Mice carrying a conditional floxed *Meis2* allele were generated by CRISPR–Cas9–mediated gene editing, following previously described methods^61,62^. Cas9 target (protospacer) sequences in introns 2 and 3 of the *Meis2* gene were determined by using the MIT CRISPR tool (http://crispr.mit.edu), and the loxP sites flanking exon 3 were inserted (**Extended Data Fig. 11a,b)**. Single-guide RNAs (sgRNAs) targeting these protospacers were transcribed *in vitro* and purified using the MEGAShortscript kit (Invitrogen), and the MEGAclear kit (Invitrogen), respectively. Single-stranded oligodeoxynucleotide (ssODN) repair templates containing loxP sites were synthesized by IDT Technologies. The floxed allele was generated in two steps: first by introducing the 5′ loxP site, followed by breeding and subsequent targeting of the 3′ loxP site. sgRNA–Cas9 ribonucleoproteins (RNPs) and the corresponding ssODN repair template were electroporated into C57BL/6J (Jackson Laboratory) zygotes^62^. Embryos were then transferred into the oviducts of pseudopregnant CD-1 foster females using standard methods. Founder animals were identified by PCR and sequencing of the targeted loxP sites. Correct targeting and germline transmission of the conditional allele were confirmed by breeding with C57BL/6J mice. Genotyping was performed by PCR using the following primers: Forward: 5′-CTCGGCTGATTGAGGGTGTAGTG-3′, Reverse: 5′-AGAGACACACGCACGGAGATG-3′.

### Mouse tissue immunohistochemistry

For mouse brain staining, animals were perfused transcardially with 10 mL 1× DPBS, followed by 10 mL 1× DPBS containing 4% PFA. Brains were post-fixed overnight at 4 °C in 4% PFA in 1× DPBS and subsequently cryoprotected in an ascending sucrose gradient (10%, 20%, 30%) at 4 °C for one week at each concentration, until equilibrated. Tissue was embedded in optimal cutting temperature (OCT) compound (Fisher, 23-730-571), frozen, and sectioned at 20–40 µm on a Leica cryostat (CM3050S), with section thickness dependent on developmental stage (20 µm for PCD 13–PCD 18, 30 µm for PD 0–PD 7, and 40 µm for PD 30–adult). Sections were washed in 1× DPBS at room temperature (3 × 5 min) to remove OCT and permeabilized in 1× DPBS containing 0.6% Triton X-100 for 1 h. Blocking was performed in 1× DPBS containing 5% normal donkey serum and 0.3% Triton X-100 for 1 h at room temperature (RT). Sections were incubated with primary antibodies at 4 °C overnight, washed (3 × 5 min) in 1× DPBS containing 0.3% Triton X-100, and incubated with secondary antibodies (Jackson Laboratory; 1:1000) together with DAPI (1:10,000; Invitrogen) for 2 h at room temperature, followed by washes (3 × 10 min). Sections were mounted on Superfrost Plus slides (Fisherbrand, 22-037-246) and coverslipped with Fluoromount-G (Invitrogen, 00-4958-02). Primary antibodies included: MEIS2 (1:1000, Santa Cruz Biotechnology, sc-81986), BCL11B (1:1000, Sigma-Aldrich, MABE1045), SATB2 (1:1000, Abcam, ab92446), PLXNC1 (1:250, R&D Systems, AF5375), SEMA7A (1:250, R&D Systems, AF1835), RORB (1:500, Novus Biologicals, NBP2-45610), BHLHE22 (1:1000, Sigma-Aldrich, HPA064872), VGLUT2 (1:500, Synaptic Systems, 135404), L1CAM (1:500, Millipore Sigma, MAB5272), tdTomato (1:2000, Scigen, AB8181), ALDH1A3 (1:500, Proteintech, 25167-1-AP; 1:500, Abcam, ab308527), FOXP2 (1:1000, Sigma-Aldrich, MABE415), GFP (1:2000, Aves Labs, NC9510598), NTNG1 (1:100, R&D Systems, AF1166), CASPASE3 (1:500, Cell Signaling Technology, 9661S), IBA1 (1:500, Cell Signaling Technology, 79394SF), and NeuN (1:500, Invitrogen, 702022). Because SEMA7A and PLXNC1 antibodies were raised in sheep and goat, respectively, they could not be simultaneously detected with conventional secondary antibodies due to cross-reactivity between goat and sheep IgGs. To enable co-staining, SEMA7A was conjugated to Alexa Fluor 647 (Invitrogen, A2186) and PLXNC1 to Alexa Fluor 568 (Invitrogen, A2184) using antibody conjugation kits. Conjugated antibodies were used in place of unconjugated primaries, and the secondary antibody step was omitted.

### EdU administration and detection

To label newly generated cortical neurons, pups received an intraperitoneal injection of EdU at 50 mg/kg on PCD 16. EdU solution was prepared from the Click-iT™ EdU Cell Proliferation Kit for Imaging, Alexa Fluor™ 488 dye (Thermo Fisher, C10337) in accordance with the manufacturer’s instructions. Brains were collected at PD 7 and processed for histological analyses. For co-labeling EdU with antibody markers, IHC was performed prior to EdU detection, as EdU Click-iT chemistry is incompatible with several antibody epitopes. Briefly, sections were permeabilized and blocked, followed by incubation with primary and fluorophore-conjugated secondary antibodies according to standard procedures. After completing all antibody labeling steps, EdU incorporation was detected using the Click-iT reaction cocktail from the C10337 kit, following the manufacturer’s protocol. Sections were counterstained with DAPI and mounted in antifade medium.

### Human and macaque tissue immunohistochemistry

Fixed frozen sections were equilibrated to RT and washed in 1× PBS for 10 min. Sections were refixed with 1.6% PFA for 10 min at RT, baked at 60 °C for 30 min to improve tissue adherence, and cooled to RT. Samples were then incubated in acetone for 10 min at RT, washed three times in PBS, and subjected to antigen retrieval in sodium citrate buffer (10 mM citric acid monohydrate, 0.05% Tween-20, pH 6.0) by microwave heating to boiling, followed by cooling to RT. After washing, slides were incubated in autofluorescence-quenching buffer (2.25% H₂O₂ and 10 mM NaOH in PBS) for 90 min at 4 °C under a broad-spectrum LED light source for additional photobleaching. Sections were washed in PBS and blocked in buffer containing 5% donkey serum and 1% BSA diluted in staining buffer (2.5 mM EDTA, pH 8.0; 0.5× PBS; 0.25% BSA; 0.01% NaN₃; 0.122 M Na₂HPO₄; 0.078 M NaH₂PO₄ in ddH₂O) for 45 min at RT. Primary antibodies diluted in blocking buffer were applied overnight at 4 °C. Slides were washed with staining buffer and post-fixed with 1.6% PFA for 10 min at RT, followed by a 5 min incubation in ice-cold methanol at 4 °C. After washing in PBST (0.1% Tween-20 in PBS), secondary antibodies (Jackson Laboratory; 1:500 in 5% donkey serum, 1% BSA in PBST) were applied for 2 h at RT. Nuclei were stained with DAPI for 10 min at RT, followed by two washes in PBST and a final wash in PBS. Sections were mounted with Fluoromount-G (SouthernBiotech, 0100-01). Primary antibodies included: MEIS2 (1:500, Santa Cruz Biotechnology, sc-81986), RARA (Abcam, ab41934), RARB (Proteintech, 14013-1-AP), RXRG (ABclonal, A1877), SATB2 (1:1000, Abcam, ab92446), ALDH1A3 (1:250, Proteintech, 25167-1-AP), and ALDH1A3 (1:500, Abcam, ab308527). Unless otherwise specified, microscopy images were acquired with a 10× objective on an Olympus VS-200 Slide Scanner. For immunohistochemistry quantification, cortical regions were identified and named according to the Allen Brain Atlas. Cortical layers were identified using DAPI or cortical layer markers (SATB2, BCL11B, FOXP2). Fluorescence intensity of IHC-positive signals was quantified using OlyVIA software to measure regional fluorescence intensity. For each cortical region, fluorescence intensity was obtained by averaging three subregions within the area. The number of IHC-positive cells was quantified using the Count Tool in Photoshop for cell counting. For each cortical region, cell numbers were obtained by averaging three subregions within the area. Region selection or division into equal bins were performed following guidelines in Photoshop.

### Section and whole-mount in situ hybridization

Section and whole-mount *in situ* hybridization using antisense digoxigenin (DIG)-labelled RNA probes were performed as described previously^18^, with the slight modification of adding 5% dextran sulfate to the hybridization buffer for whole-mount experiments. Embryonic mouse brains were fixed in 4% PFA overnight at 4 °C, cryosectioned at 20 µm, and stored at −80 °C until use. Commercially available cDNAs for riboprobe synthesis included mouse: *Cbln2* (Horizon Discovery; MMM1013-202798518; Clone Id: 6412317; Acc # BC055682); *Cyp26b1* (Horizon Discovery; MMM1013-202798233; Clone Id: 6400154; Acc # BC059246); *Etv5* (Horizon Discovery; MMM1013-202764508; Clone Id: 4036564; Acc # BC034680); *Bhlhe22* (Horizon Discovery; MMM1013-202797810; Clone Id: 5686844; Acc # BC053007). The *Plxnc1* probe was generated from the *Plxnc1* cDNA amplified using the following primers: forward 5’-CAGCCAATCAAACCTTGAGCAC-3’ and reverse 5’-GTTGTTGAATAGAGGCCCAGTGAC-3’. Mouse *Meis2* cDNA was kindly provided by Dr. John L. R. Rubenstein (University of California, San Francisco). Images of brain sections were acquired on a VS200 microscope (Olympus). Whole-mount *in situ* hybridization samples were imaged, and color balance was manually adjusted to normalize background hue across images without altering signal intensity. *Meis2* intensity in the mPFC was quantified using Photoshop.

### β-Galactosidase histochemical staining

Brains were dissected from PD 0 RARE-lacZ mouse pups and fixed in 4% PFA for 2 h at 4 °C, followed by embedding in OCT compound (Thermo Fisher Scientific, 23-730-572). Frozen brains were sectioned at 20 µm on a Leica cryostat (CM3050S). β-Galactosidase staining was performed following a published protocol^63^ using Red-gal (Sigma-Aldrich, RES1364C-A102X) as the chromogenic substrate. Staining signal intensity was quantified using Photoshop.

### Plasmid construction

For the construction of expression vectors used for luciferase assays, protein-coding regions of mouse *Rxrg*, *Rarb*, and *Meis2* were PCR-amplified and inserted into the pCAGIG vector (Addgene plasmid #11159). Mouse *Rxrg* (clone ID 30608242) and *Rarb* (clone ID 5707723) were purchased from GE Healthcare. Mouse *Meis2* cDNA was kindly provided by Dr. John L. R. Rubenstein. For the luciferase reporter plasmid, mouse *Meis2* promoter region was PCR-amplified from genomic DNA and inserted into the pGL4.24 vector (cat. no. E8421, Promega). Primers for *Meis2* promoter amplification: forward 5’-GAAAGTGAGCTAGGTTGAAGAGTCC-3’ and reverse 5’-CGAGAAAGAGAGAGAGGGAAAGACA-3’.

### Luciferase assays

Neuro2a mouse neuroblastoma cell line was purchased from ATCC. The cell line was authenticated by morphology or genotyping, and no commonly misidentified lines were used. All lines tested negative for mycoplasma contamination, checked monthly using the MycoAlert Mycoplasma Detection Kit (Lonza). Neuro2a cells were transfected using Lipofectamine 2000 (cat. no. 11668019, Thermo Fisher Scientific) with either mouse or human p*CAGIG-Rxrg, Rarb, Meis2,* or empty pCAGIG, together with pGL4.24 luciferase reporter vector carrying *Meis* promoter generated as described above. The Renilla luciferase plasmid (pGL4.73, cat. no. E6911, Promega) was co-transfected to control for transfection efficiency. The luciferase assays were performed 48 h after transfection using the Dual-Luciferase Reporter Assay System (cat. no. E1910, Promega) according to the manufacturer’s instructions. Luciferase activity was measured and quantified by GloMax-Multi Detection System (Promega).

### Anterograde and retrograde tracing in mice

For PD 30 injections, mice were anesthetized according to institutional protocols and positioned in a Kopf stereotaxic instrument (Model 940). The skull was exposed and the bregma point was located, which served as the zero reference point. The mPFC and MD were targeted by performing a craniotomy at the planned injection coordinates. mPFC injections were made at 2.1 mm anterior, 0.3 mm lateral, and 2.4 mm ventral relative to bregma. MD injections were made at 1.2–1.3 mm anterior, 0.40–0.42 mm lateral, and 3.0–3.2 mm ventral relative to bregma. Due to variability in mutant animals, targeting was guided by anatomical landmarks, and only confirmed injections were included in the analysis. Target regions were injected with 40 nL of anterograde AAV carrying p*CAG-tdTomato* (Addgene, 59462-AAV9), retrograde AAV carrying p*CAG-tdTomato* (Addgene, 59462-AAVrg), or retrograde AAV carrying p*CAG-Gfp* (Addgene, 37825-AAVrg). For spinal cord injections, the cervical spinal cord of PD 30 and adult mice was surgically exposed, and 500–1000 nL of retrograde AAV carrying p*CAG-tdTomato* (Addgene, 59462-AAVrg) was injected into the corticospinal tract at C3 (0.2 mm mediolateral; 0.5 mm dorsoventral), covering the dorsal funiculus at this level. The needle was withdrawn after 5 min, the incision sutured, and mice were recovered on a heating pad.

Injection site validation followed a two-step procedure: (1) visual inspection under a dissecting microscope during sample collection for low-magnification screening, and (2) microscopy-based confirmation of injection sites in two- or three-dimensional histological preparations. Following incubation, brains were collected and processed for histology as described in mouse-tissue immunohistochemistry. Three representative sections from each anterior–posterior level were imaged on an Olympus VS200 slide scanner. All anterograde and retrograde tracing samples from control and *Meis2* cKO mice were imaged using identical acquisition parameters. Anterograde tracing was quantified using OlyVIA software to measure tdTomato⁺ regional fluorescence intensity. Retrograde tracing was quantified with the Count Tool in Photoshop to determine the number of GFP⁺ cells. For anatomical analysis, mouse brain sections were subdivided into equal bins using guidelines in Photoshop, and regions were delineated and named according to the Allen Brain Atlas.

### DiI tracing

Brains were dissected at PD 30, and fixed in 4% PFA overnight at 4 °C. The following day, brains were cut sagittally in half to expose the mPFC and thalamus regions. DiI (1,1-Dioctadecyl-3,3,3,3-tetramethylindocarbocyanine) crystals less than 100 mm in diameter were placed just below the surface of mPFC and MD regions. Brains were then embedded in 4% low-melt agarose (Invitrogen) and incubated at 37 °C in 4% PFA for three weeks to allow propagation of the dye along neural projections. Brains were then sectioned at 80 µm using a Leica vibratome and incubated in DAPI (1:10,000 in PBS) for 10 min at room temperature before being washed in PBS. Sections were mounted on glass slides, sealed with Fluoromount-G, and imaged on the same day. For the reconstruction of scanned images into a three-dimensional volume, Serial Section Assembler function of NeuroInfo (ver. 2024.1.1, MBF Bioscience) was used.

### Human telencephalic organoid culture

Human induced pluripotent stem cell line YB7-GFP was authenticated by morphology or genotyping and confirmed to be free of mycoplasma contamination using the MycoAlert Mycoplasma Detection Kit (Lonza). For maintenance of pluripotency, cells were dissociated into single cells with Accutase (Thermo Fisher Scientific, 00-4555-56) and plated at a density of 1 × 10⁵ cells per cm² on Matrigel-coated 6-well plates (Falcon) in mTeSR1 medium (STEMCELL Technologies, 85850) supplemented with 5 µM Y-27632 (Sigma-Aldrich, SCM075). ROCK inhibitor was removed after 24 h, and cells were maintained for an additional 4 days before passaging. Telencephalic organoids were generated following a directed differentiation protocol^18^. Briefly, cells were dissociated with Accutase and resuspended in neural induction medium containing 100 nM LDN193189 (STEMCELL Technologies, 72147), 10 µM SB431542 (Selleck Chemicals, S1067), and 2 µM XAV939 (Sigma-Aldrich, X3004-5MG) to achieve dual SMAD and WNT inhibition. Cells (10,000 per well) were plated in 96-well v-bottom ultra-low-attachment plates (Sumitomo Bakelite). To promote survival and aggregation, 10 µM Y-27632 was added for the first 24 h. After 10 days in static culture, organoids were transferred to 6-well ultra-low-attachment plates (Millipore Sigma) and maintained on an orbital shaker at 90 rpm. Beginning on day 18, organoids were cultured in maturation medium supplemented with 1× CD lipid concentrate (Thermo Fisher Scientific, 11905031), 5 µg ml⁻¹ heparin (STEMCELL Technologies, 07980), 20 ng ml⁻¹ BDNF (Abcam, 9794), 20 ng ml⁻¹ GDNF (R&D Systems, 212-GD), 200 µM cAMP (Sigma-Aldrich, 20-198), and 200 µM ascorbic acid (Sigma-Aldrich, A92902). On day 148, all-trans retinoic acid (Sigma-Aldrich, R2625) was added for 48 h prior to sample collection.

For histological preparation, organoids were fixed in 4% PFA at 4 °C, cryoprotected in 20% sucrose, and embedded in OCT compound (Thermo Fisher Scientific, 23-730-572). Sections (10 µm) were cut on a Leica cryostat (CM3050S), washed in PBS (3 × 5 min), and blocked in PBS containing 0.5% Triton X-100 and 10% donkey serum (Jackson ImmunoResearch Laboratories, 017-000-121) for 2 h at room temperature. Sections were incubated with primary antibodies diluted in blocking buffer overnight at 4 °C, washed (3 × 5 min), and incubated with fluorescent secondary antibodies in 10% donkey serum for 2 h at room temperature. After a final PBS wash (3 × 5 min), sections were coverslipped with Vectashield mounting medium (Vector Laboratories, H-1000). Primary antibodies included MEIS2 (1:500, Santa Cruz Biotechnology, sc-81986), ALDH1A3 (1:250, Proteintech, 25167-1-AP; 1:500, Abcam, ab308527) and NeuN (1:500, Invitrogen, 702022). Images were acquired on a Zeiss LSM800 confocal microscope and processed with ZEN (Zeiss) and ImageJ software. Z-stack images were analyzed using Volocity (v6.3.1) and Spotfire (v11.2.0).

### Mouse tissue collection and nuclei Isolation for mouse sn-multiome data

mPFC and MC tissues were dissected from P0 wild-type mouse brains under RNase-free conditions. All procedures were performed on ice to preserve RNA integrity. Dissected tissues were immediately transferred to pre-chilled 1.5 mL tubes and processed individually for single-nucleus Multiome profiling. Nuclei isolation was performed following a modified 10x Genomics protocol optimized for mouse cortical tissue. Briefly, tissue samples were homogenized in 1 mL of ice-cold lysis buffer using a Dounce homogenizer (30 strokes with a loose pestle followed by 30 strokes with a tight pestle). To minimize air bubble formation, the pestle was not lifted above the buffer surface during homogenization. The lysate was filtered through a pre-wetted 40 μm cell strainer into a 50 mL conical tube, followed by the addition of 3 mL iodixanol. After gentle inversion, samples were centrifuged at 1000 x g for 30 min at 4 °C using a swing-bucket rotor. Following centrifugation, the supernatant was carefully aspirated, and the nuclear pellet was sequentially resuspended, each followed by 5 min incubation on ice. The resuspended nuclei were filtered through a 35 μm strainer, pelleted again at 500 x g for 10 min at 4 °C, and resuspended in 10x Genomics Nuclei Buffer supplemented with RNase inhibitor and DTT. The final nuclei concentration was adjusted to 3.3-4.0 x 10⁶ nuclei/mL based on manual hemocytometer counts. All buffers were freshly prepared and maintained ice-cold throughout the procedure.

### Mouse sn-Multiome library preparation

Nuclei were processed immediately after quantification using the 10x Genomics Chromium Next GEM Single Cell Multiome ATAC + Gene Expression v1.1 kit according to the manufacturer’s protocol. Reagents were thawed and equilibrated following 10x Genomics guidelines: ATAC Buffer and 20X Nuclei Buffer were thawed to room temperature, while ATAC Enzyme B and Barcoding Enzyme Mix were kept on ice. For each reaction, 10 μL of Transposition Mix (7 μL ATAC Buffer B + 3 μL ATAC Enzyme B) was prepared on ice. Nuclei were added to the Transposition Mix and incubated at 37 °C for 1 h, followed by immediate cooling to 4 °C. Subsequent GEM generation and barcoding were carried out on the Chromium Controller (Chip J). Each lane contained ∼10,000 nuclei mixed with 60 μL Master Mix (Reducing Agent B, Template Switch Oligo, Barcoding Reagent Mix, and Barcoding Enzyme Mix). Following droplet encapsulation, GEM-RT incubation was performed at 37 °C for 45 min and 25 °C for 30 min. GEMs were then quenched with 5 μL Quenching Agent and gently mixed. Resulting emulsions were either processed immediately or stored at −80 °C for up to 4 weeks before downstream library construction.

### Sequencing and data preprocessing for mouse sn-multiome data

Single-nucleus RNA-seq and ATAC-seq libraries were generated using the 10x Genomics Chromium Next GEM Single Cell Multiome ATAC + Gene Expression kit following the manufacturer’s protocol. Libraries were sequenced on an Illumina NovaSeq 6000 platform to achieve a target depth of ∼25,000 paired end reads per nucleus for RNA and ∼20,000 reads per nucleus for ATAC. Raw sequencing data were processed using Cell Ranger ARC (v2.0.1, 10X Genomics) with default parameters. The mouse reference genome mm10 was used for alignment and annotation. Separate outputs for snRNA-seq and snATAC-seq modalities were generated and stored as count matrices and fragment files, respectively. For quality control, nuclei were filtered based on multiple metrics: 1) RNA modality: nCount_RNA < 25,000, nFeature_RNA > 500, and mitochondrial percentage < 5 %. 2) ATAC modality: TSS enrichment score > 5 and nucleosome signal < 2. Potential doublets were identified and removed using scDblFinder (v1.14) based on gene expression profiles. After QC filtering and doublet removal, cell-type annotation was performed using only and snRNA-seq modality prior to integration with ATAC data, since RNA counts provided higher resolution and coverage across cell types compared to ATAC profiles. Cell-type identities were assigned by comparing gene expression patterns to canonical mouse cortical markers, including *Bcl11b* (CThPN), *Satb2* (upper-layerCPN), *Fezf2* (L6, CThPN), *Gad1* and *Slc32a1* (InterN), and *Mog* (oligodendrocyte lineage). Raw sequencing data have been deposited at GEO under accession number GSEXXXXXX.

### Human tissue dissection and processing

Donor ages spanned the mid-fetal window as defined in our previous work^16^. Post-conception age was calculated as gestational age in weeks minus two weeks. The postmortem interval (PMI) was defined as the time (hours) between death and freezing of the tissue. Tissue dissections followed established protocols^16^. The dlPFC was identified using known anatomical landmarks. Samples from 12–24 post-conception weeks primarily comprised cortical plate and, in some cases, also included the underlying subplate.

### Human tissue nuclear isolation and multiome capture

Four human postmortem brain specimens were used for the 10x Genomics multiome assay (**Supplementary Table 1**). Nuclei were isolated following established protocols^64,65^. To minimize experimental bias, frozen brain tissue was pulverized in liquid nitrogen to a fine powder using a mortar and pestle (Coorstek, #60316 and #60317) when the dlPFC region weighed more than 30 mg. Donor tissue was aliquoted into 1.5 mL tubes in portions of 10–30 mg. All buffers were prepared using molecular-grade reagents unless otherwise specified and were maintained at 4 °C until use. 1 mL of lysis buffer (250 mM Sucrose (Sigma, cat# S0389), 25 mM KCl (Sigma, cat#60142), 5 mM MgCl2 (Sigma, cat# M1028),20 mM Tris-HCl pH 7.5 (Invitrogen, cat#15567-027), 0.1% Igepal (Sigma, cat# I8896), 1X protease inhibitor (Roche, cat#11836170001 or cat#5056489001), 1 mM DTT (Sigma, cat# 43816), 0.08 U/ul RNase Inhibitor (Roche, cat# 3335402001) was added directly to each tissue aliquot and vortexed. The suspension was transferred to a pre-chilled dounce homogenizer on wet ice containing an additional 1 mL of lysis buffer. To recover residual tissue, another 1 mL of lysis buffer was added to the aliquot tube, vortexed, and combined with the homogenizer. Tissue was homogenized using loose and tight pestles, 30 strokes each, with constant pressure and without introducing air bubbles. The homogenate was filtered through a 40 µm cell strainer (Corning, 352340) pre-wetted with 250 µL of lysis buffer. 3ml of iodixanol gradient buffer (25 mM KCl, 5 mM MgCl2, 20 mM Tris-HCl pH 7.5, 50% Iodixanol (Sigma, cat# D1556), 1% BSA + 10x Protease Inhibitor (premade stock solution), 1 mM DTT, 0.08 U/ul RNase Inhibitor) was added to the filtered homogenate, mixed by inversion 10x, and centrifuged at 1000g for 30 minutes in a swinging bucket at 4C. Following centrifugation, samples were removed, and debris and supernatant was carefully removed. The pellet was resuspended using 0.5ml of nuclear permeabilization buffer (10 mM NaCl, 3 mM MgCl2, 10 mM Tris-HCl, pH 7.5, 0.01% NP-40 substitute (Sigma, cat# 74385), 0.01% Tween-20 (BioRad, cat# 166-2404), 0.001% Digitonin (Thermofisher, cat#BN2006), 1X protease inhibitor, 1mM DTT, 1% BSA, 1U/ul Rnase Inhibitor) and incubated on ice for 5 minutes. Subsequently, 0.5ml of wash buffer: 10mM NaCl, 3 mM MgCl2, 10 mM Tris-HCl, pH 7.5, 0.1 % Tween-20, 1 U/ul Rnase Inhibitor, 1 mM DTT, 1 % BSA, was added and the nuclear suspension was filtered through a 30um cell strainer into a new 1.5ml tube. A small aliquot of nuclear suspension was removed for counting. The rest of the sample was centrifuged at 500g for 10mins at 4C. During this time, samples were counted (1:2 dilution for postnatal samples and a 1:4 dilution for prenatal samples) on a manual hemocytometer at 20x magnification. The nuclear suspension was spun down at 500g for 10min at 4C. Based on the calculated concentration, samples were resuspended in resuspension solution (1x Nuclei Buffer from 10x Genomics ATAC Kit A, 1 mM DTT, 1 U/ul RNase Inhibitor) in a volume that would result in approximately 8-10 million nuclear per ml. After resuspension, samples were further filtered through a 40um cell strainer. Another aliquot of each sample was removed for a second round of counting to verify that the concentration of the sample fell between 3.3 and 8 million nuclei/ml, as per the 10x Genomics GEX and ATAC manual (CG000338) for 10,000 nuclei targeted capture. Single nuclei were captured according to the 10x Genomics Multiome protocol. Gene expression (GEX) and ATAC libraries were prepared on the 10x Genomics Multiome platform and sequenced on a NovaSeq 6000 (Illumina) at the Yale Center for Genomic Analysis (YGCA) to an initial depth of 250 million reads per library. Libraries flagged during quality control were re-sequenced to achieve a minimum of 25,000 reads per nucleus. Raw data are available at GEO under accession number GSEXXXXXX.

### Human sn-multiome data alignment and processing

Sequencing data were processed using 10x Genomics Cell Ranger pipelines. Multiome libraries were first run with cellranger-ARC (v2.0.1). To ensure optimal recovery of high-quality cells from each modality, RNA and ATAC data were subsequently processed separately: RNA libraries with cellranger (v6.0.1) and ATAC libraries with cellranger-ATAC (v2.0.0). For RNA, cell calling followed the barcode rank plot–based method implemented in Cell Ranger, whereas ATAC cell calling and library-level quality control were performed using the default cellranger-atac workflow.

snRNA-seq and snATAC-seq data were processed using Seurat (v5.1.0)^66^, Signac (v1.15.0)^67^, and scDblFinder (v1.14) following standardized quality-control and integration procedures. For scRNA-seq, cells were retained only if they exhibited dataset-appropriate UMI and gene counts and contained <5% mitochondrial transcripts; low-quality clusters showing aberrant marker profiles or compromised library complexity were manually reviewed and excluded. Putative doublets were identified with scDblFinder using default settings, and clusters were annotated as doublets when they displayed mixed lineage signatures together with a doublet-score above the scDblFinder-defined 75th percentile, after which these clusters were removed from downstream analyses. Filtered RNA data were normalized using SCTransform with glmGamPoi, integrated across batches using Harmony, and subjected to PCA, UMAP, and graph-based clustering prior to marker-based annotation. scATAC-seq libraries were processed with Signac, and high-quality nuclei were selected based on nFragments > 1,000 and TSS enrichment > 4, with extreme nucleosome signal or blacklist enrichment filtered out. Peaks were called with MACS2 on pseudobulk aggregates, gene-activity scores were computed from promoter-proximal and gene-body–extended peaks, and dimensionality reduction was performed using TF-IDF and SVD followed by Harmony correction. Multi-omic integration was achieved using Seurat’s weighted nearest-neighbor (WNN) framework, aligning transcriptomic and chromatin-accessibility manifolds to generate unified clusters. Discrepant or low-quality multimodal clusters were manually inspected and removed, and final cell-type annotations were assigned based on integrated RNA markers, chromatin accessibility patterns, and regulatory element–to–gene linkage profiles. Cell types were annotated based on canonical marker genes, including caudal ganglionic eminence interneurons (CGE), excitatory newborn/migrating neurons (Ex-Newborn/Migrating), glial intermediate progenitors (gIP), immature layer 2/3 excitatory neurons (Immature L2/3), immature layer 4/5 excitatory neurons (Immature L4/5), layer 5 extratelencephalic projection neurons (L5 ET), layer 5/6 near-projecting neurons (L5/6 NP), layer 6 corticothalamic neurons (L6 CT), layer 6 intratelencephalic neurons (L6 IT), layer 6b neurons (L6b), medial ganglionic eminence interneurons (MGE), microglia (Micro), neuronal intermediate progenitors (nIP), oligodendrocyte precursor cells (OPC), and radial glia (RG) (**Extended Data Fig. 1b**).

### Regulatory network inference in human mid-fetal dlPFC

To identify TFs regulatory programs, SCENIC (pySCENIC v1.0a2)^68^ was applied to the sn-multiome data. Regulon activity scores were calculated per cell type, and top enriched regulons were identified using the regulon specificity score (RSS). To refine regulatory predictions, TF–target interactions were filtered by paired chromatin accessibility profiles, retaining only motifs accessible in relevant cell populations. A TFs network specific to the dlPFC was constructed by integrating SCENIC-derived regulons with ATAC-supported motif accessibility. TF–TF interactions were extracted and visualized in R (v4.4.1) using the igraph and ggraph packages. Node size was scaled by the number of regulon targets, and clusters were color-coded according to functional categories. The resulting dlPFC TFs regulatory network was visualized in **Extended Data Fig. 1d**, and the complete TF–target table was provided as a supplementary dataset (**Supplementary Table 2**). This network formed the basis for constructing the retinoic acid–associated regulatory network.

### Nuclear isolation and CUT&Tag library

Two human postmortem brain specimens (**Supplementary Table 1**) were used for CUT&Tag library preparation. CUT&Tag requires high-quality samples; therefore, freshly harvested or freshly flash frozen tissues are essential for optimal performance. Fresh frozen tissue was subjected to nuclei extraction followed by CUT&Tag library construction. Nuclei were isolated using the Minute™ Single Nucleus Isolation Kit for Neuronal Tissues (Invent Biotechnologies, BN-020) according to the manufacturer’s instructions, except that reagent B (myelin removal) was omitted, as myelin is sparse at mid-fetal stages and its use result in substantial nuclear loss. Pelleted nuclei were resuspended in 5% BSA and stained 1:1 with trypan blue (Invitrogen, T10282). Nuclei excluding trypan blue were were quantified using a LUNA-II™ Automated Cell Counter.

For CUT&Tag-seq (Hyperactive Universal CUT&Tag Assay Kit for Illumina, Vazyme, TD903), the basic procedure was performed following previously described methods^69^. Specifically, the protocol proceeded as follows: 1×10⁵–2×10⁵ nuclei were washed with 100 µL wash buffer and centrifuged at 600g for 3 min at room temperature. Cell pellets were resuspended in 100 µL wash buffer. Concanavalin A–coated magnetic beads were washed twice with 100 µL binding buffer, and 10 µL of activated beads was added to the suspension and incubated for 10 min at room temperature. After incubation, the buffer was removed and bead-bound nuclei were resuspended in 50 µL antibody buffer. Primary antibodies (1 µg per 1×10⁵ nuclei) were added and incubated overnight at 4 °C. The following antibodies were used: RARA (Abcam, ab41934)^70–73^, RARB (Invitrogen, PA1-811)^74^, RXRG (ABclonal, A1877), and MEIS2 (Abcam, ab244267); H3K4me3 (ABclonal, A22146) and H3K27ac (ABclonal, A22396) as positive controls; rabbit IgG (EpiCypher, 13-0042) and mouse IgG (Diagenode, C15400001-15) served as negative controls. After removing the primary antibody solution, 0.5 µg of the appropriate secondary antibody (EpiCypher, 13-0047 and 13-0048) was added in 50 µL Dig-wash buffer and incubated for 1 h at room temperature. Cells were washed three times with Dig-wash buffer to remove unbound antibodies. Cells were then incubated with 0.04 µM Tn5 transposase in 100 µL Dig-300 buffer for 1 h at room temperature, followed by three washes with Dig-300 buffer to remove excess transposase. Bead-bound nuclei were resuspended in 50 µL tagmentation buffer and incubated for 1 h at 37 °C. Tagmentation was terminated by adding 5 µL of 20 mg/mL Proteinase K, 100 µL Buffer L/B, and 20 µL DNA Extract Beads, followed by incubation for 10 min at 55 °C. DNA was purified using DNA Extract Beads. PCR amplification was used to enrich Tn5-fragmented DNA and to incorporate i5/i7 adapters (TruePrep Index Kit V2 for Illumina, Vazyme, TD202), sequencing primers, and sample indices for Illumina sequencing. Amplified products were purified with 1.2× SPRIselect reagent (Beckman Coulter, B23318) and subjected to Illumina PE150 Nova sequencing at the YGCA. Raw data are available at GEO under accession number GSEXXXXXX.

### CUT&Tag data processing

Raw FastQ files were processed with Trim Galore to remove low-quality bases and adapter sequences. Cleaned reads were aligned to the human reference genome (GRCh38/hg38) using Subread (v2.0.1)^75^. PCR duplicates, unpaired reads, and low-quality alignments were removed using Sambamba (v0.6.6)^76^ and SAMtools (v1.21)^77^. The resulting BAM files were converted into RPKM-normalized bigWig tracks with deepTools (v3.5.6)^78^ and visualized in IGV (v2.16.2)^79^ using the “autoscale” option to enable cross-sample comparison. For downstream visualization, bigWig files from biological replicates were merged, and peaks bound by RARA, RARB, and RXRG were combined into a unified RAR/RXR set. Peak calling for H3K4me3, H3K27ac, RAR/RXR, and MEIS2 CUT&Tag data was performed using three independent algorithms: MACS2 (v2.2.9.1)^80^, HOMER (v4.11)^81^, and LANCEotron (v1.2.7)^82^, followed by reproducibility assessment with IDR (v2.0.4.2). Corresponding consensus peak sets were merged using the following criteria: (1) for biological replicates, only peaks with >50% reciprocal overlap were retained; and (2) across algorithms, only peaks identified by at least two methods were retained (**Supplementary Table 3; Supplementary Table 10**).

To quantify the enrichment of each chromatin mark, read counts were extracted from the aligned BAM files using bedtools multicov. For each histone modification or TFs dataset, biological replicates were jointly processed with the corresponding consensus peak sets, generating a peak-count matrix. The resulting count tables were imported into R for normalization using the edgeR package. Normalization factors were computed with the trimmed mean of M values (TMM) method, and normalized expression values were expressed as counts per million (CPM) and log₂-transformed CPM (logCPM). The normalized matrices contained both raw read counts and normalized CPM/logCPM values for each biological replicate. To avoid bias introduced by arbitrary abundance cutoffs, all peaks were retained for downstream analyses, although CPM- or logCPM-based filtering can optionally be applied to remove low-abundance regions. For peak annotation, two complementary approaches were applied: ChIPseeker (v1.40.0)^83^, which favors proximal annotation, and rGREAT (v2.6.0)^84^, which favors distal annotation. All genes assigned to annotated peaks were used for genome-wide analyses, including enrichment and network construction. For specific loci of interest, peak assignments were manually inspected to ensure accuracy. To improve GO enrichment specificity, we excluded entries corresponding to non-coding or poorly characterized loci. Specifically, annotated genes containing the terms *“microRNA”*, *“non-protein coding”*, or *“uncharacterized”* in the GENENAME field were removed prior to downstream analyses. Filtering was performed in R using case-insensitive regular expressions, and only protein-coding entries were retained for the identification of downstream targets overlapping with RAR/RXR binding sites. De novo motif enrichment analysis of the merged peak sets was performed using HOMER to identify potential TFs binding motifs.

### Human mid-fetal PFC-enriched gene analysis

Independent spatiotemporal human brain exon microarray and RNA-seq datasets were obtained from BrainSpan^37^. For the early and late mid-fetal periods^16^, a total of 105 mRNA samples corresponding to 11 prospective neocortical areas—including the pial surface, marginal zone, cortical plate (layers 2–6), and adjacent subplate zone—from developmental windows 2 and 4 (post-conception weeks 13–22) were analyzed. The neocortical regions included the orbital (oPFC or OFC), dorsolateral (dlPFC or DFC), ventrolateral (vlPFC or VFC), and medial (mPFC or MFC) prefrontal cortex, as well as the primary motor cortex (M1C) from the frontal lobe; the primary somatosensory cortex (S1C) and posterior inferior parietal cortex (IPC) from the parietal lobe; the primary auditory cortex (A1C), posterior superior temporal cortex (STC), and inferior temporal cortex (ITC) from the temporal lobe; and the primary visual cortex (V1C) from the occipital lobe. To identify PFC-enriched genes, neocortical areas were divided into two groups: the prefrontal cortex (PFC; dlPFC, mPFC, oPFC, vPFC) and non-prefrontal cortex (non-PFC; M1C, S1C, IPC, A1C, V1C, STC, ITC). Raw gene-level counts were analyzed using edgeR^85^. Lowly expressed genes were filtered using *filterByExpr*, and library sizes were normalized with the TMM method. A design matrix with the group factor (PFC versus non-PFC) was fitted using the quasi-likelihood pipeline (*estimateDisp*, *glmQLFit*, *glmQLFTest*). P values were adjusted by the Benjamini–Hochberg false discovery rate (FDR), and differentially expressed genes (DEGs) were defined at FDR < 0.05 (optionally with |log₂FC| > 0.58) (**Supplementary Table 4**). Log₂ fold-change values (PFC versus non-PFC) and FDR estimates from the edgeR analysis were incorporated into the dlPFC regulatory network as node-level annotations, providing an expression-based measure of PFC enrichment for each gene. In addition, only protein-coding genes were retained for bar plot visualization (**Extended Data Fig. 3a**), with non-coding or poorly characterized transcripts excluded.

### Construction of the retinoic acid regulatory network

The RA-GRN was constructed by integrating multi-omics datasets. First, CUT&Tag profiling of RA receptors was performed to identify downstream regulatory targets. Second, the human dlPFC regulatory network was inferred using pySCENIC, retaining only genes associated with regulatory peaks, which served as the framework for the regulatory architecture. The intersection of SCENIC-predicted regulons with CUT&Tag-identified RA receptor targets was used to define high-confidence RA regulatory interactions. To annotate network nodes, FDR and log₂ fold-change values (PFC versus non-PFC) from edgeR analysis were incorporated as expression-based features. Node degree, calculated in R (igraph package), was used to represent regulatory connectivity. The integrated RA regulatory network was visualized in R using igraph and ggraph, and the complete set of regulatory interactions is provided in **Supplementary Table 5**.

### Mouse bulk RNA-seq library and data processing

PD 0 pups were sacrificed, and brains were rapidly dissected into fresh ice-cold Hanks’ balanced salt solution (Gibco, 14175-095). The mPFC was minced and digested for 15–20 min at 37 °C in DMEM (Gibco, 10566024) containing 80 U ml⁻¹ papain (Sigma-Aldrich, P3125) and 200 U ml⁻¹ DNase I (Invitrogen, 18047019). Tissue was dissociated into single-cell suspension by gentle pipetting, and the reaction was quenched with DMEM supplemented with 10% FBS. Cells were passed through a pre-wetted 40 µm cell strainer (Corning, 352340) and pelleted at 300g for 5 min. The pellet was resuspended in DMEM containing 10% FBS, and viable cells were quantified using 0.4% trypan blue (Invitrogen, T10282) and a LUNA-II™ Automated Cell Counter.

RNA-seq libraries were prepared from 20,000–50,000 dissociated mPFC cells using the VAHTS Universal V10 RNA-seq Library Prep Kit for Illumina (Vazyme, NR606-01), incorporating i5/i7 adapters from the VAHTS Multiplex Oligos Set 4/5 (Vazyme, N322-01). Amplified libraries were purified with SPRIselect reagent (Beckman Coulter) and sequenced on an Illumina NovaSeq 6000 platform (PE150) at the YCGA. Raw sequencing data have been deposited at GEO under accession number GSEXXXXXX.

FASTQ files were processed with Trim Galore to remove low-quality bases and adapter sequences. Clean reads were aligned to the mouse reference genome (GRCm38/mm10) using Subread. Gene-level counts were obtained with *featureCounts*, and differential expression analysis (*Meis2* cKO versus control) was performed using DESeq2, with differentially expressed genes defined at padj < 0.05 (optionally with |log₂FC| > 0.58). Sex-associated genes and ribosomal protein–related genes were excluded when generating volcano plots and bar charts for DEGs (**Supplementary Table 9**).

### Analysis of human mid-fetal brain spatial transcriptomics

Spatial transcriptomic data were obtained from a publicly available resource^48^, accessible at http://donglab.life/brainAtlas.html. For the present analysis, sections corresponding to PCW 14 were selected.

### Comparative transcript analysis of *MEIS2*

To assess the evolutionary conservation of *MEIS2* across species, RefSeq transcript sequences were retrieved using the R package rentrez (v1.2.3). Searches were performed in the NCBI Gene database for *Homo sapiens*, *Pan troglodytes*, *Macaca mulatta*, and *Mus musculus*, and linked RefSeq RNA accessions were obtained. Transcripts annotated as “PREDICTED” or “non-coding” were excluded to retain only validated protein-coding isoforms. For each retained RefSeq accession, the corresponding nucleotide sequence was downloaded in FASTA format and parsed into DNA strings using the Biostrings package (v2.72.0). Sequences were compiled into a DNAStringSet object for multiple sequence alignment using ClustalW as implemented in the R package msa (v1.30.0)^86^. The resulting alignment was converted into a phylogenetic data object (phyDat) using phangorn (v2.11.1)^87^, and pairwise distances were estimated using the maximum-likelihood method (dist.ml). A phylogenetic tree was constructed using the neighbor-joining (NJ) algorithm, and visualized in R (v4.4.1) with annotated branch lengths. Pairwise distance matrices and corresponding similarity matrices were exported for downstream analysis.

### Motif discovery and conservation analysis of *MEIS2*

Position weight matrices (PWMs) for *MEIS2*-associated binding motifs were obtained from MEME-based motif discovery analyses. Sequence logos were generated in R (v4.4.1) using the ggseqlogo package^88^ to visualize base-specific conservation across motif positions. For comparative motif analysis, multiple PWMs were imported as position frequency matrices (PFMs) using the motifStack package^89^ and visualized as stacked logos or radial motif-based trees to assess conservation and divergence across datasets.

### Comparative analysis of MEIS2 protein conservation

Protein sequences of *MEIS2* orthologs from human (*Homo sapiens*), chimpanzee (*Pan troglodytes*), rhesus macaque (*Macaca mulatta*), and mouse (*Mus musculus*) were retrieved from UniProt using the UniProt REST API. Only manually curated, reviewed entries were retained for downstream analysis. Protein sequences were imported into R (v4.4.1) as AAStringSet objects using the Biostrings package (v2.72.0). Multiple sequence alignment (MSA) was performed with the ClustalW algorithm implemented in the msa package (v1.30.0)^86^. The aligned sequences were exported in FASTA format and visualized using ggmsa (v1.1.4)^90^, generating both global alignments and sequence logos highlighting conserved and variable residues. To quantify residue conservation relative to the human sequence, the alignment was converted into position-wise matrices using seqinr (v4.2-36)^91^, and amino acid positions were classified as conserved or variable. Comparative heatmaps were generated with ggplot2 (v3.5.1), in which residues identical to the human sequence were coded as conserved and differences as variable. Protein structural domains of human MEIS2 were annotated from UniProt (MEIS N-terminal domain, DNA-interaction region, and transcriptional activation domain), and mapped alongside sequence variability using patchwork (v1.2.0) for combined visualization of functional domains and residue conservation. For each species, the proportion of conserved amino acids relative to human MEIS2 was quantified, providing a residue-level conservation score. Summary statistics including total aligned sites, number of matched residues, and percent conservation were calculated and visualized.

### Weighted gene co-expression network analysis

Spatiotemporal human brain exon microarray and RNA-seq datasets were obtained from BrainSpan^37^. Genes previously identified as RA-GRN members were extracted, and their expression values across PFC samples were used to construct a spatiotemporal expression matrix spanning fetal through aging developmental periods. Weighted gene co-expression network analysis (WGCNA v1.72-5)^92^ was performed following standard procedures. A soft-thresholding power was selected using the scale-free topology criterion. The resulting adjacency matrix was transformed into a topological overlap matrix (TOM), and genes were hierarchically clustered based on TOM dissimilarity. Co-expression modules were identified using dynamic tree cutting. Module eigengenes were calculated for each module, and their temporal trajectories across developmental stages were evaluated. Modules showing coherent temporal expression patterns were considered candidate regulatory modules associated with RA signaling in the PFC (**Supplementary Table 6**).

### Gene set enrichment analysis of RA-GRN modules using NDD and SCZ database

To assess whether disease-associated genes were enriched within individual modules of the RA-GRN, we performed a hypergeometric test using the set of all module genes as the background. For each module, the number of overlapping genes with the disorder-associated gene sets was compared against the expected overlap by chance, and *P* values were computed using the *phyper()* function in R, followed by multiple-testing correction using the Benjamini–Hochberg method. Modules with FDR < 0.05 were considered significantly enriched and are outlined in green in the figure. The NDD gene set was obtained from the NIMH Neurodevelopmental Disorders priority gene list (https://grants.nih.gov/grants/guide/notice-files/NOT-MH-24-370.html), and the SCZ gene set was derived from a previous study^59^, consisting of all genes located within CNVs associated with schizophrenia (**Supplementary Table 7**).

### Weighted gene set enrichment analysis of RA-GRN modules using SFARI-ASD database

Weighted gene set enrichment analysis was performed to assess the association between co-expression modules and ASD–related genes. Each gene was assigned an ASD relevance score based on the EAGLE score from SFARI databases (https://gene.sfari.org/database/human-gene/) (**Supplementary Table 7**). Genes with missing scores were assigned a value of zero. To ensure consistency, gene symbols were harmonized between the ASD gene list and module annotations, and duplicated entries were removed. A ranked list of all expressed genes was generated based on their ASD relevance score (higher scores indicate stronger ASD association). Co-expression modules identified from transcriptomic data were treated as predefined gene sets, with each module containing all genes assigned to it. Enrichment analysis was conducted using the fgsea R package (v1.30) with 10,000 permutations. The algorithm calculates an enrichment score (ES) for each module, representing the degree to which module genes are overrepresented at the top of the ASD-ranked gene list. To correct for module size and multiple testing, normalized enrichment scores (NES) and Benjamini–Hochberg adjusted FDR values were computed. Modules with FDR < 0.05 were considered significantly associated with ASD. The resulting NES values reflect the direction and magnitude of enrichment—positive NES indicates enrichment among high-confidence ASD genes, while negative NES indicates depletion. Leading-edge genes contributing most to each enrichment signal were extracted from the fgsea output for downstream visualization and functional analysis.

### MAGMA analysis of RA-GRN modules using GWAS data of human psychiatric disorders

MAGMA (v2.9.0)^93^ was used to test whether RA-GRN gene modules were enriched for common variants associated with human psychiatric disorders. GWAS summary statistics were obtained from the Psychiatric Genomics Consortium (PGC; https://pgc.unc.edu/for-researchers/download-results/) for schizophrenia (10.6084/m9.figshare.14681220)^94^, bipolar disorder (10.6084/m9.figshare.14671998)^95^, major depressive disorder (10.6084/m9.figshare.14672085)^96^, and attention-deficit hyperactivity disorder (10.6084/m9.figshare.22564390)^97^. For gene-level analysis, the 1000 Genomes Project European reference panel was used to estimate linkage disequilibrium, applying the SNP-wise mean model. For gene-set analysis, technical confounders including gene size, gene density, mean minor allele count, and their log-transformed values were included as covariates in the regression model. Nominal P values were adjusted for multiple testing using the with Benjamini–Hochberg method. Full results are provided in **Supplementary Table 8**.

### Gene set enrichment analysis

Functional enrichment analyses were performed using clusterProfiler (v4.12.0)^98^ in R. Gene lists derived from human and mouse datasets were used as input. For human data, GO analyses were conducted with the *enrichGO* function and the org.Hs.eg.db annotation database (v3.19.1). For mouse data, enrichment analyses were performed with the corresponding org.Mm.eg.db annotation database (v3.19.1). Enrichment significance was evaluated with the Benjamini–Hochberg method to control the false discovery rate (FDR < 0.05). Results were visualized using the *barplot* function in clusterProfiler.

### Behavioural tests

Male mice aged 8–10 weeks were used for all behavioral experiments. All mouse lines used for behavioral experiments were maintained on a C57BL/6J genetic background. For behavioral testing, mice were group-housed under standard conditions (1–3 mice per cage, same sex per cage) on a 12h light/dark cycle with ad libitum access to food and water. Littermate controls were used whenever feasible; when littermates were not available in sufficient numbers, age-matched C57BL/6J controls from the same colony were used, and all mice were bred and housed under identical conditions. We performed a series of tests to assess cognitive, anxiety-related, motor, and balance behaviours in mice. Mice were acclimatized in the testing room for 30 min before the start of each experiment. The apparatus was sanitized with 70% ethanol before the first trial and with water for subsequent trials. To minimize behavioural alterations caused by ethanol scent, cage dust was rubbed on the apparatus for the initial control trial; data from this trial were excluded from analyses. Behavioural tests were performed in the order described below, with 24h intervals between tests. All experiments were performed during the light phase.

Y-maze. Mice were placed at the centre of the Y-maze and allowed to explore freely for 10 min. Arm entries were recorded using Noldus EthoVision XT software. Spontaneous alternation (%) was calculated as the ratio of successive entries into three different arms divided by the total number of possible alternations (total entries – 2). Reduced alternation was interpreted as impaired cognitive function.

Open-field test. Mice were placed in the centre of a rectangular arena (50 × 50 cm) and allowed to explore freely for 10 min. Movement was recorded using Noldus EthoVision XT software, which automatically tracked distance traveled, velocity, and time spent in centre versus periphery zones. Reduced time in the centre and increased time in the periphery were interpreted as elevated anxiety-like behaviour.

Rotarod. Locomotor coordination and balance were assessed using an accelerating rotarod (Ugo Basile Original RotaRod for Mice). One day before testing, mice were habituated and trained at a fixed speed of 5 rpm until they were able to remain on the rotarod for 300 s. Mice unable to reach 300 s after three training trials were excluded. On the test day, mice were subjected to three trials on a rotarod accelerating from 5 to 40 rpm over 300 s, with 15-min inter-trial intervals. The average latency to fall was used as a measure of motor coordination.

Nest test. Nest-building behavior was assessed as a measure of goal-directed and executive function–related spontaneous activity. Adult mice were individually housed in standard home cages containing a thin layer of bedding and a single piece of pressed cotton nestlet. The test was initiated at the beginning of the dark phase (∼18:00), and animals were left undisturbed for 16 hours. At the end of the testing period (∼10:00 the following day), each cage was photographed, and the remaining intact nestlet and the constructed nest were evaluated. Nest-building performance was scored by two experimenters blinded to genotype according to a standard 5-point scale^99^: 0, Nestlet untouched (>90% intact); 1, Nestlet partially shredded (50–90% intact); 2, Nestlet mostly shredded but no identifiable nest (<50% intact, flat); 3, Nestlet shredded and partial nest with defined walls; 4, Nearly perfect nest with a crater-like structure and walls >½ of mouse height; 5, Fully enclosed nest with a complete dome and entrance hole.

### Statistical analysis

All data are presented as mean ± s.e.m. and were analyzed using Prism v10.1.2 (GraphPad Software, San Diego, CA, USA). Data distribution was assessed for normality and homogeneity of variance using the Shapiro–Wilk and Brown–Forsythe tests; no transformations were applied. For comparisons between two groups, Two-tailed unpaired t-test, Unpaired t test with Welch’s correction or Mann–Whitney U tests (for nonparametric data) were applied. For comparisons among three or more groups, Ordinary/RM one-way ANOVA, two-way RM ANOVA, or Friedman test (for nonparametric data) were performed, followed by Sidak, Dunn or Tukey post hoc tests. All tests were two-tailed. P values < 0.05 were considered statistically significant (*P ≤ 0.05, **P ≤ 0.01, ***P ≤ 0.001, ****P ≤ 0.0001; for simplicity, ****P ≤ 0.0001 are denoted as *** in figures). A summary of statistical analyses is provided in **Supplementary Table 11**.

## Supplementary Tables

**Supplementary Table 1 |** Human brain sample information

**Supplementary Table 2 |** SCENIC inferred core regulators and downstream interactions in the human mid-fetal dlPFC regulatory network

**Supplementary Table 3 |** Integrated proximal and distal annotations of human mid-fetal dlPFC RARs CUT&Tag

**Supplementary Table 4 |** Human mid-fetal PFC-enriched genes defined from BrainSpan transcriptomic database

**Supplementary Table 5 |** Integrated datasets for construction of the RA-associated gene regulatory network in human mid-fetal dlPFC

**Supplementary Table 6 |** Gene modules of RA-associated genes across human brain development

**Supplementary Table 7 |** GSEA analysis of RA gene modules in relation to human psychiatric disorder database

**Supplementary Table 8 |** MAGMA analysis of RA gene modules in relation to human psychiatric disorder GWAS

**Supplementary Table 9 |** Differentially expressed genes in the mPFC of PD 0 Meis2 cKO mice identified by RNA-seq

**Supplementary Table 10 |** Human mid-fetal dlPFC MEIS2 CUT&Tag peak annotations

**Supplementary Table 11 |** Overview of all statistical tests

